# Age-based approach to characterize the dynamics of cellular processes

**DOI:** 10.1101/2025.09.14.676129

**Authors:** Elad Noor, Kirill Jefimov, Ersilia Bifulco, Evgeny Onischenko

## Abstract

Cells continuously produce and degrade multiple small and large molecules, essential for maintaining homeostasis. The study of these dynamics has gained momentum since the development of pulse-chase and wash-in/out methods, utilizing fluorescent or isotopic labeling of cellular components to assess properties such as turnover rates or half-lives. However, standard analyses of these experiments often depend on simplifications such as the homogeneity of analyzed molecules or their immediate labeling, which do not always hold. Here, we present a rigorous analytical framework that interprets the readouts of dynamic labeling experiments as the distribution of metabolic ages, defined as the time that molecules have spent within a cell, and show that metabolic ages can be quantified by dynamic labeling with minimal assumptions. Using age-based interpretation, we demonstrate how the experimentally observed labeling dynamics is connected to a variety of dynamic parameters including half-lives, decay rates, and residence times and how these interpretations are affected by the conditions of delayed input or complex degradation patterns. To aid in the experimental quantification of dynamic parameters, we introduce a compartmental model framework including an open-source software package. We illustrate the framework’s practical utility by quantifying dynamic parameters and determining the kinetic pool structure of budding yeast proteins at optimal and suboptimal growth temperatures.

**Significance Statement:** To be functional, cells must balance the production and degradation of biological molecules. This is often studied by labeling newly-made molecules with isotopic or fluorescent labels. However, determining parameters of these processes, such as degradation rates, is not easy and is challenged by non-instantaneous labeling and complex degradation patterns. We describe a generic framework for interpreting the results of dynamic labeling experiments based on the concept of metabolic age, defined as the time since a molecule entered the metabolic system. By analyzing the effects of heat stress on protein stability in yeast, we illustrate how this framework and its implementation in a custom opensource package enable us to standardize the determination of various dynamic parameters of metabolism.

## 1 Introduction

Living cells are incredibly complex dynamic systems with thousands of different types of molecules that are constantly produced and degraded [38]. The stability of RNAs and proteins determines their abundance and quality, which is important to maintain cellular hemostasis [5, 46]. Cells also employ complex regulatory mechanisms for dynamic control, such as chemical modifications or subcellular localization [37, 45, 48]. However, it is no simple task to characterize these effects. One common approach is based on dynamic labeling under steady-state conditions (that is, constant cellular growth conditions) when the content of labeled versions of nutrients incorporated into target molecules is measured over time after the label was added externally [5, 12–14, 39, 40]. A canonical example is the isotopic labeling of amino acids that can be used to determine the lifespan and turnover rates of cellular proteins by measuring the dynamics of their incorporation [24]. Similarly, ^13^C-glucose is used to investigate phosphosugar exchange in central carbon metabolism [21, 33]. Dynamic labeling can also be implemented with florescence-based approaches [43, 48, 50].

Common to all these strategies is the way dynamic parameters such as lifespan, turnover, or half-life are determined – typically by assuming that the analyzed molecules can be represented by a single well-mixed pool which starts to become labeled immediately [36]. However, these conditions are not always met. For example, non-exponential degradation patterns have previously been reported for RNA and proteins [5, 29], while a significant delay in labeling by amino acids and other labeled metabolites has been described in unicellular and multicellular organisms [17, 20, 26, 35]. In addition, in many cases the analyzed metabolic system does not retain constant size, as, for example, growing cell cultures. How these factors influence dynamic labeling and how they can be accounted for has yet to be rigorously addressed.

In other fields that also study complex dynamic systems, the concept of *age distribution* was introduced to describe a variety of dynamic characteristics. In geophysics, for example, it is often applied to model the exchange of carbon and water in natural reservoirs, where age is defined as the time that has passed since the atoms entered the dynamic system (e.g., water in the form of rain or atmospheric CO_2_ converted to sugars via photosynthesis) [4, 15, 16, 41]. Similar approaches have been developed for the study of demographics, and pharmacokinetics [9, 31, 44]. The probability distribution of the ages allows us to describe the dynamic properties of such systems in the most general terms and with minimal assumptions [4].

Here, we adapt the concept of age distribution to probe the dynamics of biological systems using dynamic labeling approaches. We show that the results of wash-out labeling experiments in steady-state systems reflect the age distribution of the observed molecular pool, namely the *metabolic age distribution*. We demonstrate how this interpretation allows one to connect the experimental readouts with a variety of dynamic parameters, to account for confounding factors in dynamic labeling experiments, and to highlight their limitations.

As a starting point, in Sections 2.1-2.2 we define the metabolic age distribution and demonstrate its equivalence to the observed steady-state labeling dynamics. In Sections 2.3-2.5 we use this equivalence to connect the observed labeling dynamics with various parameters including the residence time, decay rate, or age of the observed molecules under conditions of delayed input, complex degradation patterns, and cell proliferation. To facilitate the quantification of these parameters in real experiments, in Sections 2.6-2.7, we introduce compartmental models (CM) that subdivide a metabolic system into distinct parts that represent physical compartments or biochemical reaction pools. Finally, in Sections 2.8 and S1.8.1, we illustrate the practical application of the age-based interpretation and CM representations by quantifying the dynamic parameters of the yeast proteome at optimal and suboptimal growth temperatures and the assembly order of the nuclear pore complex.

## 2 Results

### 2.1 Steady-state labeling dynamics: conventions and definitions

To mathematically describe dynamic labeling experiments, we will use strict definitions and assumptions about the properties of biological systems (see Appendix S1.1). Here, we refer to biological systems that are analyzed using dynamic labeling – for example, cell cultures – as *metabolic systems*. Specific sets of molecules within the metabolic system are called *metabolic pools*, for example, vacuolar and protein-borne amino acid pools. The pools that are experimentally analyzed are called *observed*, and they are *unobserved* otherwise, and together comprise a *whole* metabolic system. We consider that the metabolic systems are in a dynamic steadystate during the experiments, i.e. that all of their biochemical parameters, including the relative sizes of metabolic pools or the rates of biochemical reactions, remain constant. These conditions can be met in nondividing cells, in exponentially growing cell cultures that remain in the same culturing conditions, or when cells are examined over sufficiently short time intervals where their metabolism remains essentially unchanged.

We strictly define the dynamic properties of metabolic systems. The term *metabolic age* (𝒜 ) defines the time that has passed since the metabolite particles entered the metabolic system, while *residence time* (𝒯) is defined as the time spent within the whole system or its observed pools between the moment of entry and the moment of exit (Figure 1A). Note that the term *transit time* is sometimes used in the literature to convey the same meaning [4, 15, 16]. We can also define the total time that currently observed metabolite particles spend in the system, *residence time of the living (RTL)*, which is analogous to a survival characteristic *lifespan of the living* used in demographic studies [44]. This period can also be divided into metabolic age and *the remaining residence time of the living (RRTL)* (Figure 1A). Lastly, the term *decay rate* (*κ*) defines the fractional escape rate of metabolite particles from the metabolic system or its parts.

**Figure 1:**
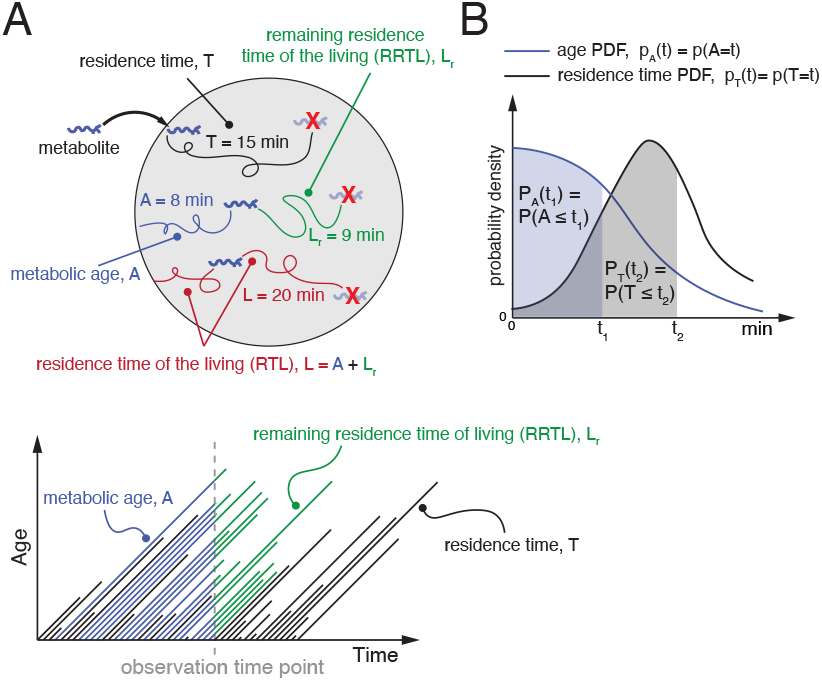
Definition of dynamic parameters. (A) top: Schematic of a metabolic system (e.g. a cell culture) fed by an external metabolite (nutrient). The time elapsed since the metabolite entry defines its metabolic age 𝒜. The period between entry and escape from the system defines residence time 𝒯. The total time the currently present metabolite particles spend in the system defines residence time of the living ℒ, while remaining residence time of the living ℒ_*r*_ defines how long these molecules will remain in the system from the moment of observation. bottom: Age/time diagram illustrating relationships between dynamic parameters. The diagonal tracks represent “life tracks” of metabolite particles starting from the “moment of birth” (entry) and ending at the “moment of death” (escape form the system). Dashed - the moment of observation. (B) Example of steady-state distributions (PDFs) of metabolic ages and residence times *p*_𝒜_ and *p*_𝒯_ . Shaded areas illustrate the corresponding cumulative probabilities (CDFs) *P*_𝒜_ and *P*_𝒯_ for times *t*_1_ and *t*_2_ respectively.

We consider metabolic ages and residence times to be random variables described by constant probability density functions (PDF), *p*_𝒜_ (*t*) and *p*_𝒯_ (*t*), as shown in Figure 1B. We can also define them through the cumulative distribution functions (CDF), that is, the fraction of particles younger than *t* among all particles within the system – *P*_𝒜_ (*t*) and the fraction of particles spending less than time *t* in the system among all particles leaving the system – *P*_𝒯_ (*t*). As is true for any probability distribution, the CDF and PDF are related through differentiation, e.g.:

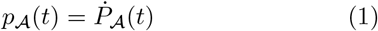

where the dot above a function symbol represents its time derivative, i.e. *ġ* = *dg/dt*. Note that we use the variable *t* to represent both age and time, although they are not exactly the same. However, this duality has a reason that will become clear in the following section.

Throughout this work, we focus on a specific “wash-out” labeling scenario in which the metabolic system is initially fully labeled by an externally supplied metabolite and at time *t* = 0 the external supply is immediately switched to the unlabeled variant without disrupting steady-state cell metabolism (Figure 2). The metabolic label can represent an entire molecule, a chemical group, or even a single atom, as long as it can be experimentally detected and quantitatively compared to the unlabeled variant. Crucially, the two variants must be metabolically indistinguishable (or the effect of swapping them is negligible). The readout of such experiments is the *labeling dynamics(curve)* – denoted by *f* (*t*) – defined as the fraction of the label that remains in the system at times *t >* 0. For example, the time at which *f* (*t*) = 1*/*2 is defined as the *half-life* (denoted *t*_½_ ), i.e. the time it takes to wash out exactly half of the labeled metabolite molecules.

**Figure 2:**
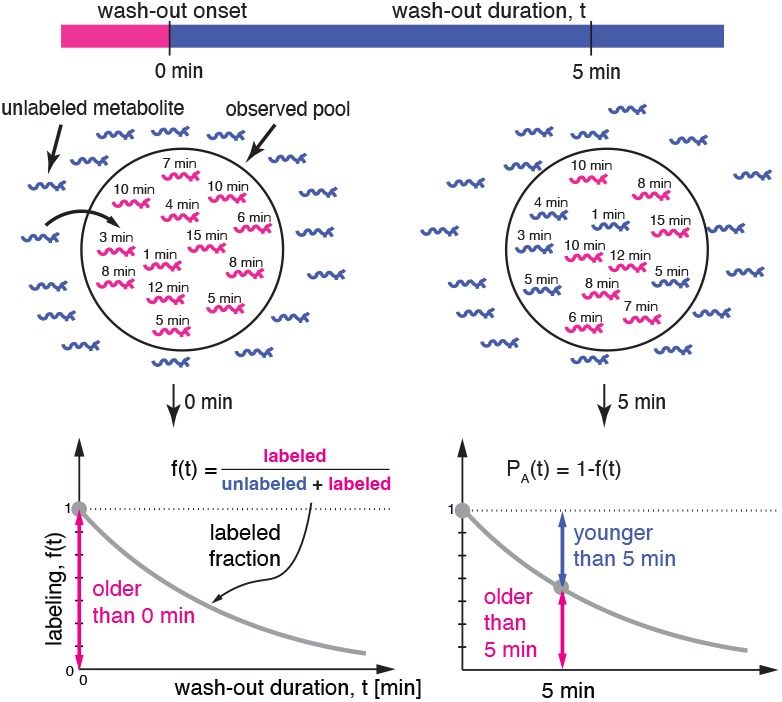
Equivalence of metabolic age distributions and labeling dynamics. Label type (blue or pink) reflects age of metabolite particles in the metabolic system. Graphs below show the dynamics of labeled fraction *f* (*t*), where 1 − *f* (*t*) is equivalent to the metabolic age CDF.

A common example of this scenario is when stable isotopes are used as labels and their relative abundance is determined by quantitative mass spectrometry. Although here we focus on these types of experiments because they correspond to the simplest notation, one can easily adapt all relationships derived on this basis also for “wash-in” and “pulse-chase” labeling, as well as other dynamic labeling techniques, including radioisotopes and genetically encoded fluorophores, which we address in Appendix S1.2.

### 2.2 Steady-state labeling dynamics defines metabolic age distribution

We can now demonstrate that, in steady-state dynamic labeling experiments, the labeling dynamics and the metabolic age distribution are unequivocally connected. Consider *t* = 0 to be the time at which the wash-out starts. Since all molecules start as labeled – if we encounter after some time *t >* 0 an unlabeled one, then it must have entered the system after the wash-out began and therefore is younger than *t*. On the flip side, any labeled molecule is necessarily older than *t* (Figure 2). In other words, the fraction of labeled/unlabeled metabolites and the fraction of its old/young instances are, in fact, two sides of the same coin. As this relationship is valid for any time *t >* 0, we can conclude that the unlabeled fraction of the metabolite is equal to the CDF of its *metabolic age P*_𝒜_ (*t*), that is, the probability that the time it spent in the metabolic system is shorter or equal to *t* (Figure 2B). Since labeled and unlabeled fractions add up to 1, we can now write:

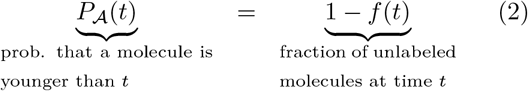

The PDF of the metabolic age can be found using the relation to the CDF in Equation 1, showing that it is defined by the slope of the labeling curve:

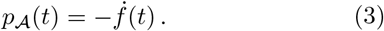

As a consequence, since *p*_𝒜_ (*t*) takes only nonnegative values, steady-state labeling dynamics *f* (*t*) must be a (weakly) monotonically decreasing function. Therefore, if *f* (*t*) increases at some point, the system must be out of steady-state (Figure 3A).

**Figure 3:**
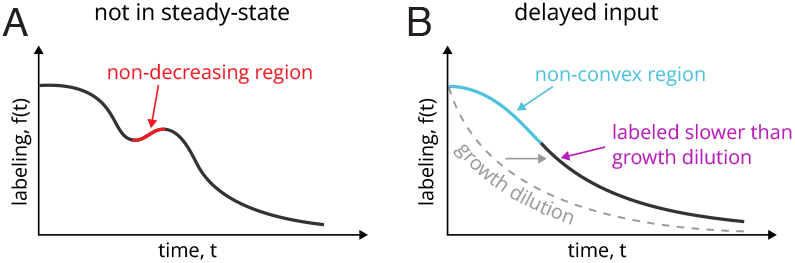
Criteria of non-steady-state dynamics and delayed input. (A) Non-monotonous labeling dynamics indicates that the system is not in steady-state. (B) When the labeling happens more slowly than the dilution defined by growth and/or has an inflection point (i.e. a non-convex labeling curve) then the input must be delayed.

Using the age-based interpretation and Equation 2, we can further see that the half-life *t*_½_ can be mapped to the median metabolic age – *P*_𝒜_ (*t*_½_ ) = 1*/*2. This provides a new interpretation for this metric commonly used in dynamic labeling experiments [17, 36] (Figure 4). We can also see that the mean metabolic age turns out to be relatively simple to calculate as well and is equal to the area under the labeling curve (Appendix S1.3.1):

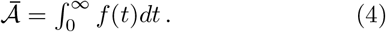

**Figure 4:**
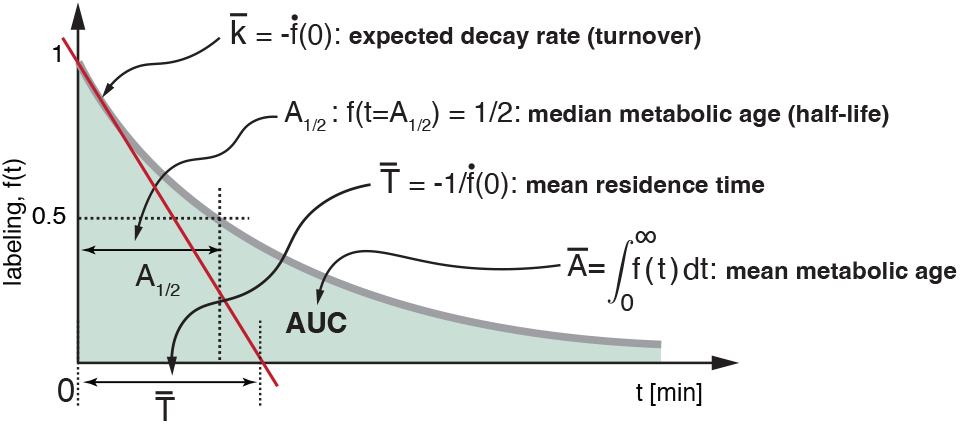
Relation of some cumulative dynamic parameters to the geometry of the labeling curve in nongrowing systems. The half-life, *t*_½_, defines median metabolic age of the measured metabolite. The mean metabolic age, 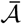, corresponds to the area under the labeling curve (AUC). The expected decay rates, 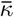, corresponds to the magnitude of the curve’s slope at time 0 or its tangent at the initial point (red line). The mean residence time, 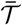, corresponds to the point where this tangent line intersects with the time axes.

Metabolic ages can also be characterized conditionally, focusing on a specific range (Appendix S1.3.6).

Since the median and mean values of metabolic age depend only on the shape of the labeling curve, they can be determined directly from the labeling data (Figure 4). For example, the mean age 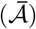 can be estimated geometrically by dividing the area under the curve into trapezoids and calculating their total area (Appendix S1.10.2).

In conclusion, the metabolic age distribution is unequivocally related to steady-state labeling dynamics, which makes it quantifiable without prior assumptions.

### 2.3 Other dynamic parameters are directly related to labeling dynamics when input is not delayed

Using the above connections, we can next show that labeling defines other dynamic parameters when the time it takes to enter an observed pool from the external environment can be neglected. For example, this is the case for soluble lysine molecules entering the cell (Figure 9B), in contrast to protein-borne lysine molecules which are detectable only after some delay. In Section 2.6, we show how one can deal with delayed inputs.

To envision these connections, we avoid specific assumptions about any particularities of internal metabolism by thinking of the metabolic system as a “black box”, which allows metabolite particles to enter and leave the system (Figure 5).

**Figure 5:**
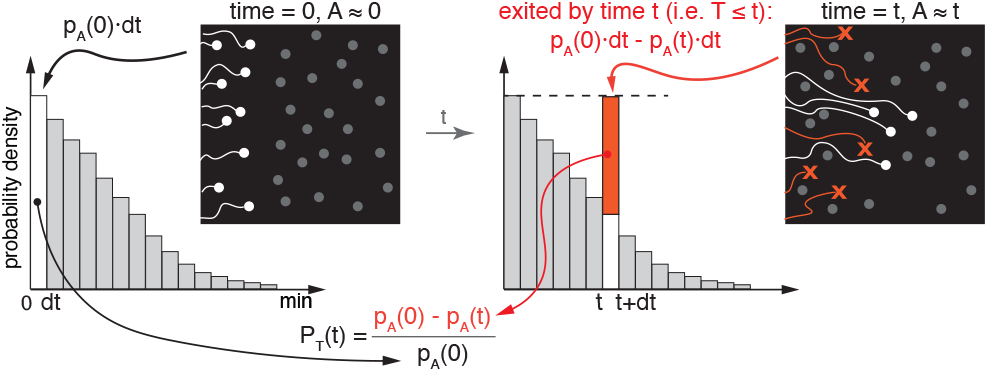
Metabolic age and residence time distributions are directly related in steady-state metabolic systems. “Black-box” representation of a steady-state metabolic system that allows particles (white) to enter and leave. Histogram illustrates the distribution of metabolic ages and its relationship to the CDF of residence times.

Since in the absence of delay, all particles entering the system also have an age 𝒜= 0 at the time of entry, we can focus on a small batch of “newborn” particles that just entered (within a short time interval *dt*) and quantify the initial relative size of this batch using the PDF of age as *p*_𝒜_ (0)*dt*. At some later time point *t* the age of batch members remaining in the system would be equal to *t* while its relative size would decrease, as some particles have left the system, and correspond to *p*_𝒜_ (*t*)*dt* (Figure 5). Importantly, the missing particles were those that left the system before the time point *t*. In other words, the decrease in batch size, *p*_𝒜_ (0)*dt* − *p*_𝒜_ (*t*)*dt*, represents those members that had a residence time shorter than *t*. This allows us to determine the residence time CDF, *P*_*𝒯*_ (*t*), as the fraction of the initial batch that left the system using the age PDF, as shown in Figure 5:

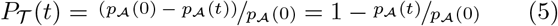

Taking the derivative (similar to Equation 1) and replacing the age PDF with the labeling dynamics (based on Equation 3), we can get:

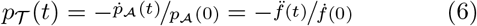

Therefore, the residence time PDF, similar to age, can be directly inferred from the labeling dynamics. We can also analogously draw the connections to the related parameters, residence time of the living ℒ and remaining residence time of the living ℒ_*r*_ (Figure 1A). As shown in Appendix S1.3.4, their PDFs are given by 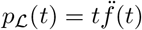 and 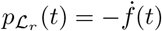.

Notably, Equation 6 also defines a criterion for nondelayed input. Since, by definition, *p*_𝒯_ (*t*) *>* 0 and 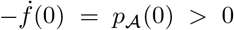 (based on Equation 3), we can conclude that 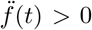, which means that the labeling curve must have a convex shape. Therefore, observing a confident increase in the magnitude of the slope of the labeling curve (i.e., a non-convex segment) would indicate a delayed input (Figure 3B). Since direct quantification of curve slopes is sensitive to measurement or biological noise, this criterion is applicable only if these contributions can be safely excluded.

Next, we can see that decay rates, i.e. the fractional rates at which particles leave the system, generally depend on their metabolic age. We define the age-dependent decay rate *κ*(𝒜) as the rate at which particles of metabolic age 𝒜 leave the system. Mathematically, we can express this as 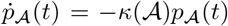, and combining this with Equation 3 we get:

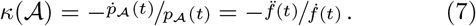

One can see that the middle term is equal to the time derivative of − ln(*p*_𝒜_ (*t*)), and therefore:

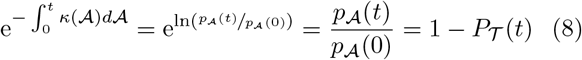

where we apply Equation 5 in the last step.

In general, the decay rates cannot be defined by a single value (see example in Figure S7A) and, similarly to residence times, depend on the first- and second time derivatives of the labeling curve. The only exception is when labeling dynamics is characterized by a single-exponential decay *f* (*t*) = e^−*λt*^, in which case *κ* = *λ*.

Using these generalized relations, one can determine cumulative characteristics for dynamic parameters such as mean or expected values (Figure 4, see Appendix S1.3.2-S1.3.5):

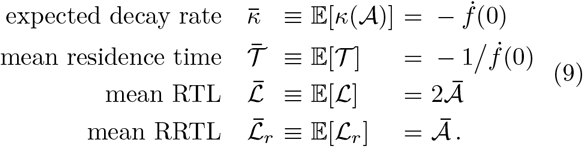

An important implication of these relations is that the mean values of the decay rate or residence time – commonly used stability characteristics – are not related to the overall shape of the labeling curve (Appendix S1.7.1 and Figure S10A) but depend solely on the initial slope 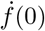. Therefore, their experimental determination requires highly accurate measurements of labeling dynamics at very short time intervals immediately after washout onset (Appendix S1.10.3), which tend to be noisy. To alleviate these difficulties, it is possible to characterize these parameters in an age-conditioned manner skipping the very short time-scales. *Age-conditioned expected decay rate*, 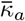 – the expectancy of the decay rate after a certain age – and the *age-cohort mean residence time* 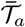 – the mean residence time among all particles that reached a certain age (see Appendix S1.3.6) can be calculated as:

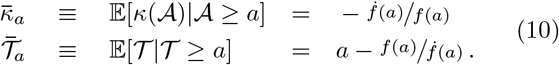

Since 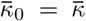 and 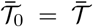, one can view these age-conditioned dynamic parameters as generalizations of Equation 9.

### 2.4 Exponential growth affects the connection between labeling dynamics and dynamic parameters

Up to this point, we assumed that metabolic systems have a constant size albeit many experiments are done in growing cell cultures, i.e., when the biomass (or volume) of the whole system increases over time. If the composition of the culture remains uniform and the culture grows exponentially, all its different pools also grow exponentially at exactly the same rate, a condition called *balanced growth*. In this case, for every unit volume within the system, the pool composition and the flux magnitudes remain constant, and we can treat such constant volumes mathematically as if they were in steadystate.

Since the labeling and age distributions are both defined as fractions relative to all particles in the system, the relationship between these two functions will remain unchanged and follow Equations 1-4. However, there are additional constraints on labeling and mean age imposed by the growth rate *µ* (as shown in Appendix S1.4.4):

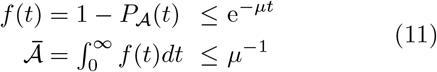

Unlike in the case of the age distribution, the relationships between residence time and decay rate to labeling dynamics are affected by growth. Consider a fixed volume of the growing system (which remains *V*_0_, as in Figure 6). All particles entering this fixed volume at *t* = 0 will eventually escape in one of two ways: by decaying or by moving outside and contributing to the growth of the system. The former is quantified as before by *κ*(𝒜), while the latter (which we call growth dilution) is equal to *µ*, as it must keep the wider system in balanced growth. The two rates are mutually exclusive and, therefore, the *total escape rate λ*(𝒜) is given by their sum. On the flip side, *λ*(𝒜) still follows the relation to the age distribution and labeling dynamics in the fixed-volume system, i.e. as in Equation 7. Therefore, we can write:

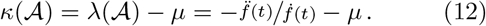

**Figure 6:**
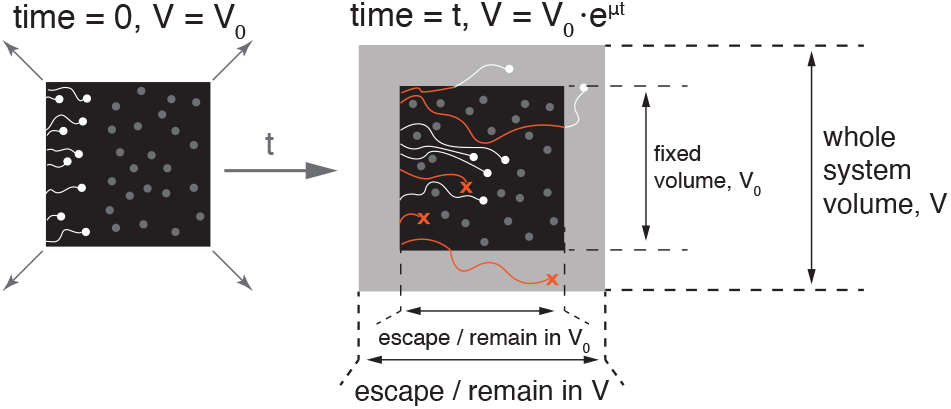
Dynamics of exponentially growing metabolic systems. Time-evolution of an exponentially growing system. After time *t* following the initial time point, the entire system expands from its initial volume *V*_0_ to *V*_0_e^*µt*^. During this time, only some of the unlabeled particles that entered at *t* = 0 would remain in the volume equivalent to *V*_0_ (black box, white dots). The rest would either decay (red crosses) or still exist in the newly created volume (white dots in the gray frame).

Since residence times and decay rates describe the life trajectory of single particles, the mathematical relationship between *P*_𝒯_ (*t*) and *κ*(𝒜) is not affected by growth. Therefore, we can continue using Equation 8 and apply Equation 12 to express the residence time CDF in terms of labeling dynamics:

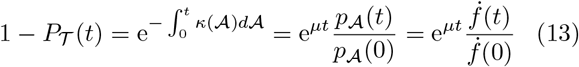

and then get PDF by taking the time derivative:

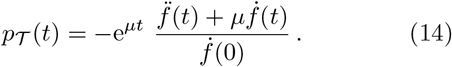

Similar relations can also be derived for the PDF and mean of the RTL and RRTL:

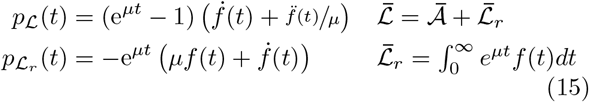

where we can appreciate that it is consistent with our previous results for non-growing systems when *µ* → 0 (see Appendix S1.4 for the derivation details).

In summary, balanced growth affects the relation of observed labeling dynamics with dynamic parameters. Its effect varies, and the correction required can be either none (for the ages), a simple growth rate subtraction (for the decay rates), or a more complicated expression (for the residence times and related parameters). For example, as a consequence of growth, the expected decay rate and the mean residence time are no longer inversely related (see Appendix S1.4 and Figure S7B).

### 2.5 Determination of dynamic parameters requires compensation for labeling delay

When linking between labeling dynamics and dynamic parameters, we assumed that the input is immediate, i.e. that the age of particles entering the observed system is 𝒜 = 0 at the moment of entry. However, this condition is rarely met in practice as the external molecules typically take some time to reach the observed pools. For example, incorporation of amino acids into cellular proteins can be significantly delayed in animal tissues when the label is introduced as a dietary supplement [17, 20]. Delayed input can have a clear signature, such as a flat initial slope 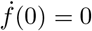 (Appendix S1.5.5), areas of nonconvexity 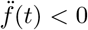, and, in case of growing systems, a wash-out dynamics which is slower than the growth dilution – *f* (*t*) *> e*^−*µt*^ (see Figure 3B, S7D, and Appendix S1.4.4). In special cases, when analyzed particles can be labeled multiple times (e.g. tryptic peptides labeled at a single or multiple positions by amino acid labels), the delay can also be detected by slower initial wash-out for species with higher labeling multiplicity (Appendix S1.5.2).

Although the effect of delay is not always trivial, we can always define a compensation procedure that reduces the entry ages of all input particles to 𝒜 = 0 so that the compensated labeling dynamics can then be used to determine all the dynamic parameters as described in previous sections. This procedure can be exemplified in the case when an observed subsystem is fed by a *single-source* pool with some known age distribution *p*_𝒜°_ (*t*) (Figure 7). This is a reasonable description of cases such as metabolic labeling of RNA or proteins by smallmolecule precursors. The age 𝒜 of each particle in this case represents the sum of the age at which it entered the observed system, 𝒜°, and the time it spent in it from the moment of entry, which we denote by 𝒜_*r*_. The 𝒜_*r*_ is equivalent to the age of the particles in a *reduced* system into which they enter without delay (Figure 7).

**Figure 7:**
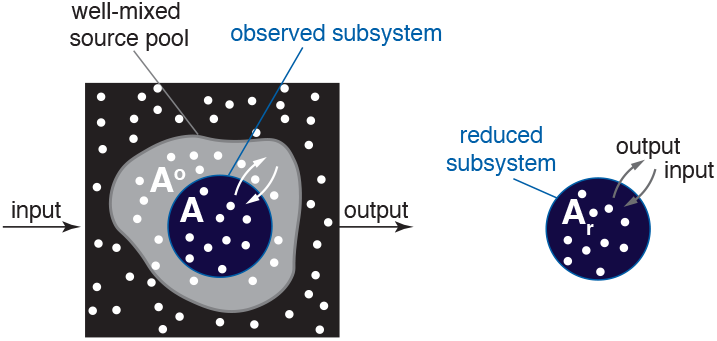
Determining dynamic properties of systems with delayed input. Example of a system with delayed input. (left) The observed subsystem (blue) receives particles from unobserved parts which represent a single-source pool with a known metabolic age distribution 𝒜°. (right) Compensating for the input delay is equivalent to resetting the ages of all particles entering the subsystem to 𝒜 = 0, i.e. as if it were fed directly from the external environment. The reduced metabolic ages 𝒜_*r*_ reflect the time spent by the particles in the observed subsystem.

Since 𝒜° and 𝒜_*r*_ are independent variables, the probability distribution of their sum is given by a convolution of the two PDFs:

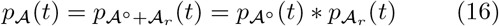

where − stands for convolution, i.e., 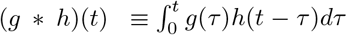. This allows us to connect observed labeling dynamics *f* (*t*) with those in the input *f* °(*t*) and in the reduced system *f*_*r*_(*t*) (Appendix S1.9) as:

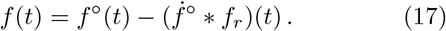

Consequently, determining dynamic parameters in the case of delayed input requires knowing both the observed labeling *f* (*t*) and the labeling dynamics of the input *f* °(*t*). For example, mean age can be determined by subtracting the mean age of the input from the observed mean age (see Appendix S1.9):

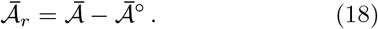

or, equivalently, by calculating the area between the curves of *f* °(*t*) and *f* (*t*) (see Equation 4). More generally, *f*_*r*_(*t*) and then the dynamic parameters can be found by fitting *f*_*r*_(*t*) based on the convolution relation using Equation 17.

In summary, the observed labeling dynamics, even if we know them precisely, is insufficient to determine all the dynamic parameters of an observed system with labeling delay (see Appendix S1.5.3 and Figure S7C for an example). Rather, a compensation procedure is required to virtually reduce the age of the particles upon entry to zero. In the following section, we describe the general implementation of this compensation procedure using compartmental models.

### 2.6 Connecting labeling dynamics with dynamic parameters using compartmental models

The relations between labeling dynamics and the dynamic parameters described so far do not require assumptions about the internal structure of metabolic systems. However, because experimental data are usually discrete and sparse, the practical usage of these relations to quantify dynamic parameters can be complicated, as they have high sensitivity to measurement noise and require a high sampling frequency. As an alternative, one can explicitly represent the internal structure of the metabolic system using a set of pools (compartments) connected by material transfer fluxes, denoted a *compartmental model* or CM in short [1, 22, 23] – and frequently used in pharmacokinetic analysis [9]. The word *compartmental* is used here in a wide context and can represent not only physical compartments but also other types of physical or chemical separation attributes. For example, these can be a set of membrane-delimited organelles connected by transport routes or metabolic states in a pathway connected by enzyme-catalyzed chemical reactions (Figure 8).

**Figure 8:**
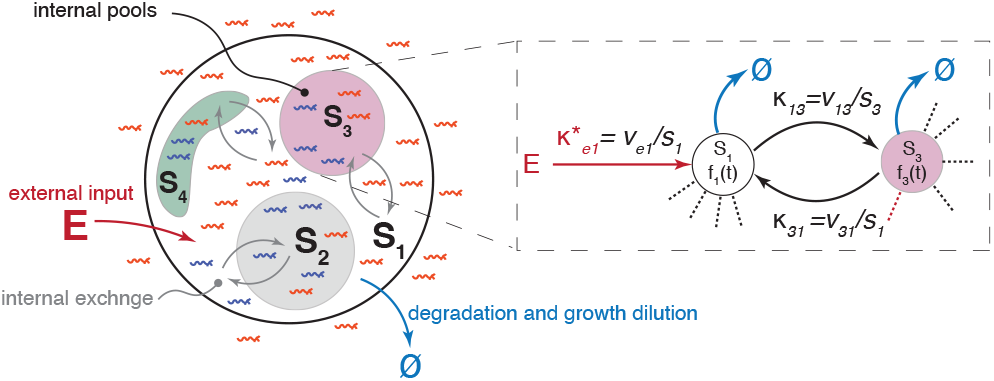
Description of steady-state labeling dynamics using compartmental models. Parametric description of particle propagation through internal metabolic pools via contributed turnovers that represent connecting flux rates from source to recipient pools normalized by the size of the receiving pool.

Mathematically, the CM-based description can be seen as a Markov process in which the possible fates of each particle depend only on its current state and not on its history [32]. For example, in a CM representation of glycolysis, we require that the fate of cytosolic glucose-6-phosphate does not depend on how or when it was imported into the cell. Since this is arguably a fair assumption for any biochemical process, CMs (with an unlimited number of compartments and flux routs) should be able to recapitulate any valid age distribution.

In steady-state systems described by CMs, the number of compartments (pools), their sizes (denoted *s*_*i*_), and the rates of material transfer between them, are all constant. Therefore, we can define the transfer rates by the *contributed turnovers κ*_*ji*_ ≡ *v*_*ji*_*/s*_*i*_ and *κ*_*ei*_ ≡ *v*_*ei*_*/s*_*i*_ that represent influxes from the internal and external pools, respectively, normalized by the size of the receiving pool (Figure 8 and Appendix S1.6).

Defining **f** (*t*) as the vector of labeling dynamics across all states and using the above conventions, one can see that **f** (*t*) relates to its time derivative via a linear relationship:

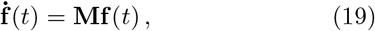

where **M** is referred to as the *state transition matrix* and comprises combinations of contributed turnovers (Figure S5E, see Appendix S1.6.2). Equation 19 is a first-order linear homogeneous ODE (standard for continuous-time Markov processes [32]) with a known analytical solution [6, 47]:

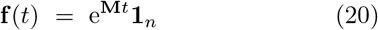

where e^**M***t*^ is a matrix exponential that can be calculated analytically in simple cases or numerically [2] in general. Using this solution, we can define the *system labeled fraction* as:

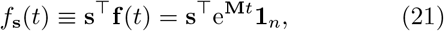

where **s** is the vector of the relative pool sizes (which sums up to 1). The distributions as well as cumulative values of various dynamic parameters can then be found using relations to labeling dynamics described in Sections 2.2 – 2.4 (see Appendix S1.6.5 for the full derivations). The main results are summarized in Table 1.

**Table 1:**
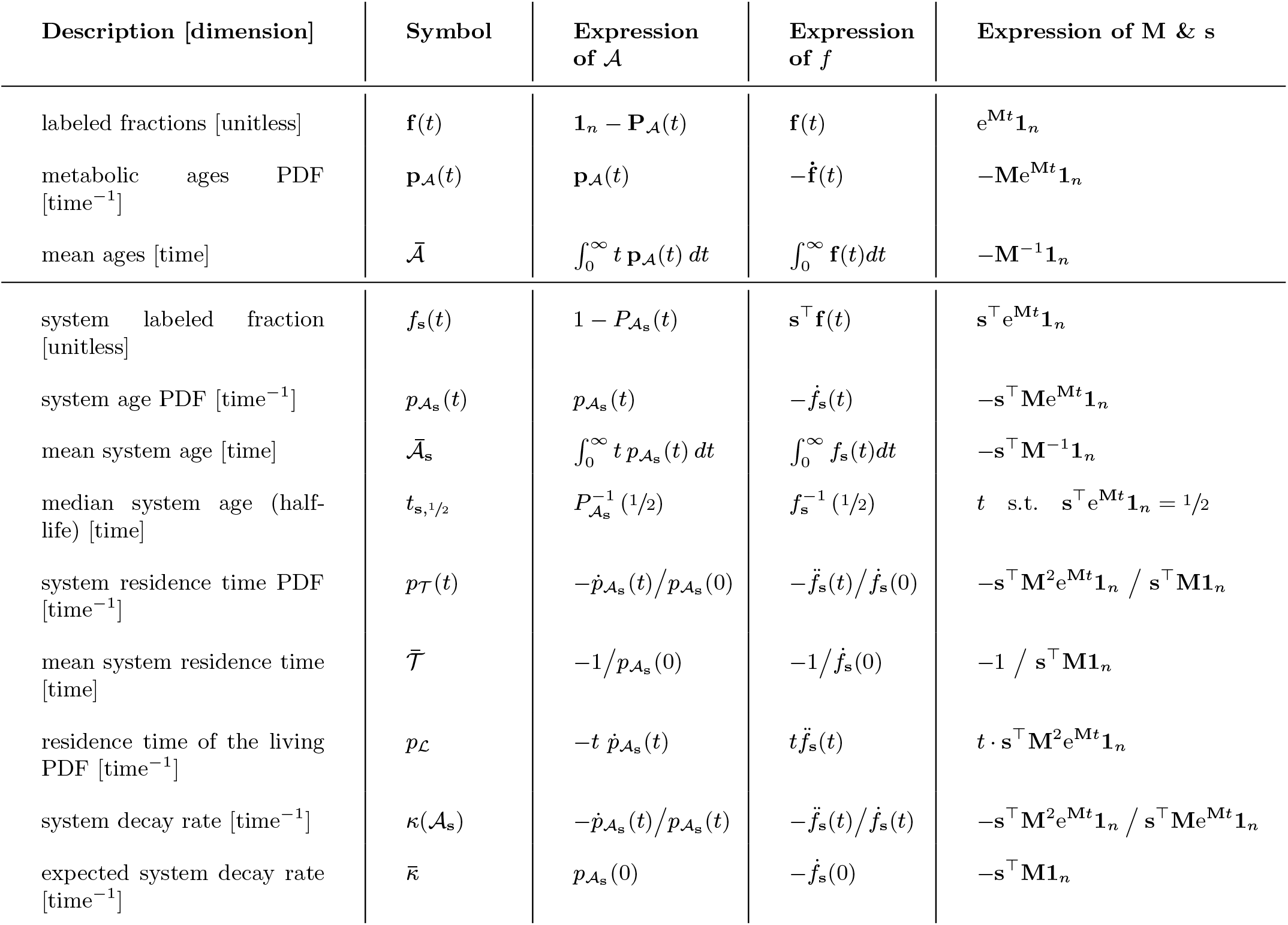
Dynamic parameters of steady-state systems and their relation to the metabolic age, experimental labeling dynamics, and compartmental model parameters. The expressions connecting labeling dynamics, age distributions, and residence times with CM parameters. Some of these can be defined for each individual state, and also for the whole system (i.e. the weighted average across all the pools, in proportion to their sizes). For a more comprehensive summary, see Appendix Table S1.

| Description [dimension] | Symbol | Expression of $\mathcal{A}$ | Expression of $f$ | Expression of M & s |
| --- | --- | --- | --- | --- |
| labeled fractions [unitless] | $\mathbf{f}(t)$ | $\mathbf{1}_n - \mathbf{P}_{\mathcal{A}}(t)$ | $\mathbf{f}(t)$ | $\mathbf{e}^{\mathbf{M}t} \mathbf{1}_n$ |
| metabolic ages PDF [time<sup>-1</sup>] | $\mathbf{p}_{\mathcal{A}}(t)$ | $\mathbf{p}_{\mathcal{A}}(t)$ | $-\dot{\mathbf{f}}(t)$ | $-\mathbf{M} \mathbf{e}^{\mathbf{M}t} \mathbf{1}_n$ |
| mean ages [time] | $\bar{\mathcal{A}}$ | $\int_0^\infty t \mathbf{p}_{\mathcal{A}}(t) dt$ | $\int_0^\infty \mathbf{f}(t) dt$ | $-\mathbf{M}^{-1} \mathbf{1}_n$ |
| system labeled fraction [unitless] | $f_s(t)$ | $1 - P_{\mathcal{A}_s}(t)$ | $\mathbf{s}^\top \mathbf{f}(t)$ | $\mathbf{s}^\top \mathbf{e}^{\mathbf{M}t} \mathbf{1}_n$ |
| system age PDF [time<sup>-1</sup>] | $p_{\mathcal{A}_s}(t)$ | $p_{\mathcal{A}_s}(t)$ | $-\dot{f}_s(t)$ | $-\mathbf{s}^\top \mathbf{M} \mathbf{e}^{\mathbf{M}t} \mathbf{1}_n$ |
| mean system age [time] | $\bar{\mathcal{A}}_s$ | $\int_0^\infty t p_{\mathcal{A}_s}(t) dt$ | $\int_0^\infty f_s(t) dt$ | $-\mathbf{s}^\top \mathbf{M}^{-1} \mathbf{1}_n$ |
| median system age (half-life) [time] | $t_{s,1/2}$ | $P_{\mathcal{A}_s}^{-1}(1/2)$ | $f_s^{-1}(1/2)$ | $t \text{ s.t. } \mathbf{s}^\top \mathbf{e}^{\mathbf{M}t} \mathbf{1}_n = 1/2$ |
| system residence time PDF [time<sup>-1</sup>] | $p_{\mathcal{T}}(t)$ | $-\dot{p}_{\mathcal{A}_s}(t)/p_{\mathcal{A}_s}(0)$ | $-\ddot{f}_s(t)/\dot{f}_s(0)$ | $-\mathbf{s}^\top \mathbf{M}^2 \mathbf{e}^{\mathbf{M}t} \mathbf{1}_n / \mathbf{s}^\top \mathbf{M} \mathbf{1}_n$ |
| mean system residence time [time] | $\bar{\mathcal{T}}$ | $-1/p_{\mathcal{A}_s}(0)$ | $-1/\dot{f}_s(0)$ | $-1 / \mathbf{s}^\top \mathbf{M} \mathbf{1}_n$ |
| residence time of the living PDF [time<sup>-1</sup>] | $p_{\mathcal{L}}$ | $-t \dot{p}_{\mathcal{A}_s}(t)$ | $t \ddot{f}_s(t)$ | $t \cdot \mathbf{s}^\top \mathbf{M}^2 \mathbf{e}^{\mathbf{M}t} \mathbf{1}_n$ |
| system decay rate [time<sup>-1</sup>] | $\kappa(\mathcal{A}_s)$ | $-\dot{p}_{\mathcal{A}_s}(t)/p_{\mathcal{A}_s}(t)$ | $-\ddot{f}_s(t)/\dot{f}_s(t)$ | $-\mathbf{s}^\top \mathbf{M}^2 \mathbf{e}^{\mathbf{M}t} \mathbf{1}_n / \mathbf{s}^\top \mathbf{M} \mathbf{e}^{\mathbf{M}t} \mathbf{1}_n$ |
| expected system decay rate [time<sup>-1</sup>] | $\bar{\kappa}$ | $p_{\mathcal{A}_s}(0)$ | $-\dot{f}_s(0)$ | $-\mathbf{s}^\top \mathbf{M} \mathbf{1}_n$ |

These relationships can also be demonstrated directly (Appendix S1.6.1). Defining **p**_𝒜_ (*t*) as the vector of all age PDFs across all states, we can derive an ODE in a similar fashion to the labeling dynamics one (Equation 19):

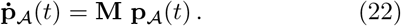

This directly provides a general analytical solution for the metabolic age distributions for every pool of the metabolic system:

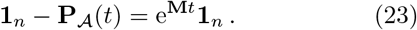

Since CMs can arguably recapitulate any valid dynamics of a metabolic system, this approach can be used to extract dynamic parameters, e.g. those in Table 1, by realistically approximating continuous labeling dynamics based on otherwise discrete and noisy experimental data, or to test hypotheses about the structure of metabolic systems by comparing the explanatory power of alternative CM representations. In particular, CMs provide a powerful way to compensate for delayed input (see Section 2.5). If the whole system, including both the observed and unobserved parts, is represented by a CM, the observed part whose labeling is delayed can always be reduced to the equivalent directly-labeled system by removing all rows and columns corresponding to the unobserved pools from the **M** matrix and renormalizing the remaining observed pool weights (see Appendix S1.6.3 for further details). The reduced CM denoted by **M**_*r*_ and **s**_*r*_ can then be used directly to determine the dynamic parameters of the observed system as if it were labeled without delay, which we illustrate in Section 2.8. In all these cases **M** and **s** are found by fitting the experimental values **f** (*t*_*i*_) measured at multiple time points – {*t*_*i*_} – with, for example, a non-linear least-squares solver [9] (Appendix S1.10.1).

### 2.7 Properties of labeling dynamics related to compartmental models

The CM representation defines what labeling curves can and cannot look like. One general property, which is a consequence of the general solution for the labeling dynamics from Equation 19, is that any real labeling dynamics can be described as the sum of exponents, namely 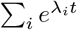 where *λ*_*i*_ are the eigenvalues of **M**, all of which have a negative real part. In turn, **M** must satisfy the requirement of mass balance in which every part of the metabolic system receives as much material as it lets out. This *mass balance constraint* can be formulated as:

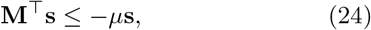

where *µ* is the growth rate in case the system is growing exponentially or equal to 0 for non-growing systems (Appendix S1.6.2). One can show (see Lemma S1.19) that given **M** and *µ*, there exists a pool size vector **s** that satisfies the mass balance constraints if and only if all real parts of the eigenvalues of **M** are smaller than −*µ*. As a consequence, for sufficiently long timescales, the values of **f** (*t*) always decrease exponentially and faster than the growth rate (Appendix S1.6.4). This makes intuitive sense, since if one or more eigenvalues were larger (less negative), the solution would include an exponent that decays more slowly than *µ*, and would therefore be inconsistent with balanced growth.

Using the CM representation, one can also show that in a metabolic chain (a set of consecutive interconversions of metabolites), the upstream metabolic pools are always younger and become unlabeled faster than the downstream ones, that is, at any time *t* ≥ 0:

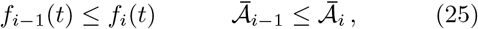

where *i* is the sequential pool index (see Appendix S1.7.4). This property enables the use of dynamic labeling as a simple way to determine the order of events in metabolic chains. As a practical example, using this approach we reanalyze the assembly sequence of the nuclear pore complex based on dynamic labeling data from [35] and show that it recapitulates previous findings with fewer assumptions and in a more rigorous manner (see Appendix S1.8.1).

### 2.8 Example: quantifying dynamic parameters of yeast proteome in dynamic SILAC experiments

To illustrate the utility of the age-based framework, we applied it to determine dynamic parameters of the budding yeast proteome at 30^°^C and at 37^°^C – corresponding to mild heat stress conditions – using dynamic SILAC (stable isotope labeling with amino acids in cell culture) [13]. This method has been widely used to explore the dynamics of the yeast proteome [8, 28] that focused on the half-life of protein labeling, i.e., the median metabolic age of the amino acid label in the protein pools [36, 40]. However, the effects of delayed input, complex degradation patterns, and exponential growth have not been explored in detail. Using our framework, we sought to bridge these gaps.

For dynamic labeling, we grew yeast cells deficient in lysine biosynthesis in a light-lysine medium and quickly switched to an equivalent medium with heavy lysine to initiate wash-out. Following the switch, the collected cultures were analyzed to determine protein labeling dynamics, i.e., the fraction of light lysine in the protein pools, and to record biomass growth (Figure 9A). At both temperatures, the cultures maintained exponential growth in-line with balanced growth conditions (Figures S11A) and at 37^°^C also showed an elevated abundance of heat shock proteins, which confirmed adaptation to heat stress (Figure S11B).

**Figure 9:**
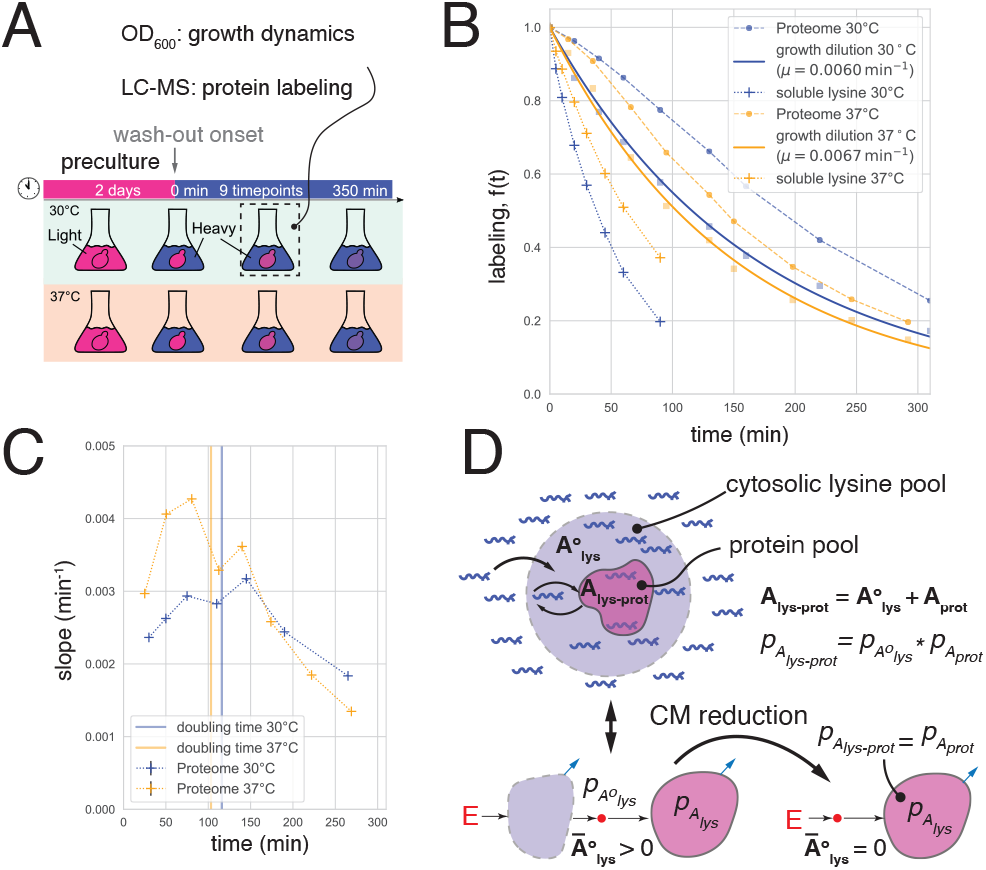
Lysine input is significantly delayed. (A) Outline of dynamic SILAC assay. Yeast cells are precultured in light lysine medium at 30^°^C or 37^°^C and light lysine is washed out by switching to heavy lysine medium. Culture samples are drafted to determine the culture growth dynamics and protein labeling.(B) Labeling dynamics of all proteins combined (dashed, circles) quantified at 30^°^C (blue) and 37^°^C (gold) as compared to the growth dilution of the same culture (solid) and to directly measured labeling dynamics of soluble lysine (dashed, crosses). (C) Magnitude of the labeling curve slope calculated between neighboring time points. Vertical lines depict culture doubling times at 30^°^C (blue) and 37^°^C (gold). Note that the slope magnitude increases at the initial time points indicating nonconvexity. (D) top: “Single-source” delayed input scenario: metabolic age of lysine in the observed protein pool 𝒜_lys-prot_ is defined by the sum of its age in the cytosolic pool when used for protein biosynthesis 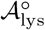 and the age of the protein itself 𝒜_prot_ and correspond to a convolution relationship between the respective age distributions 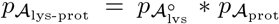 . bottom: CM representation of the “single-source” scenario: a lysine input subsystem feeds lysine with the age distribution 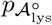 to the proteins’ subsystem. The protein ages can be found by reducing the CM to the protein subsystem (see Appendix S1.6.3).

We first tested whether the delay in the lysine input should be taken into account, using constraints on the labeling dynamics in directly labeled systems as delayed input criteria (Sections 2.3 - 2.4, Figure 3B). As a conservative test, we quantified the labeling dynamics of the whole protein-borne lysine pool based on the combined signal of all lysine-containing peptides. In both 30^°^C and 37^°^C, the labeling of this entire pool decreased more slowly than the growth dilution and was concave (had an increasing slope) during the first 1-2 hours – two indications pointing to delayed input (see Figures 9B-C). Observing such a global delay in the labeling of proteins indicates that there must be large pools of lysine preceding translation. In support of this view, yeast contain large and dynamic vacuolar stores of soluble lysine [10, 34] and, as expected, our direct measurements show that soluble lysine labeling is rather slow at both temperatures (Figure 9B).

To account for delayed input, we describe the dynamics of protein labeling in terms of the metabolic age of lysine. Since all protein molecules are produced from the same cytosolic pool of lysine, its age in the protein pool (𝒜_lys-prot_) can be seen as the sum of age when used for translation, denoted by 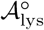 and the age of the protein itself (𝒜_prot_). In this “single-source input” scenario (Figure 9D) the probability distributions of these tree ages and the labeling dynamics corresponding to these age distributions are related by convolution relationships 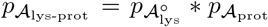 and 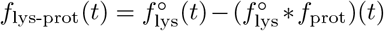 as described in Section 2.5, where 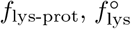 and *f*_prot_ correspond to the observed labeling dynamics, the dynamics of the lysine input, and reduced protein labeling. To compensate for delayed input, we use these relationships in two ways: First, we identify a set of *reference proteins* whose reduced labeling can be determined without knowing the input delay and use their observed labeling dynamics to calculate the delay. Second, we use the calculated delay and observed labeling of each protein to determine the dynamic parameters. To implement these calculations, we describe the single-source input scenario with CMs in which the lysine input is represented by a set of compartments that receive it from the external environment and feed through a single small source pool (representing a mixture of different ages and avoiding additional delays) to the observed protein subsystem (Figure 9D). In this representation, *f*_prot_(*t*), and hence all dynamic parameters of proteins, can be determined by reducing such CMs to the protein subsystem (Appendix S1.6.3).

One example of proteins that can be used as reference are non-degraded proteins. Without input delay, the labeling of non-degraded proteins is fully governed by growth dilution, i.e. *f*_prot_(*t*) = e^−*µt*^, and therefore their mean metabolic age is 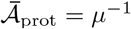 (Equation 11). The growth rate itself can be independently obtained from the known biomass growth dynamics (Figure 9D). Importantly, non-degraded proteins will also be those with the highest mean age of lysine in their pools, since protein degradation can only decrease the mean age. Furthermore, when lysine input is delayed, its mean age 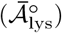 will be a constant amount added uniformly to all proteins: 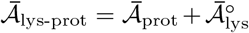 (Equation 18). Therefore, the delay does not change the mean age ranking of the proteins, and the non-degraded ones will remain the oldest. We used this fact to select the reference proteins as the ones with the highest values of 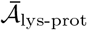 and assuming that they are not degraded. To implement these calculations, we used the simplest delayed-labeling CM where a single lysine input pool feeds a single observed protein pool (Figure 10A).

**Figure 10:**
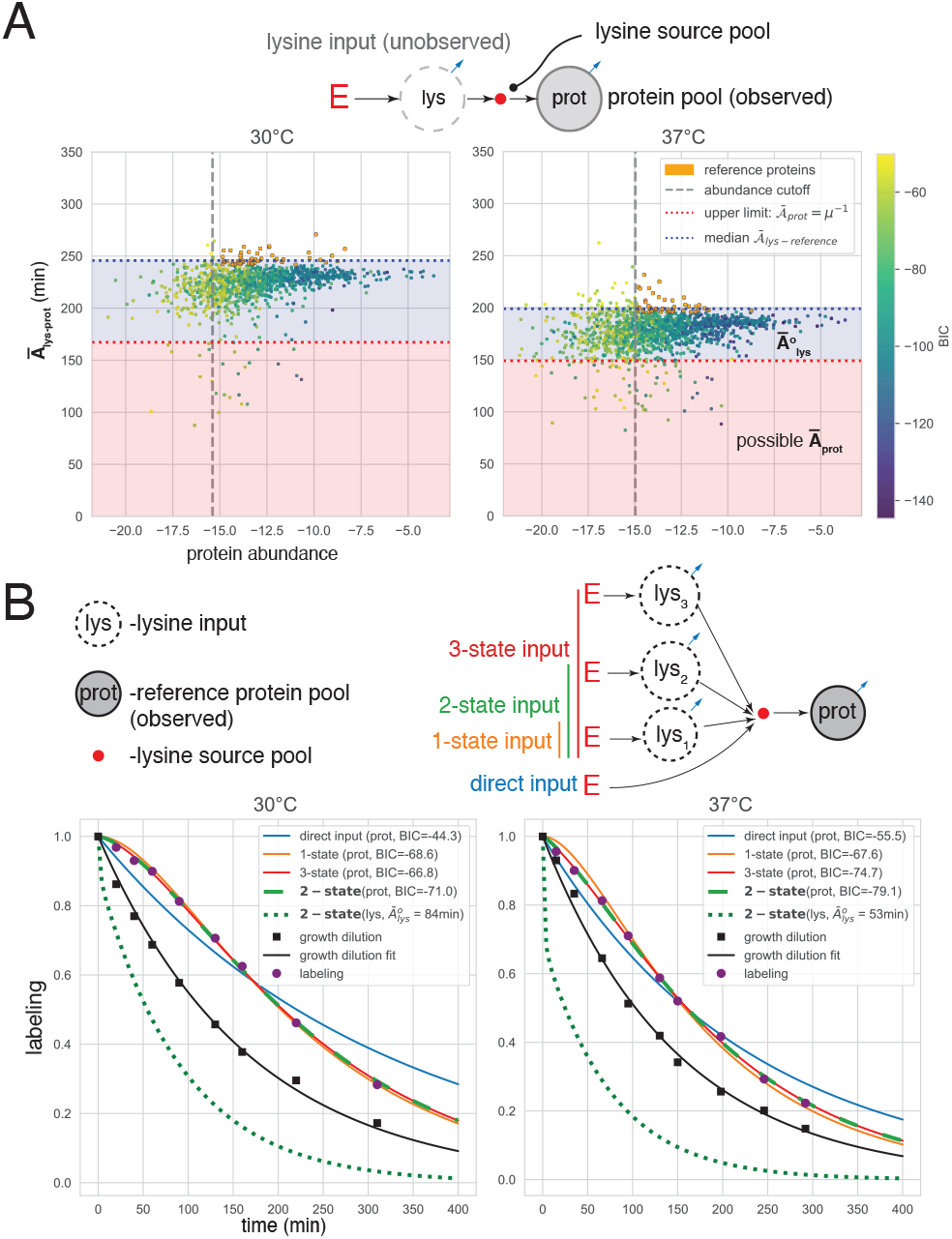
Quantification of lysine input delay. (A) Selection of reference proteins based on lysine metabolic age. Lysine mean age (y-aixs) was quantified by fitting labeling dynamics with a 2-state CM in which the lysine source and the protein pool are each represented by a single compartment and connected by a small source pool. Among the 50% most abundant proteins, we selected the top 5% by lysine mean age to be the reference proteins (gold points). The proteins with poor fits (BIC *>* −50) were excluded from this selection. The red and blue dashed lines represent the theoretical upper bound for the reduced metabolic age of proteins 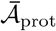 (i.e. assuming a direct input), and the median of mean metabolic ages of lysine in the reference protein pools 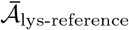, respectively. The fact that 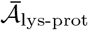 of nearly all proteins exceeds the reduced age limit by up to 50-100 minutes is an indication of delayed input. The mean age of lysine input 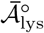 creating this delay can be estimated as the distance between the red and blue lines. (B) top: CMs representations of different lysine input scenarios of reference protein labeling. bottom: CM fits (solid) calculated with the reference protein labeling dynamics (purple dots). Estimated labeling dynamics of lysine input (dotted) calculated from the best CM fit. The input lysine mean age 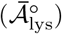 for the best CM and the BIC for all CMs appear in the legend. Growth dilution calculated using whole cell biomass observations is shown in black.

Next, to calculate the delay, we use a CM that represents protein labeling with delayed input, considering that the labeling curve, i.e., the combined labeling dynamics of all reference proteins, is produced by a single non-degraded protein pool fed by some lysine source subsystem (Figure 10B). We considered four possibilities of such input: direct, a 1-state lysine source (the same as the one used for the original ranking), or 2-state and 3-state lysine sources with different turnover, and identified the best configuration based on the BIC values. At both temperatures, the input was best explained by the 2-state option. Notably, the quantified mean age of the lysine input was ≈80 min at 30^°^C and ≈50 min at 37^°^C, that is, each lysine molecule spends roughly an hour inside the cell before being incorporated into a newly-made protein. This timescale is compatible with the observation that the total proteome labeling curve became fully convex only after ≈2 hours, i.e. after the labeled fraction of the lysine input had been nearly depleted (Figure 9C).

Having established the parameters of the input delay, we model the labeling dynamics of all proteins by CMs in which a protein subsystem is fed by the calculated lysine input (Figure 10B). Since proteins can potentially have complex degradation patters, in addition to a single pool model, we also tried to represent them using two or three parallel pools with different decay rates and, once again, selected the best model based on the BIC of the fit. Although most of the labeling dynamics at both temperatures could be described using a single-pool model, there was a small fraction of proteins (≈10% in both temperatures) that required a two-pool model and only few cases required three pools (Figure 11A). Interestingly, structural ribosomal proteins were significantly enriched in the two-pool category (Figures 11B and S12). This suggests that ribosomal proteins have prominent pools with different degradation kinetics, which is in line with the recently described phenomenon of excessively produced and degraded ribosomal protein pools [42].

**Figure 11:**
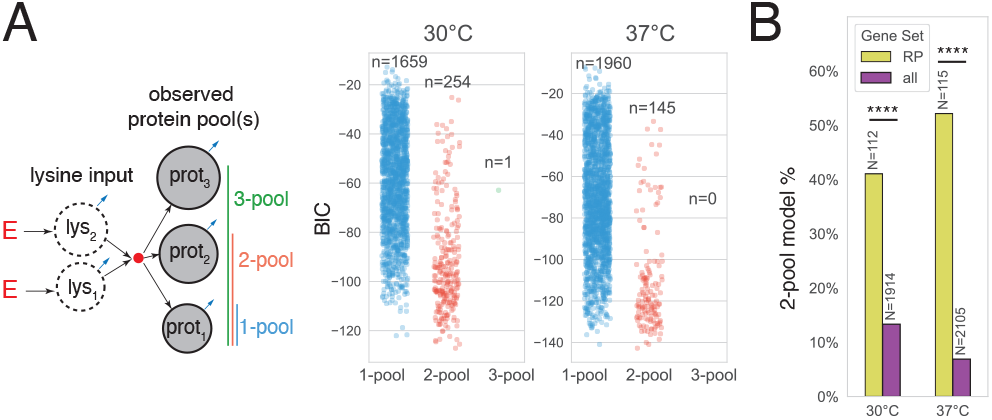
Metabolic pool structure of yeast proteins at 30^°^C and 37^°^C. (A) The labeling dynamics of yeast proteins at 30^°^C and 37^°^C were fitted using 1-, 2-, or 3-pool CMs describing the protein subsystem which is fed by the 2-state input subsystem that models lysine input delay (see Figure 10). Swarm plots depict BIC values corresponding to each model. Note that while most proteins were best explained by the 1-pool model, only one protein required the 3-pool model to describe its labeling dynamics. (B) Unlike most proteins, nearly half of all the ribosomal proteins (denoted RP, GO:0044391) were better described by a 2-pool option, constituting a significant enrichment (****: p *<* 0.0001 by Fisher exact test).

Parameterized CMs allowed us to characterize various dynamic parameters. To control for the robustness of these evaluations, we used bootstrapping and assumed that all proteins can have complex degradation patterns described by a two-pool CM. Treating the bootstrapped data as repeated independent samples, we calculated the coefficient of variation (CV) for each protein, i.e. the standard deviation divided by the mean value. We found that the mean ages of most proteins could be reliably quantified (CV *<* 0.1 for more than 90% proteins), consistent with their dependence on the integrated labeling dynamics (see Equation 4). Interestingly, while there were certain systematic age differences between temperatures (median: 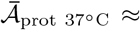 min vs. 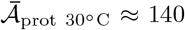 min), they could be eliminated if the mean ages were adjusted for the cell culture growth rate expressing them in units of the maximum mean age – *µ*^−1^ (median 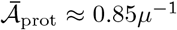 for both temperatures,

Figure 12B) – indicating that the effects of temperature on protein age can be mainly attributed to the duration of the cell cycle. As a benchmark to compare our age predictions with commonly used methods, we also quantified protein ages by fitting the log-transformed labeling dynamics with a linear regression model [40], a method that neglects the possibilities of delayed input and complex degradation patterns. Indeed, following our expectations, this led to an overestimation of mean ages beyond the growth dilution limit for the majority proteins, indicating that an objective description of protein dynamics cannot be made using these simplifications (Figure S13).

**Figure 12:**
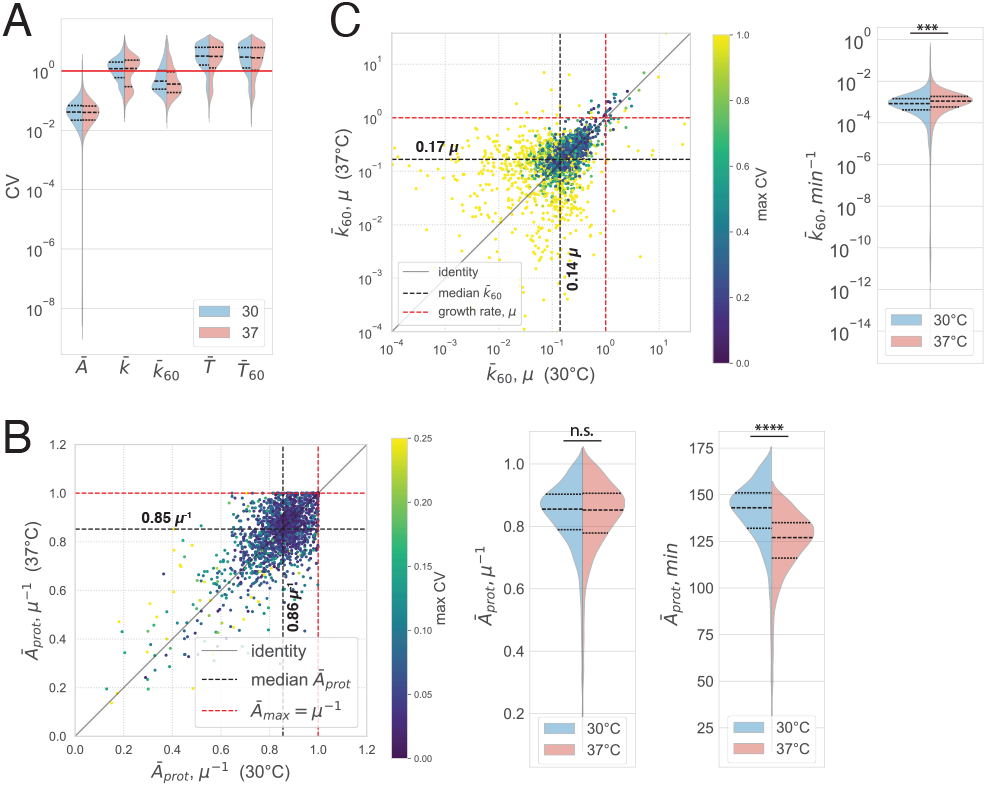
Dynamic parameters of yeast proteome at 30^°^C and 37^°^C. Dynamic parameters of yeast proteins and their coefficients of variation (CVs) quantified by fitting bootstrapped labeling dynamics with the 2-pool option (see Figure 11). (A) CV distribution of different dynamic parameters of yeast proteins. CV *<* 1 (below red line) was considered a criterion for robust quantification. Note that only mean metabolic ages 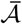 and expected decay rates for the age cohort older than 60 minutes 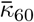 could be robustly determined for the majority of proteins. (B and C) Scatter plot (left) and cumulative distributions (right) of 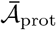 (B) and 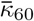 (C) at 30^°^C and 37^°^C. Note that systematic differences in mean protein ages between temperatures can be eliminated when expressing them in units of the maximum metabolic age, *µ*^−1^. ***: p *<* 0.001; ****: *p <* 0.0001 by paired Student’s t-test

Unlike metabolic ages, the quantification of the mean residence time and the expected decay rate was less reliable, resulting in median CVs greater than 1 (Figure 12A). The large uncertainties can be explained by the high sensitivity of these parameters to labeling in short time periods after the onset of wash-out (see Appendix S1.4.1), which cannot be accurately measured. Therefore, we attempted to quantify them for the age cohort older than 60 minutes, 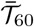 and 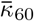, which should be less sensitive to the initial labeling dynamics (see Equation S22 and S25). Indeed, this restriction greatly reduced the uncertainties in the case of 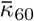 (Figure 12A). Despite the fact that 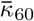 values below the median (0.14*µ* for 30^°^C or 0.17*µ* for 37^°^C) often had CVs greater than 1 (Figure 12C), we were still able to reliably characterize the stability of mature populations (older than 1 hour) for most proteins.

The decay rates naturally show the magnitude of biosynthetic flux diverted to degradation compared to that of cell growth, allowing one to formally define proteins with 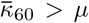, i.e., mostly degraded, as unstable. Although we were able to detect a handful of such proteins at both temperatures, most had a much lower 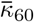 (median ≈ 0.15*µ*) pointing to the overall high stability of mature proteins. When expressing 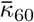 in absolute time units (e.g. *min*^−1^), there was only a minor global increase at 37^°^C (Figure 12C (right)), suggesting that mature proteins are generally not destabilized by the elevated temperature.

In conclusion, these results illustrate how the age-based framework can be used to flexibly quantify various dynamic parameters and examine the metabolic pool structure of proteins under different conditions based on the dynamic labeling data, take into account biases imposed by delayed input, and explain and evaluate the certainty in the evaluation of dynamic parameters.

## Discussion

Since the advent of pulse-chase labeling techniques in the 1940s [38, 39], dynamic characteristics of cellular metabolism, e.g., the time a molecule spends in certain parts of the metabolic system or its degradation rate, have been of central importance in understanding cellular homeostasis. In this work, we describe the analysis of steady-state cellular metabolism by dynamic labeling in the most general way. The key to our approach is the equivalence drawn between the experimentally observed labeling dynamics and the metabolic age distribution (Section 2.2). We use this equivalence to link the readouts of dynamic labeling experiments with various characteristics of metabolic dynamics including age, residence time, and decay rate (Sections 2.2–2.3).

To establish these connections, we used two alternative approaches to represent metabolic systems: either as a featureless “black box” (Section 2.3) or as a compartmental model (CMs) that explicitly describes the metabolic system as a set of compartments and their connecting fluxes (Section 2.6). Each of these descriptions has its own advantages. Black-box models, widely adopted in geophysics and geochemistry [4, 15, 16], are nonparametric, require fewer assumptions and can be used to describe the most basic principles that govern metabolic dynamics. Using this approach, we show that various dynamic parameters can be represented by continuous probability distributions or functions and are encoded in the geometry of the experimental labeling curve (Sections 2.2–2.3 and Figure 4). This approach was also instrumental in understanding how these connections are distorted by cell growth, delayed input, or non-steady-state metabolic dynamics (Sections 2.4–2.5), and how these distortions can be detected using experimental labeling dynamics (Figure 3).

Unlike the black-box representation, CMs mechanistically describe metabolic systems at an arbitrary level of detail by viewing them as a system of connected well-mixed compartments (Figure 8). Although specific CMs are sometimes used to interpret pulse-chase metabolic labeling experiments [7, 17, 18, 27], our study generalizes this description, capitalizing on a relatively well-developed mathematical backbone of CMs, the continuous-time domain Markov processes [32]. We demonstrate the utility of this framework by presenting several useful applications: inferring metabolic network topologies and dynamic parameters by fitting different models to measured data; interpolating sparse and noisy data from labeling dynamics experiments; or using CMs to compensate for delayed input (Section 2.6). We further show the practicality of these methods when applied to the analysis of the yeast proteome under mild heat stress (Section 2.8) and for determining the assembly order of the nuclear pore complex (Figure S8).

Beyond general descriptions, our analysis has important practical implications. We show that common stability characteristics – residence times and decay rates – depend on the first and second time derivatives of the labeling dynamics (Section 2.3). These interpretations are therefore highly sensitive to measurement noise. In particular, the accurate quantification of expected decay rates and mean residence times of bulk populations, often used as stability criteria, depends on measuring the initial labeling dynamics using fast and accurate sampling, which is usually unavailable. We demonstrate these limitations through simulations (Figure S10A) and by quantifying these parameters for yeast proteins based on experimental data (Figure 12A). Importantly, the full description of these parameters through probability distributions and age-dependent functions, presented in this study, enables their quantification in a more flexible and robust way. We illustrate this by describing the decay rates of the mature age cohort of yeast proteins at different temperatures where experimental data enable reliable quantification (Figure 12C).

We also address the issue of delayed input, i.e., when the label does not immediately enter the observed pool (Section 2.5). In such cases, many of the relationships between the dynamic parameters do not hold, which could lead to biases if delayed input is not taken into account. We show how these biases arise for residence times in simulations (Figure S10B) and for metabolic ages of yeast proteins using our experimental data (Figure S13). Although accounting for delayed input has been discussed in the past [7, 17, 18, 27, 35], here we provide a general framework based on the CM reduction approach (Sections 2.7 and S1.6.3). We demonstrate its utility by applying it to data from yeast dynamic SILAC experiments at different temperatures, where delayed lysine input had a significant impact on results (Figure 10). We envision that this approach facilitates aligning and comparing dynamic data, acquired by various labeling approaches and experimental conditions (e.g. using different amino acids as labels), in order to improve reproducibility. Similar delay effects arising from sequential processes may provide an explanation for the different labeling dynamics observed for the same molecule in different subcellular compartments or, for example, for different post-translationally modified protein isoforms, which are regularly attributed to underlying changes in stability [3, 19, 27, 45, 48, 49].

The key outcome of our analysis is that metabolic ages are directly quantifiable from labeling dynamics, while other dynamic parameters require additional assumptions such as the absence of delayed input and are susceptible to experimental measurement errors (Sections 2.2 and 2.5). Our analysis of the yeast proteome using dynamic labeling shows that metabolic age estimates were robust, while decay rates and residence times were not (Figure 12). Despite its advantages, age has been used as a metric of metabolic dynamics only in a few recent studies, such as [19, 30]. We provide a convenient way to rigorously quantify this parameter for target molecular pools and explore its physiological importance. As an example, by estimating the minimal timescale of NPC assembly without a priori assumptions (see Figure S8D), we show how metabolic age can be a useful measure for the time-scale of cellular processes.

As part of this study, we have developed an open-source Python package to simulate and fit CMs to dynamic labeling data aimed at aiding in the interpretation of the results of dynamic labeling experiments. Although there are publicly available compartmental modeling solutions such as `LAPM` and `CompartmentalSystems` created by Metzler et al. [32], our package adds many unique tools specifically useful to describe metabolic processes, such as modeling growing systems, implementing mass balance constraints, establishing methods for flexible extraction of various dynamic parameters, and enabling nonparametric methods for their quantification. The use of this solution is illustrated throughout this work in multiple practical examples (e.g., in Section 2.8 and Appendix S1.8.1).

## 3 Materials and Methods

### 3.1 Culture conditions, protein labeling and abundance quantification

Dynamic labeling assays were performed using a haploid BY4742 strain (*MATα his3*Δ*1 leu2*Δ*0 lys2*Δ*0 ura3*Δ*0* ) auxotrophic for lysine. Cell cultures were grown at 30^°^C or 37^°^C in standard synthetic complete medium containing 6.7 g/L yeast nitrogen base without amino acids and with ammonium sulfate, 2% glucose, necessary amino acids and specifically supplemented at 25 mg/L f.c. with normal [^12^C_6_,^14^ N_2_] L-lysine or heavy [^13^C_6_,^15^ N_2_] L-lysine isotopomer (Silantes), referred to as “light medium” and “heavy medium”, respectively. Light-to-heavy-medium exchange and sample collection was performed by filtration. The yeast samples were lysed using glass beads and cellular proteins were trypsinized by a standard in-gel digestion protocol. Tryptic peptides were analyzed with Eclipse Tribrid mass spectrometer in DIA mode, and quantified in light and heavy channels using DIA-NN 2.1.0 [11]. The fractional labeling of lysine in the protein pools and relative protein abundances were quantified based on the DIA-NN report (see Data, Materials, and Software Availability and Appendix for details).

### 3.2 Extraction of dynamic parameters using compartmental models

All CM instances used to extract dynamic parameters from labeling data or to simulate labeling curves were implemented with an open-source Python package, named `symbolic-compartmental-model`, that was developed as a part of this study (see Data, Materials, and Software Availability).

### 3.3 Bioinformatic analysis

All GO term analyses were performed using g:Profiler API [25] to acquire annotations and GO term sizes. GO term enrichment was assessed using the g:Profiler API (for 2-pool CM) or by local paired t-tests with Benjamini-Hochberg FDR correction. Gene overlaps with experimental data were computed locally. Detailed description of experimental procedures can be found in the Appendix.

### 3.4 Data, Materials, and Software Availability

The Python package for CM calculations can be found at https://gitlab.com/elad.noor/symbolic-compartmental-model. The documentation for this package can be found at https://symbolic-compartmental-model.readthedocs.io/. All Jupyter notebooks and the accompanying code that were used to analyze the labeling and growth rate data related to heat stress in yeast, as well as the bioinformatic analysis described above, can be found at https://doi.org/10.5281/zenodo.17101594. Data on protein abundance and dynamic metabolic labeling from previously published works [35] and algorithms used to define the order and timescale of the nuclear pore complex assembly, based on the metabolic age metric, are stored at https://doi.org/10.5281/zenodo.17116706.

## Acknowledgements

We thank Corentin Briat for his generous help in the mathematical proofs related to Metzler matrices, Jonas Fischer, Elisa Dultz, Yinon Bar-On, Melody Cools, and Wolfram Liebermeister for critically reading and providing useful feedback on the manuscript. Mass spectrometry-based proteomic analyses were performed by the Proteomics Unit at the University of Bergen (PROBE), a member of the National Network of Advanced Proteomics Infrastructure (NAPI) funded by the Research Council of Norway (INFRASTRUKTUR-program project number: 295910). This work was supported by a grant NFR 315615 awarded by the Research Council of Norway to Evgeny Onischenko and Elad Noor.

