## Supplementary Material for "Age-based approach to characterize the dynamics of cellular processes"

### S1 Appendix

#### S1.1 Glossary:

| Term | Definition |
|---|---|
| Metabolic system | a set of internal metabolic pools and their connecting fluxes |
| Nutrient | externally supplied metabolic material |
| Metabolite | nutrient's product resulting from internal chemical reactions |
| Metabolic age | the time the metabolite has spent in metabolic system |
| Residence time | time spent by the metabolite between the moment of entry and the moment of exit from the metabolic system or its parts |
| Decay rate | fractional escape rate from the metabolic system or its parts |
| Half-life | the time during which half of the labeled particles are washed out |
| Turnover | fractional exchange (replacement) rate |
| Compartmental model | a representation of a metabolic system as a set of compartments (states) with connecting fluxes representing material transfer rates between the compartments |

#### S1.2 Alternative scenarios for dynamic labeling

In Section 1.2, we demonstrate a direct connection between the labeling dynamics and the metabolic age distributions in steady-state dynamic labeling experiments. This additionally requires that the system begins with a fully labeled metabolite (at $t = 0$) and that the label is completely washed-out over time, so that the metabolite will be fully unlabeled at $t = \infty$. However, real-world experiments can deviate from this idealized scenario. In addition, there are other experimental designs, such as pulse-chase or step-up labeling, rather than wash-out. Here, we relax many of our assumptions and show how the labeling function can still be used to represent the metabolic age distribution.

##### S1.2.1 Incomplete labeling

We assume, as before, that $f(t)$ represents the labeled fraction. However, $f(0)$ can also have a value smaller than 1, and $f(\infty)$ can be larger than 0. Furthermore, $f(t)$ must be monotonic, but it can either increase or decrease, depending on the type of labeling shift in the medium (step-up or wash-out). In all cases, the relationship between the age CDF and the labeling will be:

$$P_A(t) = \frac{f(0) - f(t)}{f(0) - f(\infty)}\,. \tag{S1}$$

*E.N., K.J. and E.O. contributed equally to this work

| Variable/Function | Definition |
|---|---|
| $p_X(x)$ or $p_X(x)$ | probability density function (PDF) of random variable $X$ |
| $P_X(x)$ or $P_X(x)$ | cumulative density function (CDF) of random variable $X$ |
| $\mathcal{A}$ and $\bar{\mathcal{A}}$ | age and mean age |
| $\mathcal{T}$ and $\bar{\mathcal{T}}$ | residence time and mean residence time |
| $\kappa$ and $\bar{\kappa}$ | decay rate and expected decay rate |
| $\lambda$ and $\bar{\lambda}$ | total escape rate and expected total escape rate |
| $\kappa_{j\to i} \equiv \frac{v_{j\to i}}{S_i}$ | contributed turnover rate |
| $f$ | labeled fraction $\frac{\text{labeled}}{\text{labeled+unlabeled}}$ |
| $\mu$ | cell growth rate |

We can appreciate that $P_{\mathcal{A}}(0) = 0$ and $P_{\mathcal{A}}(\infty) = 1$, as required from the definition of the age CDF. The downside, however, is that the values of $f(0)$ and $f(\infty)$ must be measured or estimated.

Using this generalized form is important in several common experimental setups, such as:

1. When the natural abundance of the labeled isotope is not zero (e.g. $^{13}$C which has an abundance of ≈1%).
2. When using a pure 100% labeled version of the metabolite is difficult (e.g., very expensive).
3. When we can only detect the labeled metabolites, but not the unlabeled ones, for example, signal intensities of a fluorescently labeled protein. Since we cannot calculate the fraction of labeling for each time-point individually, Equation S1 is an alternative way to do the normalization. In this case $f(t)$ represents the observed amount of label, e.g., label signal intensity, and can have any positive value.

The last scenario is relevant when using fluorescent proteins or radioisotopic labeling.

###### S1.2.2 Pulse labeling

In pulse-chase experiments, a short metabolic labeling pulse (usually employing radioisotopic labels) is followed by a chase period during which the biomass-normalized signal intensities of the internal pools are analyzed. Unlike the wash-out experiments discussed throughout this work, the labeling of pulse-chase experiments *does not* directly correspond to the age CDF. The reason is that the label is introduced as an input only for a short (infinitesimal) time-period – not as a step function. However, we will show that the labeling function in pulse-chase experiments is proportional to the age PDF.

First, we assume that the pulse itself is very short (in comparison to the internal dynamics in the system). This means that at any time point $t > 0$, every labeled particle in the system is exactly of age $\mathcal{A} = t$. Therefore, measuring the amount of labeling over time (effectively, until it becomes undetectable) will be proportional to the distribution of ages, with a prefactor that depends on the total amount of label. Since the integral of $p_{\mathcal{A}}(t)$ must be equal to 1, we can use this to normalize the labeling curve and eliminate this prefactor.

Importantly, one must have enough measurements (frequent and across the entire relevant timescale) in order to calculate this integral. Alternatively, when fitting data to a CM, one can add the scaling factor as an extra fitting parameter.

Finally, the criteria for steady-state and delayed inputs need to be adjusted: when the input is not delayed, the labeling curve must be monotonically decreasing (but not necessarily convex). If the input is delayed, the function could increase at a certain time period, even if the system is at steady-state.

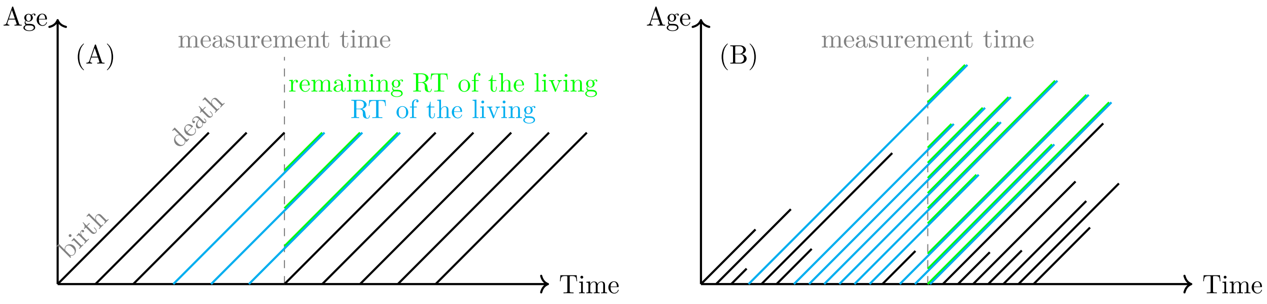


Figure S1: An idealized example of a population of particles, each represented by a single line. The x-axis represents absolute time and the y-axis represents the age of single particles. Particles are born (enter the system) in different times, and age at the same rate as time passes by, until they reach death (leave the system). (A) The residence time of the living is highlighted in cyan color, and the remaining residence time of the living is in bright green. (B) A plot with randomly sampled data, based on a simulation of a 2-state CM. The (remaining) residence time of the living is again marked by (green) cyan color.

#### S1.3 Dynamic characteristics of non-growing steady-state metabolic systems as function of labeling dynamics

##### S1.3.1 Mean age

To quantify the mean metabolic age we can treat metabolic age $A$ as a random value whose mean is equivalent to its expectancy:

$$\bar{\mathcal{A}} \equiv \mathbb{E}[A] = \int_0^\infty t \; p_\mathcal{A}(t) \; dt \tag{S2}$$

Using the equivalence of metabolic age distributions and labeling dynamics (see Equation 2) we can also express $\bar{\mathcal{A}}$ as a function of the labeling dynamics and solve it using integration by parts:

$$\bar{\mathcal{A}} = -\int_0^\infty t \; \dot{f}(t) \; dt = -t \; f(t)\Big|_0^\infty + \int_0^\infty 1 \; f(t) \; dt \tag{S3}$$

Any real metabolic system can be described by a compartmental model, $f(t) = \mathbf{s}^\top e^{\mathbf{M}t}\mathbf{1}_n$, where $\mathbf{M}$ has no positive eigenvalues. Based on Lemma S1.21, we can say with loss of generality that $\lim_{t\to\infty} t \; f(t) = \mathbf{s}^\top \left(\lim_{t\to\infty} t \; e^{\mathbf{M}t}\right) \mathbf{1}_n = 0$. Therefore, the expression for the mean age can be simplified to:

$$\bar{\mathcal{A}} = \int_0^\infty f(t) dt \tag{S4}$$

The mean metabolic age is therefore equal to the area under the labeling curve (AUC), $f(t)$.

##### S1.3.2 Mean residence time in non-growing systems

We have shown in Equation 5 that there is a direct relationship between the residence time CDF and the age PDF:

$$P_\mathcal{T}(t) = \; 1 - \frac{p_\mathcal{A}(t)}{p_\mathcal{A}(0)} \, . \tag{S5}$$

It is safe to assume that $p_\mathcal{A}(t)$ will eventually decrease exponentially to 0 at long enough time scales, since every molecule has a finite residence time (even if very large). Therefore, $P_\mathcal{T}(t)$ fulfills the condition of Corollary S1.4.1, and we can use it to calculate the mean residence time:

$$\bar{\mathcal{T}} \equiv \mathbb{E}[\mathcal{T}] = \int_0^\infty [1 - P_\mathcal{T}(t)] \; dt = \int_0^\infty \frac{p_\mathcal{A}(t)}{p_\mathcal{A}(0)} \; dt = \frac{1}{p_\mathcal{A}(0)} \underbrace{\int_0^\infty p_\mathcal{A}(t) \; dt}_{=1} = -\frac{1}{\dot{f}(0)} \tag{S6}$$

where in the last step we use the fact that the integral of the age PDF over the entire range of ages is, by definition, 1 and replaced $p_\mathcal{A}(0)$ with the $-\dot{f}(0)$ based on Equation 3.

##### S1.3.3 Remaining residence time of the living

This is the residence time that a randomly selected (living) particle has left, i.e., its residence time minus its age (see Figure S1), which we denote by $\mathcal{L}_r$ and by the acronym RRTL. At any specific time point, $P_{\mathcal{L}_r}(t)$ represents the fraction of particles whose RRTLs are smaller than $t$. If we wait until time $t$, only they will be replaced by new (unlabeled) particles, while all the others will remain labeled in the system. Therefore, $P_{\mathcal{L}_r}(t)$ is equal to the unlabeled fraction, or $1 - f(t)$. If the system is at steady-state, $1 - f(t)$ is also equal to the age CDF, so we can conclude that:

$$P_{\mathcal{L}_r}(t) = P_{\mathcal{A}}(t)\,. \tag{S7}$$

and therefore also the mean will be equal to the mean age: $\bar{\mathcal{L}}_r = \bar{\mathcal{A}}$.

Interestingly, if the system is **not** at steady-state, then the RRTL distribution will still correspond to the labeling curve, but the age distribution will not.

##### S1.3.4 Residence time of the living in non-growing systems

The residence time of the living (RTL) is defined as the time between birth and death (escape) for a randomly chosen (living) particle (see Figure S1) and, as shown in Wrigley-Field and Feehan [2022] for steady-state systems, it is a length-biased transformation of the (standard) residence time. Below, we provide here an alternative proof which reaches the same conclusion.

As the symbol for the random variable representing the RTL, we will use $\mathcal{L}$. We can express it in terms of the age and RRTL:

$$\mathcal{L} = \mathcal{A} + \mathcal{L}_r\,. \tag{S8}$$

Importantly, these random variables are not necessarily independent. Therefore, we cannot use a simple convolution to derive an expression for the distribution of the RTL based on the other two distributions.

To calculate the CDF of the RTL, we will count all of the *living* particles that are going to have a residence time shorter than $t$, by considering two mutually distinct options: (i) if the age of the particle is already larger than $t$ then its residence time will definitely also be larger and we can simply ignore it, (ii) otherwise, the age of the particle is some $\tau \leq t$ and the probability of it escaping before reaching age $t$ is given by $1 - p_{\mathcal{A}}(t)/p_{\mathcal{A}}(\tau)$. Therefore, we can find the total distribution by integrating the probability of having age $\tau$ times the probability of escaping before reaching age $t$:

$$P_{\mathcal{L}}(t) = \int_0^t p_{\mathcal{A}}(\tau) \cdot \left(1 - \frac{p_{\mathcal{A}}(t)}{p_{\mathcal{A}}(\tau)}\right) d\tau = \int_0^t p_{\mathcal{A}}(\tau) d\tau - \int_0^t p_{\mathcal{A}}(t) d\tau = P_{\mathcal{A}}(t) - t \cdot p_{\mathcal{A}}(t) = 1 - f(t) + t\dot{f}(t) \tag{S9}$$

and by taking the time derivative, we can also find the PDF:

$$p_{\mathcal{L}}(t) = -\dot{f}(t) + \dot{f}(t) + t\ddot{f}(t) = t\ddot{f}(t)\,. \tag{S10}$$

**Mean residence time of the living** Similar to the residence time itself, $P_{\mathcal{L}}(t)$ fulfills the condition of Corollary S1.4.1, and we can use it to calculate the mean residence time of the living:

$$\bar{\mathcal{L}} \equiv \mathbb{E}[\mathcal{L}] = \int_0^\infty [1 - P_{\mathcal{L}}(t)] dt = \int_0^\infty f(t) - \int_0^\infty t\dot{f}(t) dt \tag{S11}$$

By observing the two integrals in the last expressions to Equations S3 and S4, we can see that both are equal to the mean age. Therefore, the mean residence time of the living is twice as large as the mean age of the population:

$$\bar{\mathcal{L}} = 2\bar{\mathcal{A}} \tag{S12}$$

##### S1.3.5 Expected decay rate in non-growing systems

We can now use a similar workflow to determine the expected decay rates. In Equation 7, we found an expression for the decay rate, $\kappa$, as a function of the age distribution:

$$\kappa(t) = -\frac{\dot{p}_{\mathcal{A}}(t)}{p_{\mathcal{A}}(t)}\,. \tag{S13}$$

The *expected decay rate* is defined as the decay rate's expectancy over the probability distribution of ages. This can be expressed as:

$$\bar{\kappa} \equiv \mathbb{E}[\kappa(\mathcal{A})] = \int_0^\infty \kappa(t)\, p_{\mathcal{A}}(t)\, dt = -\int_0^\infty \dot{p}_{\mathcal{A}}(t)\, dt = -p_{\mathcal{A}}(t)\Big|_0^\infty = \cancel{-p_{\mathcal{A}}(\infty)} + p_{\mathcal{A}}(0) = -\dot{f}(0) \tag{S14}$$

where, as before, we assume without loss of generality that $p_{\mathcal{A}}(\infty) = 0$.

The expected decay can also be seen as the overall fractional escape rate of the system particles and quantified as the sum of decay rates over all particle ages weighted by the contribution of each age $p_{\mathcal{A}}(t)dt$:

$$\bar{\kappa} = \int_0^\infty \kappa(t) p_{\mathcal{A}}(t) dt \equiv \dot{r}_{out} \tag{S15}$$

Since in steady state the escape and entry rates are the same, $\bar{\kappa}$ is equivalent to the total fractional exchange rate of the system particles also known as *turnover rate.*

###### S1.3.6 Age-conditioned expressions for dynamic parameters of metabolic systems

Attempting to experimentally determine the mean metabolic ages would require information about labeling dynamics at infinitely long timescales (see Equation 4). Therefore, one cannot perform the calculation purely based on empirical data and without making assumptions for how to extrapolate it. As an alternative to extrapolations, quantification of mean metabolic ages can be restricted to age intervals within the timescales of dynamic labeling experiments.

**Age-conditioned mean age** Using Equation 2, we can define the age-conditioned mean value of metabolic age and its connection to dynamic labeling readouts at restricted age intervals $(0, a)$ as follows:

$$\mathbb{E}[\mathcal{A} \le a] = \frac{-\int_0^a t\dot{f}(t)dt}{P_{\mathcal{A}}(a)} = \frac{-tf(t)\Big|_0^a + \int_0^a f(t)dt}{P_{\mathcal{A}}(a)} = \frac{\int_0^a f(t)dt - af(a)}{1 - f(a)} \tag{S16}$$

The problem of extremes also arises when determining mean/expected values for residence times and decay rates, as they both require knowing the labeling dynamics infinitely close to time of labeling initiation. As an alternative, these parameters can also be conditioned to the experimentally covered timescales.

**Age-conditioned expected decay rate** We define it as the expected value of the decay rates for an interval between $(a, \infty)$. It can be connected to labeling dynamics using Equation S13:

$$\bar{\kappa}_a \equiv \mathbb{E}[\kappa(\mathcal{A}) \mid \mathcal{A} \ge a] = \frac{\int_a^\infty \kappa(t)\; p_{\mathcal{A}}(t)\; dt}{P_{\mathcal{A}}(a)} = \frac{-\int_a^\infty \dot{p}_{\mathcal{A}}(t)\; dt}{f(a)} = \frac{-p_{\mathcal{A}}(t)\Big|_a^\infty}{f(a)} = -\frac{\dot{f}(a)}{f(a)} \tag{S17}$$

**Age-cohort residence time** In the case of residence times, instead of the age-conditioned probabilities (i.e., $\mathcal{A} \ge a$), it is more useful to consider age-cohort probabilities (i.e., $\mathcal{A} = a$). Since the population of particles which are at a certain age $(a)$ are the same as those that have residence times larger or equal to $a$, we can use the notation $\mathcal{T} \ge a$ (see Figure S2). This can be practical in excluding events with short residence times much shorter than the experimentally available timescales. We therefore define the *age-cohort residence time* CDF as $P_{\mathcal{T}}(t \mid \mathcal{T} \ge a)$ – i.e., the probability that the residence time of a particle is between $a$ and $t$. Based on Equation 8, the exponent of the integral of $-\kappa(\mathcal{A})$ between 0 and $t$ describes the fraction of remaining particles at time $t$ (i.e. those who have a residence time larger than $t$). We can generalize this, by starting the integration at some arbitrary age $a$ instead of 0, thus accounting only for the decay between the ages $a$ and $t$, which will exactly give us the reciprocal CDF of the age-cohort residence time:

$$1 - P_{\mathcal{T}}(t \mid \mathcal{T} \ge a) = \mathrm{e}^{-\int_a^t \kappa(\tau)d\tau} = \frac{\dot{f}(t)}{\dot{f}(a)}\,. \tag{S18}$$

Note that for any $t \le a$, $P_{\mathcal{T}}(t \mid \mathcal{T} \ge a) = 0$ by definition. In summary, the distribution of the age-cohort residence time is given by:

$$\begin{aligned} P_{\mathcal{T}}(t \mid \mathcal{T} \ge a) &= \begin{cases} 0 & \text{if } t \le a \\ 1 - \frac{\dot{f}(t)}{\dot{f}(a)} & \text{if } t > a \end{cases} \\ p_{\mathcal{T}}(t \mid \mathcal{T} \ge a) &= \begin{cases} 0 & \text{if } t \le a \\ -\frac{\ddot{f}(t)}{\dot{f}(a)} & \text{if } t > a \end{cases}. \end{aligned} \tag{S19}$$

We can further calculate the age-cohort mean residence time, and denote it using a special symbol:

$$\bar{\mathcal{T}}_a \equiv \mathbb{E}[\mathcal{T} | \mathcal{T} \ge a] = \int_0^\infty \big[1 - P_{\mathcal{T}}(t | \mathcal{T} \ge a)\big]\, dt = \int_0^a 1\; dt\; + \int_a^\infty \frac{\dot{f}(t)}{\dot{f}(a)}\; dt = a - \frac{f(a)}{\dot{f}(a)}\,. \tag{S20}$$

See Figure S3 for a geometrical interpretation of this solution.

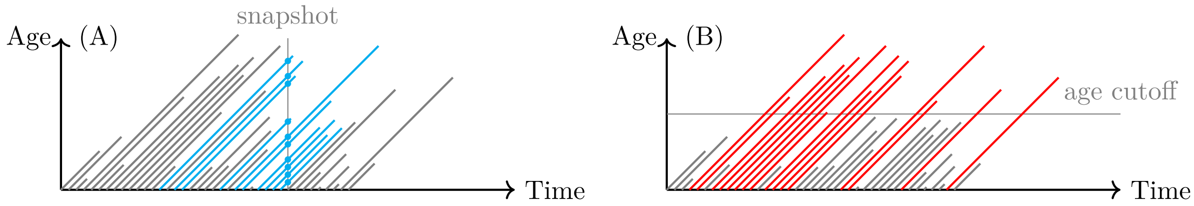


Figure S2: An example of a population of particles, whose timelines are represented by diagonal lines. The x-axis represents absolute time and the y-axis represents the age of single particles. When a particle enters the system, it starts with an age of 0, and as time passes by, it ages at the same rate (therefore, each timeline is at a 45° angle above the x-axis). When the particle leaves the system, its timeline is terminated, and the age at which this happens is the residence time. (A) We can imagine a cross-section at some time $t$ by drawing a vertical line representing a snapshot of the system at that time. The y-values of the intersections with the active timelines, marked by blue circles, represent the age distribution at time $t$. (B) As an example, the age-cohort for one specific age is marked by orange color, i.e. only those timelines that cross the cutoff age.

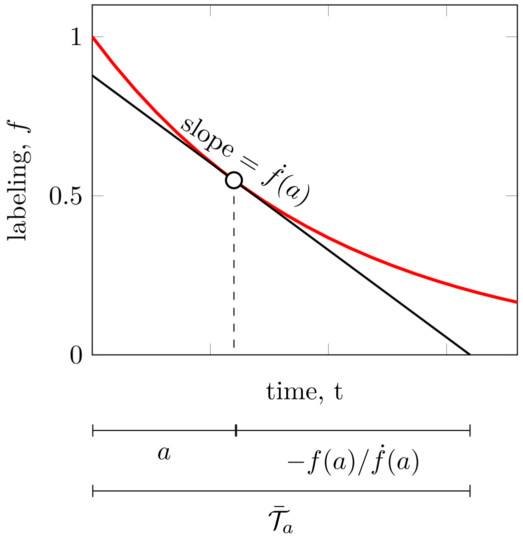


Figure S3: **Geometric interpretation of the age-cohort mean residence time.** Based on Equation S20, we can see that $\bar{\mathcal{T}}_a$ corresponds to the x-intercept of the tangent line to $f(a)$.

#### S1.4 Dynamic characteristics of exponentially growing systems

##### S1.4.1 Mean residence time and expected decay rate in exponentially growing systems

To express the expected values for the decay rate in growing systems, we can use the fact that $\kappa(\mathcal{A})$ is the same as the finite-volume total escape rate, $\lambda(\mathcal{A})$, up to a constant offset $\mu$ (see Equation 12):

$$\bar{\kappa} \equiv \mathbb{E}[\kappa(\mathcal{A})] = \mathbb{E}[\lambda(\mathcal{A}) - \mu] = \mathbb{E}[\lambda(\mathcal{A})] - \mu = \bar{\lambda} - \mu = -\dot{f}(0) - \mu \tag{S21}$$

The *age-conditioned expected decay rate* will similarly be affected:

$$\bar{\kappa}_a - \mu = -\frac{\dot{f}(a)}{f(a)} - \mu \tag{S22}$$

To express the mean residence time, we can again use Corollary S1.4.1. First, we note that $e^{\mu t} p_{\mathcal{A}}(t)$ is an exponentially decaying function because the residence time must be finite. Therefore, $P_{\mathcal{T}}(t)$ fulfills the conditions of the Corollary and we can use it to compute the mean residence time (using Equation 13):

$$\bar{\mathcal{T}} \equiv \mathbb{E}[\mathcal{T}] = \int_0^\infty [1 - P_{\mathcal{T}}(t)]\, dt = \int_0^\infty e^{\mu t}\, \frac{p_{\mathcal{A}}(t)}{p_{\mathcal{A}}(0)}\, dt = \frac{1}{\dot{f}(0)} \int_0^\infty \dot{f}(t)\, e^{\mu t}\, dt\,. \tag{S23}$$

For a more explicit solution in the case of CMs, see Appendix S1.6.5.

We can also derive, in a similar way, the age-cohort residence time for a growing system:

$$\begin{aligned} P_{\mathcal{T}}(t \mid \mathcal{T} \geq a) &= \begin{cases} 0 & \text{if } t \leq a \\ 1 - \frac{\mathrm{e}^{\mu t}}{\mathrm{e}^{\mu a}} \frac{\dot{f}(t)}{\dot{f}(a)} & \text{if } t > a \end{cases} \\ p_{\mathcal{T}}(t \mid \mathcal{T} \geq a) &= \begin{cases} 0 & \text{if } t \leq a \\ -\frac{\mathrm{e}^{\mu t}}{\mathrm{e}^{\mu a}} \frac{\left(\ddot{f}(t) + \mu \dot{f}(t)\right)}{\dot{f}(a)} & \text{if } t > a \end{cases} \end{aligned} \tag{S24}$$

and use it to calculate the age-cohort mean residence time:

$$\bar{\mathcal{T}}_a = \int_0^\infty \left[1 - P_{\mathcal{T}}(t \mid \mathcal{T} \geq a)\right] dt \; = \int_0^a 1 \; dt \; + \int_a^\infty \frac{\mathrm{e}^{\mu t}}{\mathrm{e}^{\mu a}} \frac{\dot{f}(t)}{\dot{f}(a)} \; dt = a + \frac{\int_a^\infty \mathrm{e}^{\mu t} \; \dot{f}(t) \; dt}{\mathrm{e}^{\mu a} \; \dot{f}(a)} \tag{S25}$$

which converges to $a - f(a)/\dot{f}(a)$ when $\mu = 0$, as expected from Equation S20.

##### S1.4.2 Remaining residence time of the living in exponentially growing systems

In exponentially growing systems, the only thing that changes with respect to the remaining residence time is that the fraction of particles that remain in the system after time $t$ needs to be multiplied by $\mathrm{e}^{\mu t}$ to represent the fact that the system is growing in this rate, and therefore can contain more particles. Therefore, the general expressions for the distribution of the RRTL are:

$$\begin{aligned} P_{\mathcal{L}_r}(t) &= 1 - \mathrm{e}^{\mu t} f(t) \\ p_{\mathcal{L}_r}(t) &= -\mathrm{e}^{\mu t}\left(\mu f(t) + \dot{f}(t)\right) . \end{aligned} \tag{S26}$$

The mean RRTL in a growing system will be the integral:

$$\bar{\mathcal{L}}_r = \mathbb{E}[\mathcal{L}_r] = \int_0^\infty [1 - P_{\mathcal{L}_r}(t)] dt = \int_0^\infty \mathrm{e}^{\mu t} f(t) dt \tag{S27}$$

##### S1.4.3 Residence time of the living in exponentially growing systems

As we have seen in Appendix S1.3.4, the probability of particles with age $\tau \leq t$ to still remain in the system at time $t$ is given by $p_{\mathcal{A}}(t)/p_{\mathcal{A}}(\tau)$. During this time (which is time period of length $t - \tau$), the system expands and will grow by a factor of $\mathrm{e}^{\mu(t-\tau)}$. So, the fraction of particles that remains will be given by $\mathrm{e}^{\mu(t-\tau)} p_{\mathcal{A}}(t)/p_{\mathcal{A}}(\tau)$, and the ones that escape (i.e. have a residence time smaller than $t$) is one minus that. As before, we can integrate this probability times the age PDF over $\tau$ (between 0 and $t$) to get the cumulative probability:

$$\begin{aligned} P_{\mathcal{L}}(t) &= \int_0^t p_{\mathcal{A}}(\tau) \cdot \left(1 - \frac{\mathrm{e}^{\mu(t-\tau)} p_{\mathcal{A}}(t)}{p_{\mathcal{A}}(\tau)}\right) d\tau = \int_0^t p_{\mathcal{A}}(\tau) d\tau - p_{\mathcal{A}}(t) \int_0^t \mathrm{e}^{\mu(t-\tau)} d\tau \\ &= P_{\mathcal{A}}(t) + p_{\mathcal{A}}(t) \frac{1 - \mathrm{e}^{\mu t}}{\mu} \end{aligned} \tag{S28}$$

Therefore, the general expressions for the CDF and PDF of the residence time of the living in growing systems is:

$$\begin{aligned} P_{\mathcal{L}}(t) &= 1 - f(t) + \frac{\mathrm{e}^{\mu t} - 1}{\mu} \dot{f}(t) \\ p_{\mathcal{L}}(t) &= \frac{\mathrm{e}^{\mu t} - 1}{\mu} \left(\mu \dot{f}(t) + \ddot{f}(t)\right) . \end{aligned} \tag{S29}$$

for $\mu \to 0$, we can apply L'Hôpital's rule which will give us $\frac{\mathrm{e}^{\mu t} - 1}{\mu} \to t$, consistent with the result for non-growing systems (Appendix S1.3.4).

The mean residence time of the living in a growing system can once again be found by integration:

$$\begin{aligned} \bar{\mathcal{L}} &= \int_0^\infty [1 - P_{\mathcal{L}}(t)] dt = \int_0^\infty \left(f(t) + \frac{1 - \mathrm{e}^{\mu t}}{\mu} \dot{f}(t)\right) dt = \int_0^\infty f(t) dt + \frac{1}{\mu} \int_0^\infty \left(1 - \mathrm{e}^{\mu t}\right) \dot{f}(t) dt \\ &= \bar{\mathcal{A}} \; + \; \frac{1}{\mu} \left( \cancel{(1 - \mathrm{e}^{\mu t}) f(t) \Big|_0^\infty} + \int_0^\infty \mu \mathrm{e}^{\mu t} f(t) dt \right) = \bar{\mathcal{A}} + \int_0^\infty \mathrm{e}^{\mu t} f(t) dt \, . \end{aligned} \tag{S30}$$

Comparing this to Equation S27, we can see that the even in growing systems, the relationship between the mean values still holds:

$$\bar{\mathcal{L}} = \bar{\mathcal{A}} + \bar{\mathcal{L}}_r \tag{S31}$$

##### S1.4.4 Limit for labeling dynamics in exponentially growing systems with direct input

If a system has a direct input, i.e. one that is labeled without a delay, and is also growing exponentially, the dynamics of the labeled fraction cannot exceed the dilution due to system growth, $\mathrm{e}^{-\mu t}$. To show this, we will use the fact that the total escape rates from a fixed volume in an exponentially growing system ($\lambda$) cannot be lower than the biomass growth rate ($\mu$). Then we can write:

$$\begin{aligned} -\ddot{f}(t)/\dot{f}(t) &= \lambda \geq \mu \\ \ddot{f}(t) &\geq -\mu \dot{f}(t) \\ \int_t^\infty \ddot{f}(t')\ dt' &\geq -\int_t^\infty \mu\ \dot{f}(t')dt' \\ \cancel{\dot{f}(\infty)} - \dot{f}(t) &\geq -\mu(\cancel{f(\infty)} - f(t)) \\ \dot{f}(t) &\leq -\mu f(t) \\ \int_0^t \left(\dot{f}(t')/f(t')\right)\ dt' &\leq -\int_0^t \mu\ dt' \\ \ln(f(t)) - \cancel{\ln(f(0))} &\leq -\mu t \\ f(t) &\leq \mathrm{e}^{-\mu t} \end{aligned} \tag{S32}$$

where we used $f(0) = 1$ as well as the reasonable assumption that both $f(t)$ and $\dot{f}(t)$ reach 0 at the limit $t \to \infty$, which is a fair assumption considering that the real dynamic systems are representable by compartmental models that always have this property (see Lemma S1.21).

In conclusion, if at any point the value of $f(t)$ exceeds this threshold, the input into the system is certainly delayed.

Another consequence of this limit is that metabolic ages (mean values) in directly labeled growing systems cannot be greater than the inverse growth rate $\mu^{-1}$. To see why, we simply use the above result in the expression for mean age (Equation S4):

$$\bar{\mathcal{A}} = \int_0^\infty f(t)dt \leq \int_0^\infty \mathrm{e}^{-\mu t}dt = -\mu^{-1}\mathrm{e}^{-\mu t}\Big|_0^\infty = -\mu^{-1}(0-1) = \mu^{-1}\,. \tag{S33}$$

##### S1.4.5 Exponentially growing systems with constant decay rates

Although mean residence times and expected decay rates are not inversely related in general, the inverse relation still holds for growing systems with a constant (age-independent) decay rate. To demonstrate this, we will first determine the labeling dynamics of such systems and then use it to quantify the mean residence times and compare them with the decay rate.

To determine the labeling dynamics we will focus on a fixed volume within the growing system and compute its labeling dynamics as a function of the system's decay rate $\kappa$ by using Equation 7:

$$\ddot{f}(t) = -\lambda \cdot \dot{f}(t) \tag{S34}$$

In this case, $\lambda$ is constant, since both $\kappa$ and $\mu$ are constant. Furthermore, we can constrain the solution to have $f(0) = 1$ and $f(\infty) = 0$ (i.e., the labeled fraction starts as 1 and is eventually washed away completely). The general solution to this ODE is $f(t) = \mathrm{e}^{-\lambda t}$, which we can then use in Equation S23 and compute $\bar{\mathcal{T}}$:

$$\bar{\mathcal{T}} = \frac{1}{-\lambda}\int_0^\infty (-\lambda)\mathrm{e}^{-\lambda t}\ \mathrm{e}^{\mu t}\ dt = \int_0^\infty \mathrm{e}^{-(\lambda-\mu)t}\ dt = -(\lambda-\mu)^{-1}\mathrm{e}^{-(\lambda-\mu)t}\Big|_0^\infty = (\lambda-\mu)^{-1}\,. \tag{S35}$$

Finally, since $\kappa$ is constant, its expected value is trivial, and we can again use Equation 12 to compare it to the decay rate:

$$\bar{\kappa} = \kappa = \lambda - \mu\,. \tag{S36}$$

In conclusion, the mean residence time and expected decay rate can also be inversely related for growing systems, but only if the decay rate rate is age-independent.

Systems with a constant decay rate are also characterized by simple exponential metabolic age distributions. Differentiating the solution of equation S34, we get the expression for metabolic age distributions in such systems:

$$p_{\mathcal{A}}(t) = -\dot{f}(t) = \lambda \mathrm{e}^{-\lambda t} \tag{S37}$$

We can also easily calculate the mean metabolic ages as follows:

$$\bar{\mathcal{A}} = \int_0^\infty f(t)dt = \int_0^\infty \mathrm{e}^{-\lambda} dt = \lambda^{-1} \tag{S38}$$

Finally, using equations S35 and S38 we can relate the mean age and the mean residence time:

$$\bar{\mathcal{T}} = (\lambda - \mu)^{-1} = (\bar{\mathcal{A}}^{-1} - \mu)^{-1} \tag{S39}$$

#### S1.5 Dynamics of systems with delayed input

##### S1.5.1 Initial labeling dynamics in indirectly fed systems

If all the input into the analyzed system is indirect (i.e. delayed), the observed initial labeling will not change, i.e. $\dot{f}(t=0) = 0$, and the labeling curve will always have an inflection point. We can illustrate this using an exemplary system fed by a well-mixed source and expressing the slope of the labeling curve at $t = 0$ (as in Equation 16):

$$\dot{f}(0) = -p_{\mathcal{A}}(0) = -\int_0^0 p_{\mathcal{A}^\circ}(x)\ p_{\mathcal{A}_r}(t-x)\ dx = 0 \tag{S40}$$

where we used the fact that the PDFs of $\mathcal{A}^\circ$ and $\mathcal{A}_r$ are finite. In addition, as a PDF, we know that $p_{\mathcal{A}}(t) \geq 0$ and the integral $\int_0^\infty p_{\mathcal{A}}(t)dt \equiv 1$. Therefore, at infinity, the probability must converge to zero, that is, $p_{\mathcal{A}}(\infty) = 0$, otherwise the integral would not be finite.

So, we know that $\dot{f}(0) = \dot{f}(\infty) = 0$. If $f$ were a convex function, then $\dot{f}$ would be monotonically non-increasing. Due to both limits being zero, that would necessarily mean $\dot{f}(t) = 0$ for any $t > 0$. This contradicts the fact that $f$ is not constant (it goes down from 1 to 0), and therefore we conclude that $f$ is not convex and must have an inflection point.

Using Equation S32 it is also obvious that in systems with delayed input, the labeling dynamics will always be slower than growth dilution for at least some initial period.

##### S1.5.2 Labeling dynamics of multiply-labeled metabolites in systems with delayed input

So far, we have only considered cases where each metabolite particle can be labeled (once) or unlabeled. However, there are cases where the analyzed molecules can have multiple potential labeling sites, e.g. tryptic peptides that originate from the same protein and can be labeled by a heavy amino acid at one, two or more positions (see Figure S4). Labeling of such multiply labeled metabolites can be viewed as a generalization of single labeling defined as the fully labeled fraction measured among the fully labeled and fully unlabeled metabolite species:

$$h(n,t) \equiv \frac{r(n,n,t)}{r(n,n,t) + r(0,n,t)}\,, \tag{S41}$$

where $r(i,n,t)$ is the fraction of molecules that are labeled at $i$ positions out of $n$ total, when measured at time $t$ after the onset of the wash-out. Following this definition, we can see that for single-labeled metabolites $h(1,t) = f(t)$.

Given that metabolites with different labeling multiplicities have the same age distribution (for example, tryptic peptides originating from the same protein but having different numbers of amino acid labeling positions), we can ask how the labeling dynamics for different multiplicities is affected by delayed input. To compare these dynamics we begin with formal definitions.

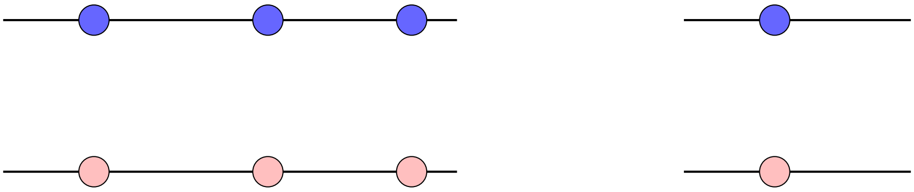

Figure S4: A schematic depiction of two peptides, one with three residues that can be labeled and another with only one. Over time, the labeled amino acids (pink) are washed-out and replaced by unlabeled versions of the same amino acid (blue).

**Definition S1.1.** Let $f^\circ(t)$ represent an input delay and be a monotonically decreasing labeling function over the real positive numbers, $f^\circ(0) = 1$, $f^\circ(t) > 0$, and $\lim_{t\to\infty} f^\circ(t) = 0$. We define the half-life $t_{1/2}$ as the time when $f^\circ$ is equal to $1/2$. Let $p_{\mathcal{A}}$ represent the age PDF of the single/multiple-labeled metabolites.

At time $t$, every position of a newly synthesized molecule would be labeled with probability $f^\circ(t)$ and unlabeled with probability $1 - f^\circ(t)$. Since the different sites are independent, the distribution of the total number of labels would follow a binomial distribution. So, the probability at time $t$ that a new molecule will have $i$ labeled sites (out of $n$ possible) would be:

$$\binom{n}{i}\Big(f^\circ(t)\Big)^i\Big(1 - f^\circ(t)\Big)^{n-i} \tag{S42}$$

In order to calculate $r(i,n,t)$, which includes molecules of all ages between 0 and $t$, rather then only the newly synthesized ones, we can use a convolution similar to Equation S129:

$$r(i,n,t) = \int_0^t \binom{n}{i}\Big(f^\circ(t-x)\Big)^i\Big(1 - f^\circ(t-x)\Big)^{n-i} p_{\mathcal{A}}(x)dx\,. \tag{S43}$$

**Lemma S1.2.** *For any $n \in \mathbb{N}$ and $t \in [0, t_{1/2})$:*

$$h(n,t) < h(n+1,t)$$

.

*Proof.* For any monotonically increasing function $\alpha(x) \geq 0$ and general function $\beta(x) \geq 0$, we can see that:

$$\int_0^t \alpha(x)\beta(x)dx \geq \int_0^t \alpha(0)\beta(x)dx = \alpha(0)\int_0^t \beta(x)dx\,. \tag{S44}$$

So, in the same way we can show that:

$$r(n+1,n+1,t) = \int_0^t f^\circ(t-x)^{n+1}p_A(x)dx \geq f^\circ(t)\int_0^t f^\circ(t-x)^n p_A(x)dx = f^\circ(t)\; r(n,n,t) \tag{S45}$$

since $f^\circ(t-x)$ is positive and monotonically increasing with $x$. We can equivalently show that:

$$\begin{aligned} r(0,n+1,t) &= \int_0^t (1 - f^\circ(t-x))^{n+1}p_A(x)dx \\ &\leq (1 - f^\circ(t))\int_0^t (1 - f^\circ(t-x))^n p_A(x)dx = (1 - f^\circ(t))\; r(0,n,t) \end{aligned} \tag{S46}$$

noting that here $1 - f^\circ(t-x)$ is **decreasing** with $x$ and therefore we have a $\leq$ sign instead of $\geq$.

So, we have proved that both of these are true:

$$\frac{r(n,n,t)}{r(n+1,n+1,t)} \leq \frac{1}{f^\circ(t)} \tag{S47}$$

$$\frac{r(0,n+1,t)}{r(0,n,t)} \leq 1 - f^\circ(t) \tag{S48}$$

Combining the last two inequalities by taking their product we get:

$$\forall t < t_{1/2} \qquad \frac{r(0,n+1,t)}{r(0,n,t)} \cdot \frac{r(n,n,t)}{r(n+1,n+1,t)} \leq \frac{1 - f^\circ(t)}{f^\circ(t)} < 1 \tag{S49}$$

where the last inequality is due to the fact that $f$ is monotonically decreasing, and therefore

$$\forall t < t_{1/2}\;\; : \;\; f^\circ(t) > f^\circ(t_{1/2}) = 1/2.$$

By moving some of the terms, we can see that $\frac{r(0,n,t)}{r(n,n,t)} > \frac{r(0,n+1,t)}{r(n+1,n+1,t)}$. So, now we can finally show that:

$$\forall t < t_{1/2} \qquad h(n,t) = \frac{1}{1 + \frac{r(0,n,t)}{r(n,n,t)}} < \frac{1}{1 + \frac{r(0,n+1,t)}{r(n+1,n+1,t)}} = h(n+1,t) \tag{S50}$$

■

The conclusion of Lemma S1.2 is that species with a higher multiplicity of labeling have higher labeling values compared to those with a lower multiplicity during the initial wash-out period before the input half-life, $t_{1/2}$. This property can be used in practice to detect delayed input, i.e., by comparing the labeling of metabolites with different labeling multiplicities at the beginning of wash-out.

##### S1.5.3 Identifiability of dynamic parameters in dynamic systems with delayed input

Just observing the labeling dynamics in systems with delayed input is insufficient to determine their dynamics parameters. To demonstrate this, we will use the example of a system fed by a well-mixed input source introduced in the section 1.5. We can note that the convolution operation that defines the observed labeling dynamics 16 is commutative:

$$\begin{aligned} f(t) &= f^{\circ}(t) - (\dot{f}^{\circ} * f_r)(t) \\ &= f_r(t) - (\dot{f}_r * f^{\circ})(t)\,. \end{aligned} \tag{S51}$$

Therefore, the metabolic age distribution $\mathcal{A}$ (and hence $f(t)$) would be identical if we exchange the input and reduced age distributions $\mathcal{A}^{\circ}$ $\mathcal{A}_r$ making it impossible to distinguish the two alternatives without additional information (see Appendix S1.7.2 for an example).

#### S1.6 Compartmental models

For the purpose of formal description of metabolic systems, the interconversion of metabolites in a cellular metabolic pathway can be seen as a type of directed graph, known as *compartmental models*, whose nodes represent distinct metabolite pools (e.g., pools of reaction intermediates or physically separated compartments such as membrane-delimited organelles) and the interconnecting edges - relevant metabolic reactions, (see Figure S5A and B). Such representations are useful for defining the content of labeled/unlabeled particles supplied through the corresponding metabolic routes known as atom mapping networks Ravikirthi et al. [2011], where every atom is mapped between every reactant (source pool) and every reaction product (recipient pool). An example is the interconversion of carbon $C_3$ in the pool of glucose-6-phosphate to the pool of carbon $C_3$ in fructose-6-phosphate that occurs through the phosphoglucoseisomerase reaction (see Figure S5C).

We will use compartmental models to mathematically describe steady-state metabolic age distributions of particles within a metabolic network and show their equivalence to the labeling dynamics in dynamic labeling experiments. The compartmental models are treated as mathematical objects consisting of particle pools and their interconnecting fluxes, which describe systems such as atomic mapping networks in the most general way. In compartmental models, the pools are assigned sizes $s_i$ (e.g., representing metabolite concentrations in each pool), while the connecting fluxes are assigned flux rates $v_{ji}$ (the flux from $S_j$ to $S_i$) and represent the transfer rates between each two pools (Figure S5D). In dynamic steady state, all the fluxes and pool sizes are constant, and so are the age distributions. In addition, elements in each state $S_i$ are considered to have a certain *labeled* fraction $f_i(t)$, i.e. the fraction of the original (labeled) metabolite that has not yet been replaced by time $t$ following the beginning of the wash-out. This fraction can be anywhere in the range between 0 and 1 since we can ignore the discrete nature of the molecules compared to the large size of the analyzed pools.

Since all living organisms are open systems, the pools in the compartmental models that describe them must also have *external* fluxes that exchange material with the external environment. The outside pools can be practically considered much larger than the cellular pools, and therefore we can ignore the effect of the system's effluxes on the environment (i.e., effluxes are marked by $\emptyset$ – see Figure S5D). We denote the time-dependent labeled fraction of the external pool by $h_e(t)$, the irreversible influx from the external pool to $S_i$ by $v_{ei}$, and the efflux as $v_{i\emptyset}$. Note that for simplicity, we assume that there is only one external pool ($E_1$).

##### S1.6.1 Metabolic age distributions in compartmental models

We will start by mathematically describing the relationships between steady-state metabolic age distribution of an externally supplied particle within the pools of a metabolic system represented by a compartmental model. Consider a general network with $n$ states (whose pool sizes are $\mathbf{s} \in \mathbb{R}^n$) and internal reactions described by the flux-weighted connectivity matrix $\mathbf{V} \in \mathbb{R}^{n\times n}$. Each element in the matrix, $v_{ji}$, represents the unidirectional flux from $S_j$ to $S_i$. In addition, we include a set of input reactions (with fluxes $\mathbf{v}_e \in \mathbb{R}^n$) that add new material (of zero age) from an external state into each one of the states, i.e. $v_{ei}$ is the flux of new material entering $S_i$. Contributed turnovers are defined as $\kappa_{ji} = v_{ji}/s_i$ and $\kappa_{ei} = v_{ei}/s_i$, and are arranged in the matrix $\mathbf{K}$ for the inner states and in the vector $\mathbf{k}_e$ for the external inputs.

We assume that the system has reached a steady-state, both in terms of the abundances and the ages of the elements in each state. Let $p_{\mathcal{A}_i}(t)$ denote the steady-state age distribution of elements in $S_i$. We will focus on the group of elements in $S_i$ whose age is within a specific age interval $[t, t+dt]$. The size of this group is $P(t \le \mathcal{A}_i < t+dt)\cdot s_i$. We can also write the mass-balance equation for this group, by considering four transitions affecting it during a single time step $dt$: (i) elements from $S_i$ of that were aged between $t-dt$ and $t$ enter the interval; (ii) all the elements from the interval age enough to leave it; (iii) a flux of new elements at the right age that enter $S_i$ from $S_j$ (for each of the other states including external ones); and (iv) a flux of elements at the right age leave $S_i$ to $S_j$ (or leave the system completely) and therefore never reach the interval (thus subtracting from the the first group). Since we are at steady-state, all these four effects must cancel out:

$$
\begin{aligned}
0 = \quad & \underbrace{P(t-dt \leq \mathcal{A}_i < t)\cdot s_i}_{\text{(i)}} \; - \; \underbrace{P(t \leq \mathcal{A}_i < t+dt)\cdot s_i}_{\text{(ii)}} \\
& + \; \underbrace{\sum_j P(t-dt \leq \mathcal{A}_j < t)\cdot v_{ji}}_{\text{(iii)}} \; - \; \underbrace{P(t-dt \leq \mathcal{A}_i < t)\cdot \left(\sum_j v_{ij} + v_{i\emptyset}\right)}_{\text{(iv)}} .
\end{aligned}
\tag{S52}
$$

Now, at the limit of $dt \to 0$, the first two terms can be approximated by $-\dot{p}_{\mathcal{A}_i}(t)dt \cdot s_i$ and in the latter two terms we can replace by $P(t-dt \leq \mathcal{A}_j < t)$ with $p_{\mathcal{A}_j}(t)dt$:

$$
\begin{aligned}
0 = & -\dot{p}_{\mathcal{A}_i}(t)\cancel{dt}\cdot s_i \; + \; \sum_j p_{\mathcal{A}_j}(t)\cancel{dt}\cdot v_{ji} \; - \; p_{\mathcal{A}_i}(t)\cancel{dt}\cdot \left(\sum_j v_{ij} + v_{i\emptyset}\right) \\
\dot{p}_{\mathcal{A}_i}(t) \; = & \; \sum_j p_{\mathcal{A}_j}(t)\cdot v_{ji}/s_i \; - \; p_{\mathcal{A}_i}(t)\cdot\left(\sum_j v_{ji} + v_{ei}\right)\Big/ s_i \\
\dot{p}_{\mathcal{A}_i}(t) \; = & \; \sum_j p_{\mathcal{A}_j}(t)\cdot \kappa_{ji} \; - \; p_{\mathcal{A}_i}(t)\cdot\left(\sum_j \kappa_{ji} + \kappa_{ei}\right)
\end{aligned}
\tag{S53}
$$

where in the second step we replace the sum of all outgoing fluxes, $\sum_j v_{ij} + v_{i\to\emptyset}$, with the sum of all incoming fluxes, $\sum_j vji + v_{ei}$, as they are equal due to the steady-state assumption.

This ODE can also be written in matrix notation, where we use the convention that bold lowercase letters represent column vectors and bold uppercase letters represent matrices. We denote $\mathbf{p}(\mathcal{A}=t)$ as the vector of all age probability density functions at time $t$, i.e., such that the $i$th element of the vector would be $p_{\mathcal{A}_i}(t)$. In addition, the vector $\mathbf{1}_n$ will be a column vector of ones and $\mathrm{diag}(\mathbf{x})$ is a diagonal matrix whose diagonal values are the elements of $\mathbf{x}$. Finally, by defining the state transition matrix $\mathbf{M} \equiv \left[\mathbf{K}^\top - \mathrm{diag}\left(\mathbf{K}^\top\mathbf{1}_n + \mathbf{k}_e\right)\right]$, we can now rewrite Equation S53 in matrix notation:

$$
\dot{\mathbf{p}}_{\mathcal{A}}(t) \;\; = \;\; \mathbf{M}\,\mathbf{p}_{\mathcal{A}}(t)\,. \tag{S54}
$$

We can thus see that the age distributions of states in a compartmental model are described by a system of first-order linear homogeneous ODEs. The general solution in this case is:

$$
\mathbf{p}_{\mathcal{A}}(t) \;\; = \;\; \mathrm{e}^{\mathbf{M}t}\,\mathbf{p}_{\mathcal{A}}(0) \tag{S55}
$$

and we can apply the boundary condition derived from the fact that $\mathbf{p}_{\mathcal{A}}(t)$ are PDF functions and therefore their integral must be equal to 1:

$$
\begin{aligned}
\mathbf{1}_n = \int_0^\infty \mathbf{p}_{\mathcal{A}}(t)df = \int_0^\infty \mathrm{e}^{\mathbf{M}t}df\cdot\mathbf{p}_{\mathcal{A}}(0) = \mathbf{M}^{-1}\mathrm{e}^{\mathbf{M}t}\Big|_0^\infty\cdot\mathbf{p}_{\mathcal{A}}(0) = \mathbf{M}^{-1}\left(-\mathbf{I}_n\right)\mathbf{p}_{\mathcal{A}}(0) \\
-\mathbf{M}\mathbf{1}_n = \mathbf{p}_{\mathcal{A}}(0)
\end{aligned}
\tag{S56}
$$

where we use the fact that $\lim_{t\to\infty}\mathrm{e}^{\mathbf{M}t} = \mathbf{0}$ (Lemma S1.21). Also, since $\mathbf{M}$ commutes with $\mathrm{e}^{\mathbf{M}t}$ we can write the solution as:

$$
\mathbf{p}_{\mathcal{A}}(t) \;\; = \;\; -\mathbf{M}\mathrm{e}^{\mathbf{M}t}\mathbf{1}_n \tag{S57}
$$

We can further derive an expression for the metabolic age CDF by integration:

$$
\mathbf{P}_{\mathcal{A}}(t) \;\; = \int_0^t \mathbf{p}_{\mathcal{A}}(\tau)d\tau = -\int_0^t \mathbf{M}\mathrm{e}^{\mathbf{M}\tau}d\tau\;\mathbf{1}_n = -\left(\mathrm{e}^{\mathbf{M}t} - \mathrm{e}^{\mathbf{M}0}\right)\mathbf{1}_n = \mathbf{1}_n - \mathrm{e}^{\mathbf{M}t}\mathbf{1}_n \tag{S58}
$$

##### S1.6.2 Compartmental models for dynamic labeling at metabolic steady-state

We now turn to modeling a wash-out experiment with an externally supplied metabolic label (e.g. $^{13}$C-glucose $\to$ $^{12}$C-glucose) using a CM. We aim to describe the time evolution of the labeled fractions of the internal metabolic pools, denoted

$$
f_i(t) \equiv s_i^{(1)}(t)/\left(s_i^{(1)}(t) + s_i^{(0)}(t)\right) = s_i^{(1)}(t)/s_i(t)\,. \tag{S59}
$$

where $s_i^{(1)}(t)$ is the amount that is labeled in pool $i$ at time $t$, $s_i^{(0)}(t)$ is the unlabeled amount, and $s_i(t)$ is the total.

The label in the internal pools will gradually be washed out, starting at $t = 0$. The rate of replacement (labeling) will be dictated by first-order kinetics, i.e., it will be proportional to the net balance of the influx of labeled atoms of the metabolite via connected internal and external pools and their loss through the efflux routes. Using the above notation, we can formally describe the change in the amount of labeled metabolite in each pool as:

$$\begin{aligned} s_i \dot{f}_i = \dot{s}_i^{(1)} &= \sum_{j=1}^{n} f_j v_{ji} + h_e v_{ei} - \sum_{j=1}^{n} f_i v_{ij} - f_i v_{i\to\emptyset} \\ &= \sum_{j=1}^{n} f_j v_{ji} - f_i \left( \sum_{j=1}^{n} v_{ij} + v_{i\emptyset} \right) + h_e v_{ei} \\ &= \sum_{j=1}^{n} f_j v_{ji} - f_i \left( \sum_{j=1}^{n} v_{ji} + v_{ei} \right) + h_e v_{ei} \end{aligned} \tag{S60}$$

where we replace the sum of all the outgoing fluxes, $\sum_j v_{ij} + v_{i\emptyset}$, with the sum of all incoming fluxes, $\sum_j v_{ji} + v_{ei}$, as they are equal due to the steady-state assumption. By divining both sides by $s_i$ we can also replace the fluxes with the contributed turnover rates: $\kappa_{ji} = v_{ji}/s_i$ and $\kappa_{ei} = v_{ei}/s_i$. Note that these rates are all constant due to the steady-state assumption.

$$\dot{f}_i = \sum_{j=1}^{n} f_j \kappa_{ji} - f_i \left( \sum_{j=1}^{n} \kappa_{ji} + \kappa_{ei} \right) + h_e \kappa_{ei} \tag{S61}$$

This ODE can also be written in matrix notation, where we use the convention that bold lowercase letters represent column vectors, and bold uppercase letters represent matrices. In addition, the matrix $\mathbf{K}$ and the vector $\mathbf{k}_e$ contain the contributed turnovers of internal and external fluxes (whose elements are $\kappa_{ji}$ and $\kappa_{ei}$, respectively). The vector $\mathbf{1}_n$ will be a column vector of ones and $\text{diag}(\mathbf{x})$ is a diagonal matrix whose diagonal values are the elements of $\mathbf{x}$.

$$\dot{\mathbf{f}} = \left[\mathbf{K}^\top - \text{diag}\left(\mathbf{K}^\top \mathbf{1}_n + \mathbf{k}_e\right)\right] \mathbf{f} + h_e \mathbf{k}_e \tag{S62}$$

which, if we define $\mathbf{M} \equiv \left[\mathbf{K}^\top - \text{diag}\left(\mathbf{K}^\top \mathbf{1}_n + \mathbf{k}_e\right)\right]$ and $\mathbf{g} \equiv h_e \mathbf{k}_e$, turns into (see Appendix S1.11.1):

$$\dot{\mathbf{f}} = \mathbf{M}\mathbf{f} + \mathbf{g} \tag{S63}$$

Therefore, the labeling kinetics in the dynamic labeling experiments in the most general case can be described via first-order linear inhomogeneous ODEs.

The general solution Young et al. [2008] for the inhomogeneous first order linear ODE in Equation S63 is:

$$\mathbf{f}(t_1) = \ \mathrm{e}^{\mathbf{M}(t_1 - t_0)} \mathbf{f}(t_0) + \int_{t_0}^{t_1} \mathrm{e}^{\mathbf{M}(t_1 - \tau)} \mathbf{g}(\tau) d\tau \tag{S64}$$

Without loss of generality, we can set $t_0 = 0$ and define $\mathbf{f}^\circ \equiv \mathbf{f}(0)$. Further assuming that labeling of the input is constant (denoted $h_e^\circ$) and therefore so is $\mathbf{g}$ (for the more general case, see Section S1.9) we can re-write the solution of the ODEs in a closed form as:

$$\mathbf{f}(t) = \mathrm{e}^{\mathbf{M}t} \mathbf{f}^\circ + \int_0^t \mathrm{e}^{\mathbf{M}(t-\tau)} \mathbf{g} d\tau = \mathrm{e}^{\mathbf{M}t} \mathbf{f}^\circ + \int_0^t \mathrm{e}^{\mathbf{M}(t-\tau)} d\tau \cdot \mathbf{g} = \mathrm{e}^{\mathbf{M}t} \mathbf{f}^\circ - \mathrm{e}^{\mathbf{M}t} \mathbf{M}^{-1} \mathbf{g} + \mathbf{M}^{-1} \mathbf{g}\,. \tag{S65}$$

The labeling of all internal pools must eventually reach a time-independent isotopic steady state. Indeed, since $\mathbf{M}$ has no positive eigenvalues, $\lim_{t\to\infty} \mathrm{e}^{\mathbf{M}t} = 0$ (see Appendix Lemma S1.21) we can observe that the labeling at infinitely late time points solely depends on external and internal fluxes and labeling of the external pools $\mathbf{f}^\infty \equiv \lim_{t\to\infty} \mathbf{f}(t) = \mathbf{M}^{-1}\mathbf{g} = \mathbf{M}^{-1}\mathbf{k}_e h_e^\circ$. Thus in terms of the initial and final labeling the dynamics of pool labeling can be represented as:

$$\mathbf{f}(t) = \mathrm{e}^{\mathbf{M}t} \mathbf{f}^\circ + \left(\mathbf{I}_n - \mathrm{e}^{\mathbf{M}t}\right) \mathbf{f}^\infty \tag{S66}$$

In most dynamic labeling experiments, the metabolites start as completely labeled and then the medium is switched to a completely unlabeled nutrient (or vice versa). In this case, we can assume that $\mathbf{f}^\circ = \mathbf{1}_n$ and $h_e^\circ = 0$ (and therefore $\mathbf{f}^\infty = \mathbf{0}_n$), so the labeling loss kinetic solution can be further simplified as:

$$\mathbf{f}(t) = \mathrm{e}^{\mathbf{M}t} \mathbf{1}_n\,. \tag{S67}$$

Therefore, the **M**-matrix in this form can be viewed as an array of contributed turnovers resulting from material transfer from source pools, specified by the column indices, to recipient pools, specified by the row indices, and where the diagonal elements denote the bulk fractional efflux from the recipient pools (Figure S5 E). Furthermore, the dynamics of labeling in each pool (see Equation S63 and consider $\mathbf{g} = \mathbf{0_n}$) can be seen as a balance of fractional gain of the labeled metabolite trough contributed turnovers from other pools and fractional loss through bulk efflux to internal pools and external environment (Figure S5 D).

The **M**-matrix and the pool sizes must also satisfy conditions of mass balance that we refer to as the *mass-balance constraint* i.e. that there is a non-negative efflux to the outside environment from every pool in the system. In the case where the system is growing, the efflux must be large enough to at least support the system growth (Figure S5F). We can express the mass-balance constraint using the **M**-matrix and pools sizes notation as (same as Equation 24 in the main text):

$$\mathbf{M}^{\top}\mathbf{s} \leq -\mu\mathbf{s}\,. \tag{S68}$$

The left-hand side of this equation represents the bulk efflux from each of the pools to the external environment (with a negative sign), and the right-hand side corresponds to the minimum requirement to support the growth of the system at a rate $\mu$. Note that both sides are negative vectors, so the inequality ensures that the absolute efflux is *larger* than the growth requirement.

##### S1.6.3 Reduced compartmental models

Given the **M**-matrix and **s** of the whole system (containing both unobserved and observed subsystems), the *reduced compartmental model*, denoted by $\mathbf{M}_r$ and $\mathbf{s}_r$, is defined by removing all rows and columns of the **M**-matrix except the ones that contain the observed pools and renormalizing the sizes of the observed pools $\mathbf{s}_r$ (Figure S6). Note that in reduced form, all the contributed turnovers of the observed subsystem remain unchanged, while all internal feeding fluxes become external. Reduced CMs would thus describe a *reduced* form of the observed subsystem, where labeling enters directly without delay. All dynamic characteristics of the reduced subsystem could be represented by reduced CMs and can be expressed analogously to the full CMs (see Tables 1 and S1). For example, the mean age of the reduced system would be:

$$\bar{\mathcal{A}}_r = -\mathbf{s}_r\mathbf{M}_r^{-1}\mathbf{1}_n\,. \tag{S69}$$

This approach can be practical for compensating for the contribution of precursor pools to the labeling dynamics, a confounding effect common in most of the dynamic labeling experiments and difficult to eliminate experimentally.

##### S1.6.4 General properties of labeling dynamics in compartmental models

Although the observed labeling dynamics of individual metabolite pools is dependent in a complex way on the other pool sizes and the connecting flux routes, it has some universal properties. First, we note that the labeling of any metabolite pool in any steady-state dynamic labeling assay must only decrease during the labeling pulse. Indeed, since **M** has no positive eigenvalues, the first derivatives of **f** must also be negative (see the Appendix Corollary S1.20.1).

$$\dot{\mathbf{f}} = \mathbf{M}\mathrm{e}^{\mathbf{M}t}\mathbf{1}_n \leq \mathbf{0}_n \tag{S70}$$

This makes intuitive sense since switching to a completely unlabeled medium would gradually eliminate the labeled metabolites from all of the internal pools.

For such experiments we can also define the *system labeled fraction* as the weighted average of labeled fractions among all internal pools, namely:

$$f_{\mathbf{s}} \equiv \frac{\sum_{i=1}^{n} f_i\, s_i}{\sum_{i=1}^{n} s_i} = \frac{\mathbf{s}^{\top}\mathbf{f}}{\mathbf{s}^{\top}\mathbf{1}_n}\,. \tag{S71}$$

$f_{\mathbf{s}}$ is useful in experiments in which the system's labeled fraction can be directly measured. For example, when the $^{13}$C fraction of all biomass carbons can be measured by complete oxidation of dry biomass to carbon dioxide or when bulk protein labeling can be measured using isotopically labeled amino acids.

To find $f_{\mathbf{s}}$ we can use the solution from Equation S67 and write an explicit expression for $f_{\mathbf{s}}$ based on the **M** and **s**:

$$f_{\mathbf{s}} = \frac{\mathbf{s}^{\top}\mathrm{e}^{\mathbf{M}t}\mathbf{1}_n}{\mathbf{s}^{\top}\mathbf{1}_n}\,. \tag{S72}$$

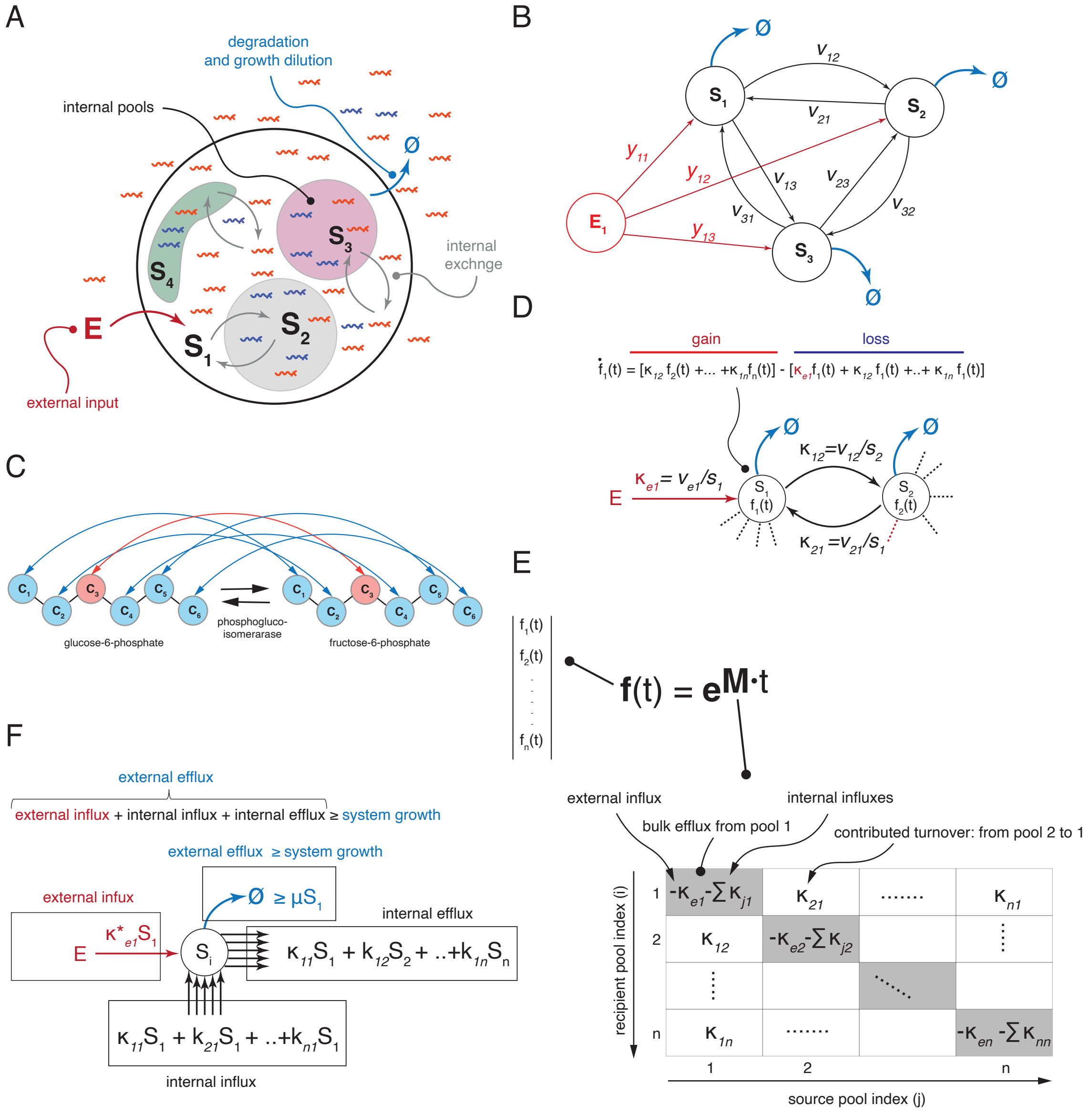


Figure S5: **Formal description of labeling dynamics using compartmental models.** (A) A cellular metabolic pathway represented by a *compartmental model* i.e. as a set of well-mixed pools illustrating physical compartments (e.g. cytosol, mitochondrion, vacuole, nuclear interior etc...) or metabolic states that exchange a labeled metabolite (blue) with an externally added unlabeled one (red). The exchange occurs due to (i) internal exchange fluxes between the internal pools (gray arrows), (ii) the external influx(es) of the labeled metabolite from the external environment (red arrow), and (iii) the effluxes to the outside environment (blue arrows). (B) Formal representation of a compartmental model wherein the internal pools and connecting fluxes are assigned certain sizes $S_i$ and flux rates $v_{ij}$, respectively. (C) Illustration of atomic-mapping network – a physical model of material transfer between metabolic pools. As an exemplary case of interconversion between glucose-6-phosphate and fructose-6-phosphate in a phosphoglucoisomerase-catalyzed reaction the carbon $C_3$ of glucose-6-phosphate will always be converted (will maps to) carbon $C_3$ of fructose-6-phosphate. Similar concept is applicable to larger units e.g. chemical groups or even macromolecules as long as they can be treated as unsplittable (atomic) units withing the compartmental model. (D) View of the fractional labeling ($f$) dynamics in the internal pools as a balance between factional gain and fractional loss rates of a labeled metabolite through the connecting fluxes, that can be determined using *contributed turnovers* ($k_{ij}$) by the incoming fluxes and the current labeling in the respective pools. (E) Construction of **M**-matrix, governing time-evolution of labeling dynamics. The off-diagonal elements of **M**-matrix represent fractional exchange rates of recipient pools contributed by the incoming fluxes from the source pools defined by the (*contributed turnovers*). The diagonal elements represent bulk turnover rates of the recipient pools (with a negative sign) contributed by the sum of all incoming fluxes (contributed turnovers) from the internal pools and the eternal environment altogether. (F) Illustration for the expression for mass-balance constraint (Equation 24) where efflux of material to the outside environment must be at least as large as needed to support the system's growth and can be conveniently computed using pools sizes and and contributed turnovers arrayed within the **M**-matrix.

375 If we further use the fact that $\mathbf{s}$ and $\mathbf{f}$ are non-negative vectors and so is $-\mathbf{s}^\top \mathbf{M}$ as it contains the output exchange

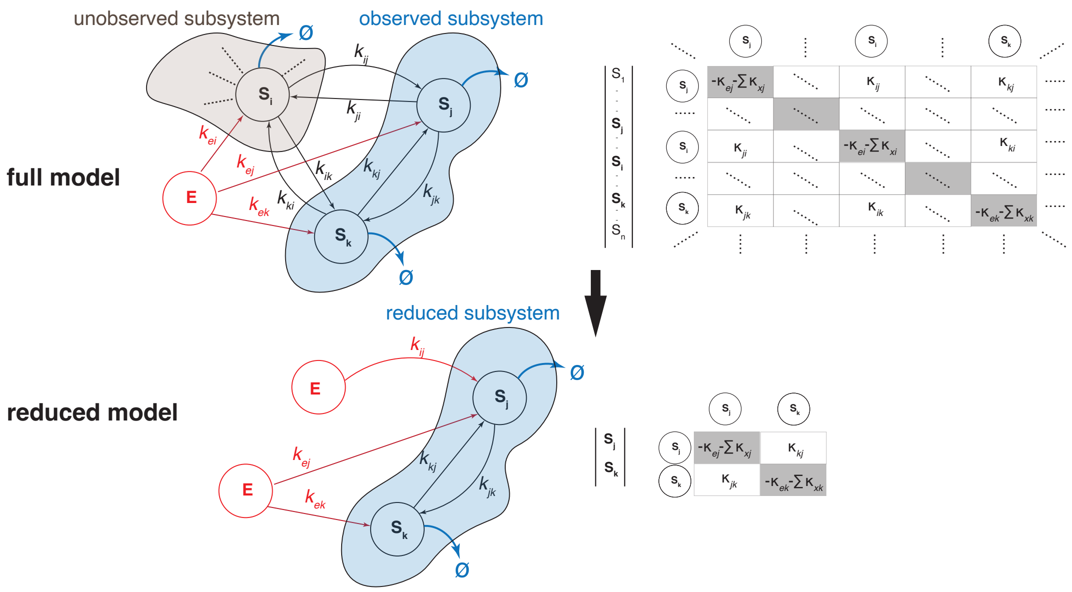


Figure S6: **Reduced compartmental models.** Example of a reduced compartmental model retaining only pools $S_j$ and $S_k$ of the observed subsystem and their connecting fluxes, which is constructed by changing all the internal incoming and outgoing fluxes for these pools to external while retaining their magnitudes. Dynamics of such *reduced subsystem* can be described using a reduced $\mathbf{M}_r$- matrix retaining only rows and columns connecting the observed pools and the re-normalized the observed pool-size vector $\mathbf{s}_r = (s_j, s_k)$. In this example the reduced model would mimic the dynamics of the observed two-pool subsystem as if it would receive particles directly from external environment instead of the ones it receives from the unobserved pools.

fluxes (see Appendix S1.11.2), we can see that the first derivative satisfies:

$$\dot{f}_{\mathbf{s}} = \frac{\mathbf{s}^\top \dot{\mathbf{f}}}{\mathbf{s}^\top \mathbf{1}_n} = -\frac{(-\mathbf{s}^\top \mathbf{M})\mathbf{f}}{\mathbf{s}^\top \mathbf{1}_n} \leq \mathbf{0}_n \tag{S73}$$

and likewise for the second derivative:

$$\ddot{f}_{\mathbf{s}} = \frac{\mathbf{s}^\top \mathbf{M}^2 \mathrm{e}^{\mathbf{M}t} \mathbf{1}_n}{\mathbf{s}^\top \mathbf{1}_n} = \frac{(\mathbf{s}^\top \mathbf{M})(\mathbf{M}\mathrm{e}^{\mathbf{M}t} \mathbf{1}_n)}{\mathbf{s}^\top \mathbf{1}_n} = \frac{(-\mathbf{s}^\top \mathbf{M})\ (-\dot{\mathbf{f}})}{\mathbf{s}^\top \mathbf{1}_n} \geq \mathbf{0}_n \tag{S74}$$

Therefore, the labeling dynamics of a cellular metabolite measured in bulk (labeling of all pools weighted by pool size) is a monotonously decreasing (Equation S73) and convex (Equation S74) function. This can be understood intuitively – the gradual washout of the labeled metabolite leads to a progressively less labeled efflux, which in turn slows down the washout of labeled metabolite.

What is less intuitive is that, as long as our system does not have completely isolated pools, i.e. all the pools are directly or indirectly connected to the outside environment, the labeling of each pool individually will exponentially decay to zero at sufficiently long time-scales (see Lemma S1.21), i.e.:

$$\forall i \;\; \exists(\alpha_i, \beta_i, t_i) \quad \text{such that} \quad \forall t > t_i \;\;\; f_i(t) \leq \beta_i \mathrm{e}^{-\alpha_i t}\,. \tag{S75}$$

This means that in practice we can basically ignore the labeling dynamics beyond a certain point, as the decay is very fast (compared, say, to any polynomial). For example, when estimating mean ages (see Appendix S1.3.1), we do not need to continue measuring $f_i$ for prolonged periods of time, but rather can assume it continues to decay exponentially after a certain point.

Together, we can conclude that in steady-state dynamic labeling experiments the labeling of metabolites must follow certain rules. The labeled fraction in every pool must always decrease over time. In the case of systems with direct input, the *rate* of change in the labeling fraction must also decrease over time. In addition, the exponential decay property of the labeling dynamics allows simple interpretations of cumulative dynamic parameters such as mean metabolic ages or residence times as geometric properties of labeling dynamics (labeling curve), as described in Sections 1.2-1.3.

##### S1.6.5 Dynamic parameters of compartmental systems

To find out how the dynamic characteristics of the system are shaped by the underlying CMs, we will use Equations 2-7 that express them in terms of the labeling dynamics, $\mathbf{f}(t)$, and then replace it with the explicit function for CMs according to Equation 20.

In this way, the metabolic age distributions can be determined for each pool within the CM by combining Equations 2 with Equation 20:

$$\begin{aligned}\mathbf{P}_{\mathcal{A}}(t) &= \mathbf{1}_n - \mathbf{f}(t) = \mathbf{1}_n - \mathrm{e}^{\mathbf{M}t}\mathbf{1}_n \\ \mathbf{p}_{\mathcal{A}}(t) &= -\dot{\mathbf{f}}(t) = -\frac{d\left(\mathrm{e}^{\mathbf{M}t}\mathbf{1}_n\right)}{dt} = -\mathbf{M}\mathrm{e}^{\mathbf{M}t}\mathbf{1}_n\end{aligned} \tag{S76}$$

To determine age distributions for subsets of pools we can use the size-weighted averages of the constituent pools. For example, assuming that the pool sizes of the whole system sum up to 1 (i.e., normalized) we can determine the *system age* CDF and PDF as:

$$\begin{aligned}P_{\mathcal{A}_\mathbf{s}}(t) &= \mathbf{s}^\top\mathbf{P}_{\mathcal{A}}(t) = \mathbf{s}^\top\left(\mathbf{1}_n - \mathrm{e}^{\mathbf{M}t}\mathbf{1}_n\right) = 1 - \mathbf{s}^\top\mathrm{e}^{\mathbf{M}t}\mathbf{1}_n = 1 - f_\mathbf{s}(t) \\ p_{\mathcal{A}_\mathbf{s}}(t) &= \dot{f}_\mathbf{s}(t) = -\frac{d\left(\mathbf{s}^\top\mathrm{e}^{\mathbf{M}t}\mathbf{1}_n\right)}{dt} = -\mathbf{s}^\top\mathbf{M}\mathrm{e}^{\mathbf{M}t}\mathbf{1}_n\end{aligned} \tag{S77}$$

We can further use the fact that system age CDF in the Equation S77 is connected with the whole system's labeling dynamics $f_\mathbf{s}(t)$ the same way as the age CDF is connected with labeling dynamics $f(t)$ in the case of general dynamic systems (Equation 2). This allows us to directly use the expressions for the dynamic parameters as a function of labeling derived for the general systems.

Equation 4 can be used to determine the mean ages in a compartment-specific way or for the whole system:

$$\begin{aligned}\bar{\mathcal{A}} &= \int_0^\infty \mathbf{f}(t)dt = \int_0^\infty \mathrm{e}^{\mathbf{M}t}\mathbf{1}_n dt = -\mathbf{M}^{-1}\mathbf{1}_n \\ \bar{\mathcal{A}}_\mathbf{s} &= \int_0^\infty f_\mathbf{s}(t)dt = \int_0^\infty \mathbf{s}^\top\mathrm{e}^{\mathbf{M}t}\mathbf{1}_n dt = -\mathbf{s}^\top\mathbf{M}^{-1}\mathbf{1}_n\end{aligned} \tag{S78}$$

Residence time distributions and decay rates are only defined for zero-input-delay systems. Since the CM as a whole satisfies this condition, we can express them using Equations 5-7 and the expression for the system labeling dynamics:

$$\begin{aligned}P_{\mathcal{T}}(t) &= 1 - \frac{\dot{f}_\mathbf{s}(t)}{\dot{f}_\mathbf{s}(0)} = 1 - \frac{\mathbf{s}^\top\mathbf{M}\mathrm{e}^{\mathbf{M}t}\mathbf{1}_n}{\mathbf{s}^\top\mathbf{M}\mathbf{1}_n} \\ p_{\mathcal{T}}(t) &= -\frac{\ddot{f}_\mathbf{s}(t)}{\dot{f}_\mathbf{s}(0)} = -\frac{1}{\mathbf{s}^\top\mathbf{M}\mathbf{1}_n}\frac{d\left(\mathbf{s}^\top\mathbf{M}\mathrm{e}^{\mathbf{M}t}\mathbf{1}_n\right)}{dt} = -\frac{\mathbf{s}^\top\mathbf{M}^2\mathrm{e}^{\mathbf{M}t}\mathbf{1}_n}{\mathbf{s}^\top\mathbf{M}\mathbf{1}_n} \\ \kappa(\mathcal{A}_\mathbf{s}) &= -\frac{\ddot{f}_\mathbf{s}(t)}{\dot{f}_\mathbf{s}(t)} = -\frac{\mathbf{s}^\top\mathbf{M}^2\mathrm{e}^{\mathbf{M}t}\mathbf{1}_n}{\mathbf{s}^\top\mathbf{M}\mathrm{e}^{\mathbf{M}t}\mathbf{1}_n}.\end{aligned} \tag{S79}$$

Furthermore, we can determine the expected values for residence times and decay rates using the corresponding results for general systems (Equation 9):

$$\bar{\kappa} = \bar{\mathcal{T}}^{-1} = -\dot{f}_\mathbf{s}(0) = \mathbf{s}^\top\mathbf{M}\mathbf{1}_n \tag{S80}$$

and the age-cohort mean residence time:

$$\mathcal{T}_a = \mathbb{E}[\mathcal{T} \mid \mathcal{T} \geq a] = a - \frac{f_\mathbf{s}(a)}{\dot{f}_\mathbf{s}(a)} = a - \frac{\mathbf{s}^\top\mathrm{e}^{\mathbf{M}a}\mathbf{1}_n}{\mathbf{s}^\top\mathbf{M}\mathrm{e}^{\mathbf{M}a}\mathbf{1}_n} \tag{S81}$$

Similarly, we can use the same procedures for expressing the dynamic parameters for the growing systems using Equations 12-14:

$$
\begin{aligned}
P_{\mathcal{T}}(t) &= 1 - \mathrm{e}^{\mu t}\ \frac{\dot{f}_{\mathbf{s}}(t)}{\dot{f}_{\mathbf{s}}(0)} = 1 - \mathrm{e}^{\mu t}\ \frac{\mathbf{s}^\top \mathbf{M} \mathrm{e}^{\mathbf{M}t} \mathbf{1}_n}{\mathbf{s}^\top \mathbf{M} \mathbf{1}_n} \\
p_{\mathcal{T}}(t) &= -\mathrm{e}^{\mu t}\ \frac{\ddot{f}_{\mathbf{s}}(t) + \mu \dot{f}_{\mathbf{s}}(t)}{\dot{f}_{\mathbf{s}}(0)} = -\mathrm{e}^{\mu t} \frac{\mathbf{s}^\top \mathbf{M}^2 \mathrm{e}^{\mathbf{M}t} \mathbf{1}_n + \mu \mathbf{s}^\top \mathbf{M} \mathrm{e}^{\mathbf{M}t} \mathbf{1}_n}{\mathbf{s}^\top \mathbf{M} \mathbf{1}_n} \\
\kappa(\mathcal{A}_{\mathbf{s}}) = \lambda(\mathcal{A}_{\mathbf{s}}) - \mu &= -\frac{\ddot{f}_{\mathbf{s}}(t)}{\dot{f}_{\mathbf{s}}(t)} - \mu = -\frac{\mathbf{s}^\top \mathbf{M}^2 \mathrm{e}^{\mathbf{M}t} \mathbf{1}_n}{\mathbf{s}^\top \mathbf{M} \mathrm{e}^{\mathbf{M}t} \mathbf{1}_n} - \mu \\
\bar{\mathcal{T}} &= \frac{1}{\dot{f}(0)} \int_0^\infty \dot{f}(t)\ \mathrm{e}^{\mu t}\ dt = \frac{\mathbf{s}^\top \mathbf{M} \left( \int_0^\infty \mathrm{e}^{(\mathbf{M} + \mu \mathbf{I}_n)t} dt \right) \mathbf{1}_n}{\mathbf{s}^\top \mathbf{M} \mathbf{1}_n} = -\frac{\mathbf{s}^\top \mathbf{M} (\mathbf{M} + \mu \mathbf{I}_n)^{-1} \mathbf{1}_n}{\mathbf{s}^\top \mathbf{M} \mathbf{1}_n} \\
\bar{\kappa} &= -\dot{f}(0) - \mu = -\mathbf{s}^\top \mathbf{M} \mathbf{1}_n - \mu \\
\bar{\mathcal{T}}_a &= a + \frac{\int_0^\infty \dot{f}_{\mathbf{s}}(t+a)\ \mathrm{e}^{\mu t}\ dt}{\dot{f}_{\mathbf{s}}(a)} = a + \frac{\mathbf{s}^\top \mathbf{M} \left( \int_0^\infty \mathrm{e}^{(\mathbf{M} + \mu \mathbf{I}_n)t} dt \right) \mathrm{e}^{\mathbf{M}a} \mathbf{1}_n}{\mathbf{s}^\top \mathbf{M} \mathrm{e}^{\mathbf{M}a} \mathbf{1}_n} = a - \frac{\mathbf{s}^\top \mathbf{M} (\mathbf{M} + \mu \mathbf{I}_n)^{-1} \mathrm{e}^{\mathbf{M}a} \mathbf{1}_n}{\mathbf{s}^\top \mathbf{M} \mathrm{e}^{\mathbf{M}a} \mathbf{1}_n} \\
\bar{\kappa}_a &= -\frac{\dot{f}(a)}{f(a)} - \mu = -\frac{\mathbf{s}^\top \mathbf{M} \mathrm{e}^{\mathbf{M}a} \mathbf{1}_n}{\mathbf{s}^\top \mathrm{e}^{\mathbf{M}a} \mathbf{1}_n} - \mu
\end{aligned}
\tag{S82}
$$

416 For quick reference, most of these results are summarized in Table S1.

| Description | unit | Symbol | Expression of $\mathcal{A}$ | Expression of $f$ | Expression of M and s | Eq. |
|---|---|---|---|---|---|---|
| labeled fractions | unitless | $\mathbf{f}(t)$ | $\mathbf{1}_n - \mathbf{P}(\mathcal{A} \le t)$ | $\mathbf{f}(t)$ | $e^{\mathbf{M}t}\mathbf{1}_n$ | 20 |
| ages CDF | unitless | $\mathbf{P}_\mathcal{A}(t)$ | $\mathbf{P}_\mathcal{A}(t)$ | $\mathbf{1}_n - \mathbf{f}(t)$ | $\mathbf{1}_n - e^{\mathbf{M}t}\mathbf{1}_n$ | S76 |
| ages PDF | time$^{-1}$ | $\mathbf{p}_\mathcal{A}(t)$ | $\mathbf{p}_\mathcal{A}(t)$ | $-\dot{\mathbf{f}}(t)$ | $-\mathbf{M}e^{\mathbf{M}t}\mathbf{1}_n$ | S76 |
| mean ages | time | $\bar{\mathcal{A}}$ | $\int_0^\infty t\ \mathbf{p}_\mathcal{A}(t)\ dt$ | $\int_0^\infty \mathbf{f}(t)dt$ | $-\mathbf{M}^{-1}\mathbf{1}_n$ | S78 |
| **Systems with direct input** | | | | | | |
| system labeled fraction | unitless | $f_\mathbf{s}(t)$ | $1 - P_{\mathcal{A}_\mathbf{s}}(t)$ | $\mathbf{s}^\top \mathbf{f}(t)$ | $\mathbf{s}^\top e^{\mathbf{M}t}\mathbf{1}_n$ | 21, S71 |
| system age CDF | unitless | $P_{\mathcal{A}_\mathbf{s}}(t)$ | $P_{\mathcal{A}_\mathbf{s}}(t)$ | $1 - f_\mathbf{s}(t)$ | $1 - \mathbf{s}^\top e^{\mathbf{M}t}\mathbf{1}_n$ | 23, S77 |
| system age PDF | time$^{-1}$ | $p_{\mathcal{A}_\mathbf{s}}(t)$ | $p_{\mathcal{A}_\mathbf{s}}(t)$ | $-\dot{f}_\mathbf{s}(t)$ | $-\mathbf{s}^\top \mathbf{M}e^{\mathbf{M}t}\mathbf{1}_n$ | S77 |
| median system age (half-life) | time | $\mathcal{A}_{\mathbf{s},\frac{1}{2}}$ | $P_{\mathcal{A}_\mathbf{s}}\left(\mathcal{A}_{\mathbf{s},\frac{1}{2}}\right) = \frac{1}{2}$ | $f_\mathbf{s}^{-1}\left(\frac{1}{2}\right)$ | $\mathbf{s}^\top e^{\mathbf{M}t_{1/2}}\mathbf{1}_n = \frac{1}{2}$ | |
| system residence time CDF | unitless | $P_\mathcal{T}(t)$ | $1 - \frac{p_{\mathcal{A}_\mathbf{s}}(t)}{p_{\mathcal{A}_\mathbf{s}}(0)}$ | $1 - \frac{\dot{f}_\mathbf{s}(t)}{\dot{f}_\mathbf{s}(0)}$ | $1 - \mathbf{s}^\top \mathbf{M}e^{\mathbf{M}t}\mathbf{1}_n \Big/ \mathbf{s}^\top \mathbf{M}\mathbf{1}_n$ | 5, S79 |
| system residence time PDF | time$^{-1}$ | $p_\mathcal{T}(t)$ | $-\frac{\dot{p}_{\mathcal{A}_\mathbf{s}}(t)}{p_{\mathcal{A}_\mathbf{s}}(0)}$ | $-\frac{\ddot{f}_\mathbf{s}(t)}{\dot{f}_\mathbf{s}(0)}$ | $-\mathbf{s}^\top \mathbf{M}^2e^{\mathbf{M}t}\mathbf{1}_n \Big/ \mathbf{s}^\top \mathbf{M}\mathbf{1}_n$ | 6, S79 |
| mean system residence time | time | $\bar{\mathcal{T}}$ | $-\frac{1}{p_{\mathcal{A}_\mathbf{s}}(0)}$ | $-\frac{1}{\dot{f}_\mathbf{s}(0)}$ | $-1 \Big/ \mathbf{s}^\top \mathbf{M}\mathbf{1}_n$ | 9, S80 |
| system decay rate | time$^{-1}$ | $\kappa(\mathcal{A}_\mathbf{s})$ | $-\frac{\dot{p}_{\mathcal{A}_\mathbf{s}}(t)}{p_{\mathcal{A}_\mathbf{s}}(t)}$ | $-\frac{\ddot{f}_\mathbf{s}(t)}{\dot{f}_\mathbf{s}(t)}$ | $-\mathbf{s}^\top \mathbf{M}^2e^{\mathbf{M}t}\mathbf{1}_n \Big/ \mathbf{s}^\top \mathbf{M}e^{\mathbf{M}t}\mathbf{1}_n$ | 7, S79 |
| expected system decay rate | time$^{-1}$ | $\bar{\kappa}$ | $p_{\mathcal{A}_\mathbf{s}}(0)$ | $-\dot{f}_\mathbf{s}(0)$ | $-\mathbf{s}^\top \mathbf{M}\mathbf{1}_n$ | 9, S14, S80 |
| **Growing systems (with growth rate $\mu$)** | | | | | | |
| system residence time CDF | unitless | $P_\mathcal{T}(t)$ | $1 - e^{\mu t}\frac{p_{\mathcal{A}_\mathbf{s}}(t)}{p_{\mathcal{A}_\mathbf{s}}(0)}$ | $1 - e^{\mu t}\frac{\dot{f}_\mathbf{s}(t)}{\dot{f}_\mathbf{s}(0)}$ | $1 - e^{\mu t}\ \mathbf{s}^\top \mathbf{M}e^{\mathbf{M}t}\mathbf{1}_n \Big/ \mathbf{s}^\top \mathbf{M}\mathbf{1}_n$ | 13, S82 |
| system residence time PDF | time$^{-1}$ | $p_\mathcal{T}(t)$ | $-e^{\mu t}\ \frac{\dot{p}_{\mathcal{A}_\mathbf{s}}(t)+\mu p_{\mathcal{A}_\mathbf{s}}(t)}{p_{\mathcal{A}_\mathbf{s}}(0)}$ | $-e^{\mu t}\ \frac{\ddot{f}_\mathbf{s}(t)+\mu \dot{f}_\mathbf{s}(t)}{\dot{f}_\mathbf{s}(0)}$ | $-e^{\mu t}\ \mathbf{s}^\top \mathbf{M}(\mathbf{M}+\mu\mathbf{I}_n)e^{\mathbf{M}t}\mathbf{1}_n \Big/ \mathbf{s}^\top \mathbf{M}\mathbf{1}_n$ | 14, S82 |
| mean system residence time | time | $\bar{\mathcal{T}}$ | $\int_0^\infty \frac{p_{\mathcal{A}_\mathbf{s}}(t)}{p_{\mathcal{A}_\mathbf{s}}(0)}\ e^{\mu t}\ dt$ | $\int_0^\infty \frac{\dot{f}_\mathbf{s}(t)}{\dot{f}_\mathbf{s}(0)}\ e^{\mu t}\ dt$ | $-\mathbf{s}^\top \mathbf{M}(\mathbf{M}+\mu\mathbf{I}_n)^{-1}\mathbf{1}_n \Big/ \mathbf{s}^\top \mathbf{M}\mathbf{1}_n$ | 9, S82 |
| residence time of the living CDF | unitless | $P_\mathcal{L}(t)$ | $P_{\mathcal{A}_\mathbf{s}}(t) - \frac{e^{\mu t}-1}{\mu}p_{\mathcal{A}_\mathbf{s}}(t)$ | $1 - f_\mathbf{s}(t) + \frac{e^{\mu t}-1}{\mu}\dot{f}_\mathbf{s}(t)$ | $1 - \frac{1}{\mu}\ \mathbf{s}^\top\left(\mathbf{M}_\mu e^{\mathbf{M}t} - \mathbf{M}e^{\mathbf{M}_\mu t}\right)\mathbf{1}_n$ | S29 |
| residence time of the living PDF | time$^{-1}$ | $p_\mathcal{L}(t)$ | $\frac{e^{\mu t}-1}{\mu}\left(\mu p_{\mathcal{A}_\mathbf{s}}(t) + t\dot{p}_{\mathcal{A}_\mathbf{s}}(t)\right)$ | $\frac{1-e^{\mu t}}{\mu}\left(\mu \dot{f}_\mathbf{s}(t) + \ddot{f}_\mathbf{s}(t)\right)$ | $-\frac{1}{\mu}\ \mathbf{s}^\top \mathbf{M}\ \mathbf{M}_\mu\left(e^{\mathbf{M}t} - e^{\mathbf{M}_\mu t}\right)\mathbf{1}_n$ | 15, S29 |
| mean residence time of the living | time | $\bar{\mathcal{L}}$ | $\int_0^\infty (1+e^{\mu t})[1-P_{\mathcal{A}_\mathbf{s}}(t)]\ dt$ | $\int_0^\infty (1+e^{\mu t})f_\mathbf{s}(t)\ dt$ | $-\mathbf{s}^\top(\mathbf{M}^{-1}+\mathbf{M}_\mu^{-1})\mathbf{1}_n$ | 15, S30 |
| system decay rate | time$^{-1}$ | $\kappa(\mathcal{A}_\mathbf{s})$ | $-\frac{\dot{p}_{\mathcal{A}_\mathbf{s}}(t)}{p_{\mathcal{A}_\mathbf{s}}(t)} - \mu$ | $-\frac{\ddot{f}_\mathbf{s}(t)}{\dot{f}_\mathbf{s}(t)} - \mu$ | $-\mathbf{s}^\top \mathbf{M}^2e^{\mathbf{M}t}\mathbf{1}_n \Big/ \mathbf{s}^\top \mathbf{M}e^{\mathbf{M}t}\mathbf{1}_n - \mu$ | 12, S82 |
| expected system dec. rate | time$^{-1}$ | $\bar{\kappa}$ | $p_{\mathcal{A}_\mathbf{s}}(0) - \mu$ | $-\dot{f}_\mathbf{s}(0) - \mu$ | $-\mathbf{s}^\top \mathbf{M}\mathbf{1}_n - \mu$ | S21, S82 |

Table S1: Summary table of main results

#### S1.7 Examples of CMs with analytical solutions

In this section, we chose a few common system topologies that exemplify the general dynamic properties of metabolic systems and cover many of the most common scenarios for applying CMs to experimental data. Each example starts by defining the network (external inputs, states, fluxes, etc.). Then, we show the corresponding **M**-matrix and show the analytical solution for the labeling function **f** (if it has a simple enough expression). To fully understand the construction of the CMs and the illustrated dynamic properties, the reader is encouraged to read the theoretical sections 1.1 - 1.7.

##### S1.7.1 Metabolic system composed of two independent pools

Even systems as simple as independent metabolic pools with different and constant decay rates can show the age dependence of the decay rates in the whole system (*systems decay rate*) and the non-trivial relationship between the mean residence time of a metabolite and its expected decay rate when the system is growing. In fact, these systems are entirely realistic and, for example, can represent coexisting pools of cellular metabolites that have different decay rates $\kappa_1$ and $\kappa_2$.

CM representation of such systems consists of independently fed pools and the contributed turnovers of the pools represent the individual decay rates. A two-pool system (Figure S7 A and B) can be represented as:

$$\begin{array}{c} E \xrightarrow{\kappa_{e1}} S_1 \\ \quad\;\;\downarrow \\ E \xrightarrow{\kappa_{e2}} S_2 \\ \quad\;\;\downarrow \end{array} \tag{S83}$$

$$\mathbf{M} = \begin{bmatrix} -\kappa_{e1} & 0 \\ 0 & -\kappa_{e2} \end{bmatrix} \tag{S84}$$

Equation 20 and the above **M**-matrix can be used to determine labeling dynamics of the system pools, $\mathbf{f}(t)$. In this case the dynamic equation can be solved analytically:

$$\mathbf{f} = \mathrm{e}^{\mathbf{M}t}\mathbf{1} = \exp\left(\begin{bmatrix} -\kappa_{e1} & 0 \\ 0 & -\kappa_{e2} \end{bmatrix} t\right)\begin{bmatrix} 1 \\ 1 \end{bmatrix} = \begin{bmatrix} \mathrm{e}^{-\kappa_{e1}t} & 0 \\ 0 & \mathrm{e}^{-\kappa_{e2}t} \end{bmatrix}\begin{bmatrix} 1 \\ 1 \end{bmatrix} = \begin{bmatrix} \mathrm{e}^{-\kappa_{e1}t} \\ \mathrm{e}^{-\kappa_{e2}t} \end{bmatrix} \tag{S85}$$

Using the range of experimentally observed decay rates of yeast proteins and typical growth rates of budding yeast as a realistic scenario (Figure S7 A and B), we can see that age-dependent decay and non-inverse relationship between mean residence times and expected decay rates can be realistically observed in live metabolic systems.

We can also use the labeling dynamics to quantify the dynamic parameters of this metabolic system (see Table S1). For example, the mean age of the system $\bar{\mathcal{A}}_\mathbf{s}$ can be quantified as:

$$\bar{\mathcal{A}}_\mathbf{s} = -\mathbf{s}^\top\mathbf{M}^{-1}\mathbf{1} = -\begin{bmatrix} s_1 \\ s_2 \end{bmatrix}^\top \begin{bmatrix} -1/\kappa_{e1} & 0 \\ 0 & -1/\kappa_{e2} \end{bmatrix}\begin{bmatrix} 1 \\ 1 \end{bmatrix} = s_1/\kappa_{e1} + s_2/\kappa_{e2} \tag{S86}$$

##### S1.7.2 Irreversible metabolic chain of length 2

It is not generally possible to determine the dynamic properties of an observed system when the input is delayed without knowing the labeling delay. To illustrate this *identifiability* problem, we will use a cascade of two metabolic pools (metabolic chain) as an example.

$$\begin{array}{c} E \xrightarrow{\kappa_{e1}} S_1 \xrightarrow{\kappa_{12}} S_2 \\ \qquad\;\;\downarrow \qquad\quad\downarrow \end{array} \tag{S87}$$

$$\mathbf{M} = \begin{bmatrix} -\kappa_{e1} & 0 \\ \kappa_{12} & -\kappa_{12}. \end{bmatrix} \tag{S88}$$

Here, $S_1$ and $S_2$ represent subsystems without and with delayed input, respectively. By plugging the system **M**-matrix into Equation 20 and using Sympy we can get the analytical expression for the labeling dynamics:

$$\mathbf{f} \equiv \begin{bmatrix} f_1(t) \\ f_2(t) \end{bmatrix} = \mathrm{e}^{\mathbf{M}t}\mathbf{1} = \exp\left(\begin{bmatrix} -\kappa_{e1} & 0 \\ \kappa_{12} & -\kappa_{12} \end{bmatrix} t\right)\begin{bmatrix} 1 \\ 1 \end{bmatrix} = \begin{bmatrix} \mathrm{e}^{-\kappa_{e1}t} & 0 \\ \frac{\mathrm{e}^{-\kappa_{e1}t}-\mathrm{e}^{-\kappa_{12}t}}{1-\kappa_{e1}/\kappa_{12}} & \mathrm{e}^{-\kappa_{12}t} \end{bmatrix}\begin{bmatrix} 1 \\ 1 \end{bmatrix}$$
$$= \begin{bmatrix} \mathrm{e}^{-\kappa_{e1}t} \\ \frac{\kappa_{12}\mathrm{e}^{-\kappa_{e1}t}}{\kappa_{12}-\kappa_{e1}} + \frac{\kappa_{e1}\mathrm{e}^{-\kappa_{12}t}}{\kappa_{e1}-\kappa_{12}} \end{bmatrix} \tag{S89}$$

Now imagine that we are analyzing the labeling of the second pool $f_2(t)$ only, and attempting to infer its intrinsic dynamic properties (defined by $\kappa_{12}$) based on these observations. We can easily notice that the labeling curve of the second pool can always be characterized by two identical solutions, as permuting $\kappa_{e1}$ and $\kappa_{12}$ gives identical formulas (see Figure S7C for the numeric simulation). Therefore, even the most precise measurements of the second pool are not sufficient to identify its intrinsic dynamic characteristics. However, this is entirely possible with the knowledge of the labeling dynamics of the first pool.

##### S1.7.3 Irreversible metabolic chain of arbitrary length

Imagine a cascade of reactions at steady-state:

$$\xrightarrow{\kappa_1} \underset{\downarrow}{S_1} \xrightarrow{\kappa_2} \underset{\downarrow}{S_2} \xrightarrow{\kappa_3} \dots \xrightarrow{\kappa_n} \underset{\downarrow}{S_n} \tag{S90}$$

Such models can be practical, for example, when describing irreversible processes such as protein turnover McShane et al. [2016]. The **M**-matrix that describes the system is therefore:

$$\mathbf{M} = \begin{bmatrix} -\kappa_1 & 0 & 0 & \dots & 0 & 0 \\ \kappa_2 & -\kappa_2 & 0 & \dots & 0 & 0 \\ 0 & \kappa_3 & -\kappa_3 & \dots & 0 & 0 \\ \vdots & & & & & \\ 0 & 0 & 0 & \dots & 0 & -\kappa_n \end{bmatrix}. \tag{S91}$$

It is possible to express the exponent of $\mathbf{M}t$ by directly deriving the Jordan canonical form. However, we chose to use Sympy to get the analytical expression for it in the case of $n = 3$:

$$\mathrm{e}^{\mathbf{M}t} = \begin{bmatrix} \mathrm{e}^{-\kappa_1 t} & 0 & 0 \\ \frac{\mathrm{e}^{-\kappa_1 t}}{1-\kappa_1/\kappa_2} - \frac{\mathrm{e}^{-\kappa_2 t}}{1-\kappa_1/\kappa_2} & \mathrm{e}^{-\kappa_2 t} & 0 \\ \frac{\mathrm{e}^{-\kappa_1 t}}{(1-\kappa_1/\kappa_2)(1-\kappa_1/\kappa_3)} - \frac{\mathrm{e}^{-\kappa_2 t}}{(1-\kappa_2/\kappa_1)(1-\kappa_2/\kappa_3)} + \frac{\mathrm{e}^{-\kappa_3 t}}{(1-\kappa_3/\kappa_1)(1-\kappa_3/\kappa_2)} & \frac{\mathrm{e}^{-\kappa_2 t}}{1-\kappa_2/\kappa_3} - \frac{\mathrm{e}^{-\kappa_3 t}}{1-\kappa_2/\kappa_3} & \mathrm{e}^{-\kappa_3 t} \end{bmatrix}. \tag{S92}$$

**Labeling dynamics** In order to find the solution for the labeling, we can now use the last result in Equation S67 to yield labeling dynamics in the case n = 3:

$$\mathbf{f} = \mathrm{e}^{\mathbf{M}t}\mathbf{1}_n = \begin{bmatrix} \mathrm{e}^{-\kappa_1 t} \\ \mathrm{e}^{-\kappa_1 t}\left(1-\frac{\kappa_1}{\kappa_2}\right)^{-1} + \mathrm{e}^{-\kappa_2 t}\left(1-\frac{\kappa_2}{\kappa_1}\right)^{-1} \\ \mathrm{e}^{-\kappa_1 t}\left(1-\frac{\kappa_1}{\kappa_2}\right)^{-1}\left(1-\frac{\kappa_1}{\kappa_3}\right)^{-1} + \mathrm{e}^{-\kappa_2 t}\left(1-\frac{\kappa_2}{\kappa_1}\right)^{-1}\left(1-\frac{\kappa_2}{\kappa_3}\right)^{-1} + \mathrm{e}^{-\kappa_3 t}\left(1-\frac{\kappa_3}{\kappa_1}\right)^{-1}\left(1-\frac{\kappa_3}{\kappa_2}\right)^{-1} \end{bmatrix}. \tag{S93}$$

This can be generalized to any arbitrary chain length:

$$f_i(t) = \sum_{j=1}^{i} \mathrm{e}^{-\kappa_j t}\left(\prod_{k=1,k\neq j}^{i} 1-\kappa_j/\kappa_k\right)^{-1} \tag{S94}$$

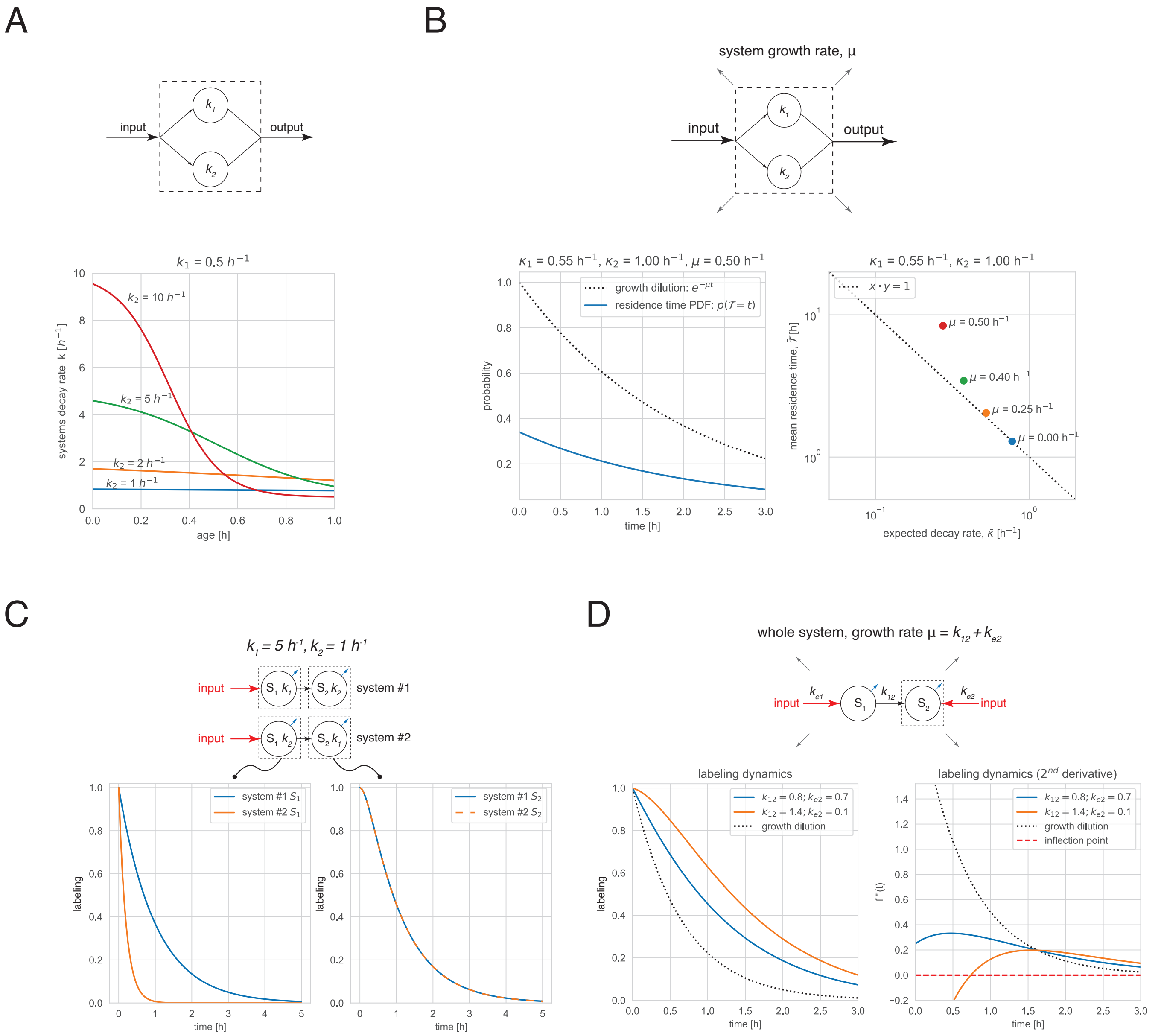


Figure S7: **Examples of compartmental models** (A) A system of two equally-sized unconnected pools with different constant decay rates $\kappa_1$ and $\kappa_2$ (dashed box). For a realistic scenario $\kappa_1$ is set to a characteristic growth rate of budding yeast, $\kappa_1 = 0.5h^{-1}$, and, for example, would correspond to a stable protein, whereas $\kappa_2$ ranges between $1h^{-1} \leq \kappa_2 \leq 10h^{-1}$ – where the upper bound represents estimated decay rate of a fast-degrading yeast protein Xie and Varshavsky [2001], Wang et al. [2010]. Note that when the two decay rates are similar (e.g. blue curve), the systems decay rate is nearly age-independent. However, as the difference increases, the dependence on age becomes pronounced (e.g. red curve).(B) A growing two-pool system as in (A). Left: an example of residence time distribution in a growing system. Right: a non-inverse relationship between the mean residence times and expected decay rates assuming that the system is growing at different rates $\mu$. Note that even relatively mild dissimilarity in the decay rates, $\kappa_1 = 0.55h^{-1}$ vs $\kappa_1 = 1h^{-1}$, leads to a visible deviation from the inverse relation (red) when the system is growing. (C) Observed labeling dynamics (dashed boxes) of the first $S_1$ (left) and second $S_2$ (right) pool in a two-pool metabolic chain. Note that changing the order of the pool's decay rates (system 1 $(\kappa_1, \kappa_2)$ versus system 2 $(\kappa_2, \kappa_1)$) results in identical labeling dynamics of the second pool $S_2$. (D) Labeling dynamics of the second pool (dashed box) in a metabolic chain when the observed pool is additionally fed by an external source. The orange curves represent a case where the the direct contribution of the external source $k_{e2}$ is small enough to observe an inflection point around 0.75 $h$, which disappears with large enough $k_{e2}$ (blue curves). However, since the labeling dynamics exceed the growth dilution (dotted line, left panel), we can conclude that the system input is delayed.

Interestingly, if we focus only on the labeling of the last state ($f_n(t)$), the function can be written as:

$$f_n(t) = \sum_{j \in \mathcal{J}} e^{-\kappa_j t} \left( \prod_{k \in \mathcal{J} \setminus \{j\}} 1 - \kappa_j / \kappa_k \right)^{-1} \tag{S95}$$

where $\mathcal{J} \equiv \{1, 2, \ldots, n\}$. Therefore, one can see that the function is symmetric to the ordering of the $\kappa_i$ values. This result recapitulates the one from section S1.7.2 showing that if we only measure the labeling of the last state, the order of decay rates is unidentifiable.

**Mean ages** To compute the mean ages of each pool, we first compute the inverse of $\mathbf{M}$ (again, using Sympy) and use it in Equation S78 which gives us:

$$\bar{\mathcal{A}} = -\mathbf{M}^{-1}\mathbf{1}_n = -\begin{bmatrix} -\kappa_1^{-1} & 0 & 0 & \ldots & 0 \\ -\kappa_1^{-1} & -\kappa_2^{-1} & 0 & \ldots & 0 \\ -\kappa_1^{-1} & -\kappa_2^{-1} & -\kappa_3^{-1} & \ldots & 0 \\ \vdots & & & & \\ -\kappa_1^{-1} & -\kappa_2^{-1} & -\kappa_3^{-1} & \ldots & -\kappa_n^{-1} \end{bmatrix} \mathbf{1}_n = \begin{bmatrix} \kappa_1^{-1} \\ \kappa_1^{-1} + \kappa_2^{-1} \\ \kappa_1^{-1} + \kappa_2^{-1} + \kappa_3^{-1} \\ \vdots \\ \sum_{j=1}^{n} \kappa_j^{-1} \end{bmatrix}, \tag{S96}$$

this leads us to the following general solution:

$$\bar{\mathcal{A}}_i = \sum_{j=1}^{i} \kappa_j^{-1}. \tag{S97}$$

The mean system age, mean residence times, and expected decay rates are:

$$\begin{aligned} \bar{\mathcal{A}} &= -\frac{\mathbf{s}^\top \mathbf{M}^{-1}\mathbf{1}_n}{\mathbf{s}^\top \mathbf{1}_n} = \frac{\sum_i s_i \sum_{j=1}^{i} \kappa_j^{-1}}{\sum_{i=1}^{n} s_i} \\ \bar{\mathcal{T}} &= -\frac{s^\top \mathbf{1}_n}{s^\top \mathbf{M}\mathbf{1}_n} = \frac{\sum_{i=1}^{n} s_i}{\kappa_1^{-1} s_1} \\ \bar{\kappa} &= -\frac{\mathbf{s}^\top \mathbf{M}\mathbf{1}_n}{\mathbf{s}^\top \mathbf{1}_n} = \frac{\kappa_1^{-1} s_1}{\sum_{i=1}^{n} s_i}. \end{aligned} \tag{S98}$$

**Non-distinct eigenvalues** For our general solution in Equation S94, we made an important implicit assumption, that none of the $\kappa_i$ are equal. If there exist $i \neq j$ where $\kappa_i = \kappa_j$, then the Jordan canonical form would not be a diagonal matrix (and also we would get that $g_{ij} = \infty$). Solving all such cases analytically for an arbitrarily large system is not possible. We can, however, solve the most simple case where and $\kappa_1 = \kappa_2 = \ldots = \kappa_n = \kappa$. I similar derivation can also be found in Sokol and Portais [2015].

$$\mathbf{M} = \begin{bmatrix} -\kappa & 0 & \ldots & 0 & 0 \\ \kappa & -\kappa & \ldots & 0 & 0 \\ \vdots & \ddots & \ddots & \vdots & \vdots \\ 0 & 0 & \ldots & -\kappa & 0 \\ 0 & 0 & \ldots & \kappa & -\kappa \end{bmatrix} \tag{S99}$$

Also in this case, Sympy can directly solve the equation, which in general turns out to be:

$$f_i(t) = \mathrm{e}^{-\kappa t} \sum_{j=0}^{i-1} \frac{(\kappa t)^j}{j!}. \tag{S100}$$

Interestingly, the sum is exactly equal to the first $i$ terms in the Taylor expansion of $\mathrm{e}^{\kappa t}$. We can expect that for an infinitely long pathway at any given finite time point $t$, there will be a place down the line where the states are still fully labeled (since the "wave" of wash-out hasn't reached them yet). The labeled fraction at the limit will be:

$$f_\infty(t) = \mathrm{e}^{-\kappa t} \sum_{j=0}^{\infty} \frac{(\kappa t)^j}{j!} = \mathrm{e}^{-\kappa t}\mathrm{e}^{\kappa t} = 1 \tag{S101}$$

##### S1.7.4 Reversible metabolic chain of arbitrary length

A general case of a metabolic chain can be represented with CM as a set of consequent pools of intermediates – pools $S_1 \to S_n$ – fed by internalized nutrient from the external source $E$. The consequent pools exchange though metabolic reactions with arbitrary rate, but that satisfy the mass-balance constraint. The pool can also have arbitrary outgoing fluxes that e.g. support cell growth or other metabolic pathways.

$$\begin{array}{ccccccccc} E \xrightarrow{\kappa_{e1}} & S_1 & \underset{\kappa_{21}}{\overset{\kappa_{12}}{\rightleftharpoons}} & S_2 & \underset{\kappa_{32}}{\overset{\kappa_{23}}{\rightleftharpoons}} & S_3 \cdots & S_{n-1} & \underset{\kappa_{n,n-1}}{\overset{\kappa_{n-1,n}}{\rightleftharpoons}} & S_n \\ & \downarrow & & \downarrow & & \downarrow & \downarrow & & \downarrow \end{array} \tag{S102}$$

Although there is no analytical expression describing the dynamics for general metabolic chains (for $n > 2$) we can still prove some general claims about monotonicity. First, we note that the $\mathbf{M}$ matrix is tridiagonal with the following pattern:

$$\mathbf{M} = \begin{bmatrix} -\kappa_{e1} - \kappa_{21} & \kappa_{21} & \dots & 0 & 0 \\ \kappa_{12} & -\kappa_{12} - \kappa_{32} & \dots & 0 & 0 \\ 0 & \kappa_{23} & \dots & 0 & 0 \\ 0 & 0 & \dots & 0 & 0 \\ \vdots & \ddots & \ddots & \ddots & \vdots \\ 0 & 0 & \dots & \kappa_{n-1,n-2} & 0 \\ 0 & 0 & \dots & -\kappa_{n-2,n-1} - \kappa_{n,n-1} & \kappa_{n,n-1} \\ 0 & 0 & \dots & \kappa_{n-1,n} & -\kappa_{n-1,n} \end{bmatrix}. \tag{S103}$$

Now, we define $\kappa_1 \equiv \kappa_{e1}$ and $\theta_1 \equiv \kappa_{21}/\kappa_{e1}$ and for any $i > 1$:

$$\kappa_i \equiv \kappa_{i-1,i} \qquad \theta_i \equiv \frac{\kappa_{i+1,i}}{\kappa_{i-1,i}}. \tag{S104}$$

One can think of $\kappa_i$ as the turnover rate for $S_i$ (if the reverse flux were 0) and $\theta_i$ as the ratio of inputs into $S_i$ from the two opposite directions. Using these definitions, we can rewrite $\mathbf{M}$ as:

$$\mathbf{M} = \begin{bmatrix} -(1+\theta_1)\kappa_1 & \theta_1\kappa_1 & \dots & 0 & 0 \\ \kappa_2 & -(1+\theta_2)\kappa_2 & \dots & 0 & 0 \\ 0 & \kappa_3 & \dots & 0 & 0 \\ 0 & 0 & \dots & 0 & 0 \\ \vdots & \ddots & \ddots & \ddots & \vdots \\ 0 & 0 & \dots & \theta_{n-2}\kappa_{n-2} & 0 \\ 0 & 0 & \dots & -(1+\theta_{n-1})\kappa_{n-1} & \theta_{n-1}\kappa_{n-1} \\ 0 & 0 & \dots & \kappa_n & -\kappa_n \end{bmatrix}. \tag{S105}$$

**Lemma S1.3.** *The mean ages for state i is given by:*

$$\bar{\mathcal{A}}_i = \sum_{j=1}^{n} \kappa_j^{-1} \left( \sum_{l=1}^{\min(i,j)} \prod_{k=l}^{j-1} \theta_k \right) \tag{S106}$$

*where the empty product is defined to be 1, i.e.* $\prod_{k=l}^{l-1} \theta_k = 1$.

*Proof.* Using Sympy, we can find the inverse of $\mathbf{M}$:

$$\mathbf{M}^{-1} = \begin{bmatrix} -\kappa_1^{-1} & -\kappa_2^{-1}\theta_1 & -\kappa_3^{-1}\theta_2\theta_1 & -\kappa_4^{-1}\theta_3\theta_2\theta_1 & \dots & -\kappa_n^{-1}\sum_{l=1}^{1}\prod_{k=l}^{n-1}\theta_k \\ -\kappa_1^{-1} & -\kappa_2^{-1}(\theta_1+1) & -\kappa_3^{-1}(\theta_2\theta_1+\theta_2) & -\kappa_4^{-1}(\theta_3\theta_2\theta_1+\theta_3\theta_2) & \dots & -\kappa_n^{-1}\sum_{l=1}^{2}\prod_{k=l}^{n-1}\theta_k \\ -\kappa_1^{-1} & -\kappa_2^{-1}(\theta_1+1) & -\kappa_3^{-1}(\theta_2\theta_1+\theta_2+1) & -\kappa_4^{-1}(\theta_3\theta_2\theta_1+\theta_3\theta_2+\theta_3) & \dots & -\kappa_n^{-1}\sum_{l=1}^{3}\prod_{k=l}^{n-1}\theta_k \\ -\kappa_1^{-1} & -\kappa_2^{-1}(\theta_1+1) & -\kappa_3^{-1}(\theta_2\theta_1+\theta_2+1) & -\kappa_4^{-1}(\theta_3\theta_2\theta_1+\theta_3\theta_2+\theta_3+1) & \dots & -\kappa_n^{-1}\sum_{l=1}^{4}\prod_{k=l}^{n-1}\theta_k \\ \vdots & \vdots & \vdots & \vdots & & \\ -\kappa_1^{-1} & -\kappa_2^{-1}(\theta_1+1) & -\kappa_3^{-1}(\theta_2\theta_1+\theta_2+1) & -\kappa_4^{-1}(\theta_3\theta_2\theta_1+\theta_3\theta_2+\theta_3+1) & \dots & -\kappa_n^{-1}\sum_{l=1}^{n}\prod_{k=l}^{n-1}\theta_k \end{bmatrix} . \tag{S107}$$

i.e. for any row $i$ and column $j$ where $i \leq j$, the element of the inverse matrix will be $-\kappa_j^{-1}\sum_{l=1}^{i}\prod_{k=l}^{j-1}\theta_k$. For cases where $i > j$, the value is equal to the diagonal value in the same column, i.e. $-\kappa_j^{-1}\sum_{l=1}^{j}\prod_{k=l}^{j-1}\theta_k$. So, in general, the sum will be over $l \in \{1, \dots, \min(i,j)\}$.

$$[\mathbf{M}^{-1}]_{i,j} = -\kappa_j^{-1}\left(\sum_{l=1}^{\min(i,j)}\prod_{k=l}^{j-1}\theta_k\right) . \tag{S108}$$

Using Equation S78 we can see that $\bar{\mathcal{A}}_i$ is minus the sum of elements in row $i$ of $\mathbf{M}^{-1}$, which proves the lemma. ∎

**Corollary S1.3.1.** The mean ages increase along the pathway, i.e. $\bar{\mathcal{A}}_{i-1} \leq \bar{\mathcal{A}}_i$.

*Proof.* Using Equation S106, we can see that the difference between the mean ages of two consecutive states is:

$$\bar{\mathcal{A}}_i - \bar{\mathcal{A}}_{i-1} = \sum_{j=i}^{n}\kappa_j^{-1}\prod_{k=i}^{j-1}\theta_k \tag{S109}$$

which is a positive number. ∎

**Corollary S1.3.2.** The labeled fractions at any given time $t$ increase along the pathway, i.e. $f_{i-1} \leq f_i$.

*Proof.* We can rewrite the ODE in Equation S63 (where we use the fact that in this case $\mathbf{g} = \mathbf{0}_n$) as:

$$\mathbf{f} = \mathbf{M}^{-1}\dot{\mathbf{f}} . \tag{S110}$$

Therefore, we get the following formula for the labeled fractions:

$$f_i = -\sum_{j=1}^{n}\kappa_j^{-1}\dot{f}_j\left(\sum_{l=1}^{\min(i,j)}\prod_{k=l}^{j-1}\theta_k\right) . \tag{S111}$$

and the difference between two consecutive states will be:

$$f_i - f_{i-1} = -\sum_{j=i}^{n}\kappa_j^{-1}\dot{f}_j\prod_{k=i}^{j-1}\theta_k \tag{S112}$$

From Equation S70 we know that $\frac{df_i}{dt} \leq 0$, and all other variables ($\kappa_j^{-1}$ and $\theta_k$) are positive. Therefore, we can see that $f_i - f_{i-1} \geq 0$. ∎

There is one more interesting observation we can make based on Equation S112. Given $f_{i-1}$, the value of $f_i$ depends only on parameters $\kappa_j$, $\dot{f}_j$, and $\theta_k$ where $j, k \geq i$. This means that if we measure $f_{i-1}$ directly, we can simply ignore the details of anything upstream of $S_i$ and directly fit the values of the model based on the parameters surrounding $S_i$ and its downstream states. Reflecting on this result, it might not be surprising since $S_{i-1}$ is the only source of label coming into $S_i$ and by measuring it we can know everything we need to know about it. In other words, information can only flow into $S_i$ via $S_{i-1}$ and its labeling pattern. In section S1.9, we address this idea in more detail.

**Maturation time in a linear reversible pathway** We define maturation time as the median age ($t_{1/2}$), i.e. the time after which a protein that starts in $S_i$ to have a 50% chance of reaching $S_{i+1}$. Since we know that even for reversible chains (Corollary S1.3.2) the labeled fractions increase along the pathway, so will the maturation times be monotonically increasing.

##### S1.7.5 Irreversible metabolic chain of length 2 with two external sources

If not all input into the observed system is delayed, the inflection of the labeling curve may not always be observed. This may happen, for example, if the direct input from the external source overwhelmingly exceeds contributions from the internal sources. We will exemplify this situation using an irreversible metabolic chain whose second (observed) pool is fed by an external source.

$$E \xrightarrow{\kappa_{e1}} \underset{\downarrow}{S_1} \xrightarrow{\kappa_{12}} \underset{\downarrow}{S_2} \xleftarrow{\kappa_{e2}} E \tag{S113}$$

The **M**-matrix corresponding to this system is:

$$\mathbf{M} = \begin{bmatrix} -\kappa_{e1} & 0 \\ \kappa_{12} & -\kappa_{12} - \kappa_{e2}. \end{bmatrix} \tag{S114}$$

We can see (Figure S7D, blue curve) that the labeling dynamics of the second pool, $f_2(t)$, does not have an inflection point ($\ddot{f}_2(t)$ does not change sign) when the external contribution $k_{e2}$ to the second pool is large enough. Interestingly, if the system growth rate, $\mu$, is sufficiently high, the growth dilution would exceed the labeling dynamics of the second pool, thus pointing to the delayed input even in the absence of the curve inflection (see Appendix S1.4.4).

##### S1.7.6 Uniformly-decaying systems

Aside from the fact that bulk metabolite labeling is a monotonously decreasing and convex function, deducing the internal metabolic routes from dynamic labeling is difficult, since the former is the result of a complex interplay between internal and external fluxes and metabolite pool sizes (see Equation S72). A special case where this is possible is when the efflux of material occurs by dilution and/or degradation at the same kinetic rate in all the internal pools. In this case, the efflux in each pool is proportional to the size of the pool, a proportionality factor denoted the *system turnover rate* $\kappa_u$, that is, $v_{i\to\emptyset} = \kappa_u s_i$. Note that if system turnover is dominated by growth (e.g., as in the case of rapidly growing cells where degradation is negligible), then system turnover will be equal to the growth rate: $\kappa_u = \mu$.

An important property of **M**, is that $-\mathbf{s}^\top\mathbf{M}$ is a vector containing the output exchange fluxes (i.e. $v_{i\to\emptyset}$, see Appendix S1.11.2). We therefore can see that:

$$-\mathbf{s}^\top\mathbf{M} = \kappa_u\, \mathbf{s}^\top\,. \tag{S115}$$

In other words, **s** is an eigenvector of **M**.

Now, let us consider the system labeled fraction (Equation S71):

$$f_\mathbf{s} = \mathbf{s}^\top\mathbf{f} \tag{S116}$$

where we dropped the denominator $\mathbf{s}^\top\mathbf{1}_n$ since without loss of generality, we can assume that it is equal to 1. Now, we can revisit Equation S63 which is an ODE describing the labeling dynamics:

$$\dot{\mathbf{f}} = \mathbf{M}\mathbf{f} + \mathbf{g}\,. \tag{S117}$$

where we can assume, as before, that **g** is zero for pulse-labeling experiments (in systems with direct input). Now, multiplying both sides by $\mathbf{s}^\top$ we get:

$$\begin{aligned} \mathbf{s}^\top\dot{\mathbf{f}} &= \mathbf{s}^\top(\mathbf{M}\mathbf{f}) = -\kappa_u\left(\mathbf{s}^\top\,\mathbf{f}\right) \\ \dot{f}_\mathbf{s} &= -\kappa_u\, f_\mathbf{s} \\ f_\mathbf{s} &= \mathrm{e}^{-\kappa_u\, t}\,. \end{aligned} \tag{S118}$$

In conclusion, for any system with direct input and uniform decay rates, the system labeling function ($f_\mathbf{s}$) behaves as if it were a single well-mixed pool. Internally, of course, the different states can diverge in their labeling dynamics, as we will see in the examples below.

##### S1.7.7 Uniformly-decaying irreversible chain of arbitrary length

Consider a system with a single system turnover (as in S1.7.6) which is an irreversible chain. In addition, the final product $S_n$ is a dead-end (e.g. it could be a direct constituent of biomass like a membrane lipid). Based on the mass-balance constraint, and the fact that the efflux from state $i$ is $\kappa_u s_i$, we can see that:

$$v_{i-1\to i} = v_{i\to i+1} + v_{i\to\emptyset} = v_{i\to i+1} + \kappa_u s_i = \ldots = \kappa_u \sum_{j=i}^{n} s_j$$
$$\kappa_i = v_{i-1\to i}/s_i = \kappa_u \underbrace{\sum_{j=i}^{n} s_j/s_i}_{\phi_i^{-1}} = \kappa_u/\phi_i\,, \tag{S119}$$

where we define $\phi_i$ as the relative pool size of $s_i$ compared to all the downstream pool (including $s_i$ itself). Now we can use the solution we derived for general irreversible chains (Equation S94):

$$f_i(t) = \sum_{j=1}^{i} \mathrm{e}^{-\kappa_u t/\phi_j} \left( \prod_{k=1, i\neq j}^{i} 1 - \phi_k/\phi_j \right)^{-1} \tag{S120}$$

and specifically for chains of length $n = 2$:

$$f_2(t) = \frac{\phi_1}{\phi_1 - 1}\mathrm{e}^{\kappa_u t/\phi_1} + \frac{1}{1-\phi_1}\mathrm{e}^{\kappa_u t} \tag{S121}$$

#### S1.8 Examples of CM applications in analyzing experimental data

##### S1.8.1 Determining the order and timescales of sequential cellular processes

Dynamic labeling and the metabolic age metric can be especially advantageous to analyze specific types of cellular processes. One such example is sequential processes, known as metabolic chains. Using the CM representation, one can show that in a metabolic chain (a set of consecutive interconversions of metabolites), the upstream metabolic pools are always younger and their labeling is washed-out earlier than the downstream ones, that is, at any time $t \geq 0$:

$$f_{i-1}(t) \leq f_i(t)\,, \tag{S122}$$

where $i$ is the sequential pool index (see Appendix S1.7.4). This property enables the use of dynamic labeling as a simple way to determine the order and time-scale of events in such processes.

To illustrate this in practice, we reanalyze the data from Onischenko et al. [2020] using our age-based framework to infer the assembly order of the yeast nuclear pore complex (NPC) based on the labeling dynamics of affinity-isolated protein complexes between the bait and prey NPC subunits (Figure S8A). In this study, the assembly sequence was inferred based on the analysis of the size of precursor pools of prey subunits before they are co-isolated with baits, which relied on fitting a prey labeling dynamics with specific CM (called KSM) and required a number of specific assumptions such as that there is a single precursor pool of the prey, that assembly is irreversible, or the preys are indestructible. In addition, to avoid biases due to errors in individual pool estimates, this framework required determining plausible assembly clusters between proteins in advance based on the correlation clustering of the labeling data. We can now illustrate how using CM and age-based interpretation of labeling dynamics can circumvent requirement of a number of these assumptions and also allow a more simple and unbiased way for estimating the NPC assembly order and time-scale.

To envision how bait-prey labeling dynamics is connected to the assembly order, we consider that from the perspective of the traced metabolite (lysine in this case) incorporated into a prey, its fate after entering the cell represents a metabolic chain in which, according to Equation S122, the later the NPC bait joins the NPC prey, the slower the labeling of lysine in the co-isolated prey pool would appear. In other words, the order in which the baits co-assemble with a given prey can be inferred by comparing the prey labeling (i.e., lysine labeling) in affinity-isolated complexes with these baits(Figure S8B). If we would focus just on the set of bait subunits, each of them co-isolates all the other baits as prey, and the order in which this set assembled with each other can be recapitulated by knowing all such partial assembly orders based on the labeling values.

To implement this, we first calculate an *age score* (see Figure S14A and Materials and Methods S2.10) as the area under the labeling curve (AUC) between the first (t = 0) and last (t = 90 min) measurement, using a 3-pool CM to interpolate the observed experimental labeling dynamics and obtain partial assembly orders of baits with each prey based on the principles described above. Indeed, although a CM is used to fit the labeling dynamics, this is merely a way to realistically interpolate the experimental labeling values without making specific assumptions about the assembly mechanism and allowing (by calculating the AUC) to compare the heights of the labeling curves between baits. To ensure model-independence, we have alternatively determined age-scores using a non-parametric method (see Appendix S1.10.3) and arrived at a nearly-identical assembly model (Figures S8C and S14E).

To determine the assembly sequence based on the partial assembly orders, we probed all possible hierarchical assembly models for consistency with those, using Spearman rank correlation for grading them (see Figure S14B and C and

Materials and Methods Section S2.11). This showed that high-scoring assembly hierarchies could not be produced randomly while the highest-scoring assembly models were in good agreement with the previously reported three-tier model of NPC assembly Onischenko et al. [2020] and correctly predicting the immediate assembly of baits interacting directly (Figures S8C and S14F). Furthermore, this framework appeared resistant to the selection of baits used for the analysis. For example, the addition of another Nsp1 bait that was not previously used in our analyzes did not change the overall assembly hierarchy and correctly predicted its initial co-assembly of this protein with other members of its biochemical sub-complex (Figure S14E). Together, this exemplifies how the age-based interpretation of metabolic labeling and CMs can be used to dissect the order of metabolic events.

The age interpretation labeling dynamics allows for gaining insight into the time scale of cellular processes. We can illustrate this in the same example by evaluating with minimal assumptions the *minimal maturation time* $\bar{T}_{\min}$ required for the NPC subunits to assemble in the mature NPC, marked by latest joining protein, Mlp1, in our assembly model (Figure S8C). To this end, we consider that the metabolic age of lysine $\mathcal{A}$ in preys co-isolated with Mlp1 is the sum of three components:

$$\mathcal{A} = \mathcal{A}^{\circ} + T + \mathcal{A}_{\mathrm{NPC}} \tag{S123}$$

where $\mathcal{A}^{\circ}$ is the age of lysine in the input, $T$ is the maturation time of the subunit (i.e., the time required for the newly produced subunit to become part of the mature NPC), and $\mathcal{A}_{\mathrm{NPC}}$ is the reduced age of the subunit in the mature NPC (see Figure S8D). Furthermore, from linearity of expectation, the additive relationship in Equation S123 applies also to the mean values, which can be used to compute the mean maturation time of the subunits as:

$$\bar{T} = \bar{\mathcal{A}} - \bar{\mathcal{A}}^{\circ} - \bar{\mathcal{A}}_{\mathrm{NPC}}\,. \tag{S124}$$

Since the reduced mean ages are bound by growth dilution ($\bar{\mathcal{A}}_{\mathrm{NPC}} \leq \mu^{-1}$, see Equation 11), we can derive the constraint on the maturation time:

$$\bar{T} \geq \bar{T}_{\min} \equiv \bar{\mathcal{A}} - \bar{\mathcal{A}}^{\circ} - \mu^{-1}\,. \tag{S125}$$

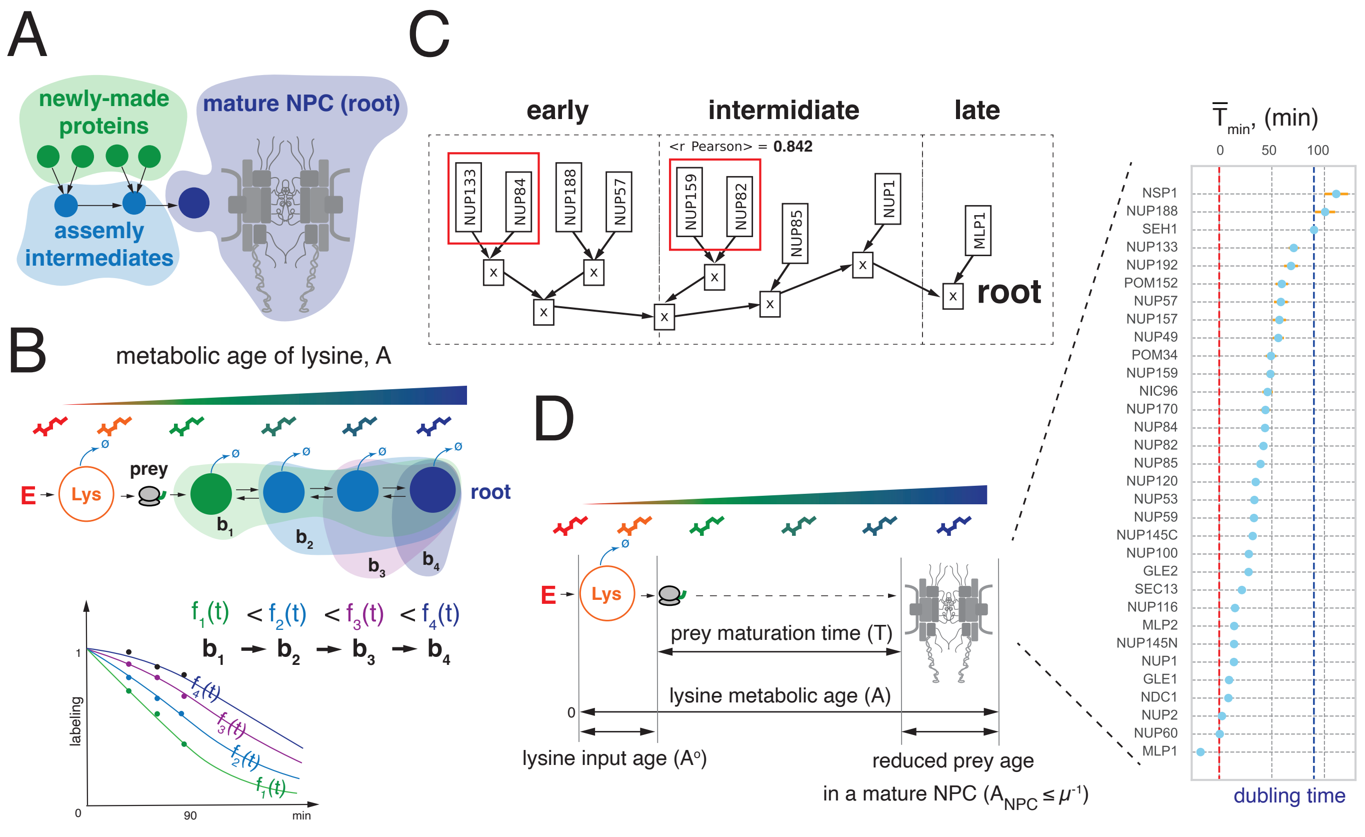


Figure S8: **Determining the order and time scale of NPC assembly using dynamic labeling.** (A) Hierarchical model of NPC assembly (assembly tree): newly-made NPC complements (leafs) first co-assemble into intermediates (branches) which in turn assemble into the mature structure (root). (B) Metabolic path of externally added lysine contained in a prey. The path is a metabolic chain that begins in the external pool (E) and ends in the pool of mature NPC (root). The age of lysine in a prey (colored areas) grows according to the order in which it co-assembles with baits which is reflected by progressively slower labeling dynamics of the prey. (C) Best-scoring assembly model of baits with each other. Red boxes depict known directly interacting proteins. (D) Minimal maturation times of NPC subunits required to assemble into Mlp1-positive NPCs.

To compute this lower bound (right side), we determined the mean metabolic age of lysine $\bar{\mathcal{A}}$ using experimental labeling values of the Mlp1 prey with Equation 4 and quantified $\bar{\mathcal{A}}^{\circ}$ based on our previous analysis of lysine labeling delay in these experiments Onischenko et al. [2020] (see Appendix S2.10). This showed that $\bar{T}_{\min}$ for most NPC components was on the order of one hour, in agreement with the slow incorporation of the Mlp1 subunit into NPCs Onischenko et al. [2020], Zsok et al. [2024] (Figure S8 D). In further support, Nsp1 – a subunit that shows an exceptionally large $\bar{T}_{\min}$ – has previously been reported to have large cytosolic precursor pools prior to incorporation into NPCs Otto et al. [2024].

Overall, these results demonstrate the utility of dynamic labeling and the metabolic age metric to determine the order and time scale of sequential cellular processes.

#### S1.9 Systems with delayed input and a single feeding source

Consider a subsystem with a delayed input (target), where the input comes only from another internal subsystem (source) as depicted in Figure 3C. In order to "compensate" for the delayed input into the target system, we need have information about the input, specifically the distribution of ages. To understand how this can be done, we first define 3 age variables: $\mathcal{A}$ – the age of a particle in the target subsystem, $\mathcal{A}^\circ$ – the age of a particle in the input stream, and $\mathcal{A}_r$ – the "reduced" age or the age in the target system if we were to reset the clock as soon as the particle entered it (i.e., equivalent to converting it a system with direct input). Therefore, by definition, $\mathcal{A} = \mathcal{A}^\circ + \mathcal{A}_r$. In order to solve the relationship between these ages, we must assume that $\mathcal{A}^\circ$ and $\mathcal{A}_r$ are independent random variables, i.e. the history of each particle in the target subsystem is erased as soon as it enters. In other words, everything that happens to the particle after entering the target subsystem is only a function of the time it spent there, and not on what happened in the source subsystem. Therefore, the age can be separated to two parts: the time that passed until it entered the target system, and the time it spent inside the target system. In general, the PDF of a sum of two independent variables is given by a convolution:

$$p_{\mathcal{A}}(t) = p_{\mathcal{A}^\circ + \mathcal{A}_r}(t) = \int_0^t p_{\mathcal{A}^\circ}(x)\ p_{\mathcal{A}_r}(t-x)\ dx\,. \tag{S126}$$

Note that if the two variables are not independent, the solution would depend on their joint distribution. We do not deal with this general case in this work.

Now, in order to equate it to labeling curves, we first need to get the expression for the cumulative distribution:

$$\begin{aligned}
P_{\mathcal{A}}(t) &= \int_0^t p_{\mathcal{A}}(\tau) d\tau \\
&= \int_0^t \left[\int_0^\tau p_{\mathcal{A}^\circ}(x)\ p_{\mathcal{A}_r}(\tau - x)\ dx\right] d\tau \\
&= \int_0^t \left[\int_0^t p_{\mathcal{A}^\circ}(x)\ p_{\mathcal{A}_r}(\tau - x)\ dx\right] d\tau \\
&= \int_0^t \int_0^t p_{\mathcal{A}^\circ}(x)\ p_{\mathcal{A}_r}(\tau - x)\ dx\ d\tau \\
&= \int_0^t \int_0^t p_{\mathcal{A}^\circ}(x)\ p_{\mathcal{A}_r}(\tau - x)\ d\tau\ dx \\
&= \int_0^t p_{\mathcal{A}^\circ}(x) \int_0^t p_{\mathcal{A}_r}(\tau - x)\ d\tau\ dx \\
&= \int_0^t p_{\mathcal{A}^\circ}(x) \left[\int_0^t p_{\mathcal{A}_r}(\tau - x)\ d\tau\ \right] dx \\
&= \int_0^t p_{\mathcal{A}^\circ}(x)\ P_{\mathcal{A}_r}(t - x)\ dx
\end{aligned} \tag{S127}$$

where we could set the limit of the inner integral to $[0, t]$ instead of $[0, \tau]$ because $\forall x > \tau\ :\ p_{\mathcal{A}_r}(\tau - x) = 0$ and therefore $\int_\tau^t p_{\mathcal{A}^\circ}(x) p_{\mathcal{A}_r}(\tau - x) dx = 0$

Now, if we define $f^\circ$, $f(t)$, and $f_r$, as the labeling function of the input stream, target subsystem, and reduced system (respectively) we can use the known relationship between labeling and age:

$$\begin{aligned} f(t) &= 1 - P_{\mathcal{A}}(t) \\ &= 1 - \int_0^t p_{\mathcal{A}^\circ}(x)\ P_{\mathcal{A}_r}(t-x)\ dx \\ &= 1 - \int_0^t p_{\mathcal{A}^\circ}(x)\ \left(1 - f_r(t-x)\right)\ dx \\ &= 1 - \int_0^t p_{\mathcal{A}^\circ}(x) + \int_0^t p_{\mathcal{A}^\circ}(x)\ f_r(t-x)\ dx \\ &= 1 - P_{\mathcal{A}^\circ}(t) - \int_0^t \dot{f}^\circ(x)\ f_r(t-x)\ dx \\ &= f^\circ(t) - (\dot{f}^\circ * f_r)(t)\,, \end{aligned} \tag{S128}$$

where $*$ indicates a convolution, i.e. $(g*h)(t) = \int_0^t g(\tau)h(t-\tau)d\tau$.

Going back to Equation S126, one can see that the terms $\mathcal{A}^\circ$ and $\mathcal{A}_r$ are commutable and therefore we can symmetrically derive:

$$\begin{aligned} f(t) &= 1 - P_{\mathcal{A}}(t) \\ &= 1 - \int_0^t p_{\mathcal{A}_r}(x)\ P_{\mathcal{A}^\circ}(t-x)\ dx \\ &= 1 - \underbrace{\int_0^t \underbrace{p_{\mathcal{A}_r}(x)}_{\text{fraction of age x}} \cdot \underbrace{\left(1 - f^\circ(t-x)\right)}_{\text{unlabeled input at entry}} \cdot dx}_{\text{fraction of unlabeled metabolite}}\,. \end{aligned} \tag{S129}$$

The last expression provides another interpretation of delayed labeling in which the observed labeled fraction can be seen as a complement to the content of the unlabeled species. This in turn is composed of the weighted sum among particles at each possible age (from 0 to $t$) in the reduced system that reserved their partial labeling (guided by the delay in labeling input) at the time of entry. This interpretation enables us to determine more complex labeling when, for example, target molecules (reduced system) receive more than one instance of the unlabeled input.

Furthermore, we can also develop the mean age formula directly using Equation S128:

$$\begin{aligned} \bar{\mathcal{A}} = \int_0^\infty f(t) = \int_0^\infty f^\circ(t)dt &- \int_0^\infty (\dot{f}^\circ * f_r)(t)dt \\ = \bar{\mathcal{A}}^\circ \quad &+ \left(\int_0^\infty -\dot{f}^\circ dt\right) \cdot \left(\int_0^\infty f_r dt\right) \\ = \bar{\mathcal{A}}^\circ \quad &+ \underbrace{\left(\int_0^\infty p_{\mathcal{A}^\circ}(t)dt\right)}_{=1} \cdot \bar{\mathcal{A}}_r \\ = \bar{\mathcal{A}}^\circ \quad &+ \bar{\mathcal{A}}_r \end{aligned}$$

where we use the well-known fact that the integral of a convolution is the product of the two integrals $\int_0^\infty (g*h)(t)dt = \left(\int_0^\infty g(t)dt\right) \cdot \left(\int_0^\infty h(t)dt\right)$.

Another way to derive Equation S128 is by considering that the labeling pattern that comes out of the source system is a time-variable input labeling function which can be used directly when solving the ODE that describes the CM (i.e. in Equation S63):

$$\dot{\mathbf{f}} = \mathbf{M}\mathbf{f} + \mathbf{g}\,. \tag{S130}$$

So far, we assumed that $\mathbf{g}$ is zero any $t > 0$, i.e. that we are performing a pulse labeling experiment where the media is switched to fully labeled at $t = 0$. However, if we instead use one that changes with time, i.e. a well-mixed source system described by $f^\circ(t)$, then we cannot drop $\mathbf{g}$ anymore. According to Equation S63, $\mathbf{g}$ represents the external fluxes (given by $\mathbf{M}\mathbf{1}_n$) multiplied by the labeling of the input ($f^\circ(t)$), i.e. $\mathbf{g}(t) = \mathbf{M}\mathbf{1}_n \cdot f^\circ(t)$. Therefore, the ODE becomes:

$$\dot{\mathbf{f}} = \mathbf{M}\mathbf{f} + \mathbf{M}\mathbf{1}_n \cdot f^\circ = \mathbf{M}\left(\mathbf{f} + \mathbf{1}_n f^\circ\right)\,. \tag{S131}$$

and its solution is based on a convolution:

$$\mathbf{f}(t) = \mathrm{e}^{\mathbf{M}t}\mathbf{1}_n + \int_0^t \mathrm{e}^{\mathbf{M}\tau}\mathbf{M}\mathbf{1}_n f^\circ(t-\tau)d\tau = \mathrm{e}^{\mathbf{M}t}\mathbf{1}_n + \int_0^t \mathbf{M}\mathrm{e}^{\mathbf{M}\tau}\mathbf{1}_n f^\circ(t-\tau)d\tau\,, \tag{S132}$$

where we assumed as always that the system starts fully labeled, i.e. $\mathbf{f}(0) = \mathbf{1}_n$, and we use the fact that $\mathbf{M}$ commutes with $\mathrm{e}^{\mathbf{M}t}$. Furthermore, $\mathrm{e}^{\mathbf{M}t}\mathbf{1}_n$ represents the labeling of the reduced system (i.e., the solution for $\mathbf{g} = 0$) and therefore $\mathbf{M}\mathrm{e}^{\mathbf{M}\tau}\mathbf{1}_n = \dot{\mathbf{f}}_r$. Finally, the solution can be written as:

$$\mathbf{f}(t) = \mathbf{f}_r(t) + \int_0^t \dot{\mathbf{f}}_r(t) f^\circ(t-\tau)d\tau = \mathbf{f}_r(t) + (\dot{\mathbf{f}}_r * f^\circ)(t)\,, \tag{S133}$$

It is, therefore, equivalent to the solution based on age distributions we got in Equation S128.

##### S1.9.1 Deconvolution of a single source described by a single exponential decay

If the source labeling is described by a single exponential decay, e.g. it is a single well-mixed pool fed by the external source: $f^\circ(t) = \mathrm{e}^{-\kappa^\circ t}$, there is a simple deconvolution method for finding the labeling function of the reduced system. First, we calculate the derivative of $f(t)$:

$$\begin{aligned} f(t) &= f_r(t) - (\dot{f}_r * f^\circ)(t) = f_r(t) - (\dot{f}_r * \mathrm{e}^{-\kappa^\circ t})(t) \\ \dot{f}(t) &= \dot{f}_r(t) - (\dot{f}_r * \dot{f}^\circ)(t) - \dot{f}_r(t) f^\circ(0) = \cancel{\dot{f}_r(t)} + \kappa^\circ(\dot{f}_r * \mathrm{e}^{-\kappa^\circ t})(t) - \cancel{\dot{f}_r(t)} = \kappa^\circ(\dot{f}_r * \mathrm{e}^{-\kappa^\circ t})(t) \end{aligned} \tag{S134}$$

where we use the general expression for the derivative of a convolution: $d(f*g)/dt = (f*(dg/dt))(t) + f(t)g(0)$.

Then, we can see that it is possible to use the derivative to eliminate the convolution term from $f(t)$:

$$f(t) + \frac{\dot{f}(t)}{\kappa^\circ} = f_r(t) - \cancel{(\dot{f}_r * \mathrm{e}^{-\kappa^\circ t})(t)} + \cancel{\frac{\kappa^\circ}{\kappa^\circ}(\dot{f}_r * \mathrm{e}^{-\kappa^\circ t})(t)} = f_r(t) \tag{S135}$$

##### S1.9.2 Example: 2-step irreversible pathway

As a demonstration of how to use Equation 17, we focus on a 2-step irreversible pathway:

$$\xrightarrow{\kappa^\circ} \underset{\downarrow}{S^\circ} \xrightarrow{\kappa^r} \underset{\downarrow}{S^r}\,. \tag{S136}$$

Since the first pool has only one input, its labeling will follow a simple exponential function, $f^\circ(t) = \mathrm{e}^{-\kappa^\circ t}$. Similarly, the "reduced" target system will be described by $f_r(t) = \mathrm{e}^{-\kappa^r t}$. Now we can use Equation 17 to find $f$:

$$f(t) = f^\circ(t) - (\dot{f}^\circ * f_r)(t) = \mathrm{e}^{-\kappa^\circ t} + \kappa^\circ \cdot (\mathrm{e}^{-\kappa^\circ t} * \mathrm{e}^{-\kappa^r t}) = \mathrm{e}^{-\kappa^\circ t} + \kappa^\circ \cdot \frac{\mathrm{e}^{-\kappa^\circ t} - \mathrm{e}^{-\kappa^r t}}{\kappa^r - \kappa^\circ} = \frac{\kappa^r\, \mathrm{e}^{-\kappa^\circ t}}{\kappa^r - \kappa^\circ} + \frac{\kappa^\circ\, \mathrm{e}^{-\kappa^r t}}{\kappa^\circ - \kappa^r}\,, \tag{S137}$$

which is the same solution as we got using a CM with the two pools (Equation S89, with replacing $\kappa_{e1} \to \kappa^\circ$ and $\kappa_{12} \to \kappa^r$).

Now we can apply the deconvolution method from Equation S135 on $f(t)$ to get the reduced form back:

$$f + \frac{\dot{f}}{\kappa^\circ} = \cancel{\frac{\kappa^r\, \mathrm{e}^{-\kappa^\circ t}}{\kappa^r - \kappa^\circ}} + \frac{\kappa^\circ\, \mathrm{e}^{-\kappa^r t}}{\kappa^\circ - \kappa^r} + \cancel{\frac{1}{\kappa^\circ}\frac{\kappa^r\,(-\kappa^\circ)\, \mathrm{e}^{-\kappa^\circ t}}{\kappa^r - \kappa^\circ}} + \frac{1}{\kappa^\circ}\frac{\kappa^\circ\,(-\kappa^r)\, \mathrm{e}^{-\kappa^r t}}{\kappa^\circ - \kappa^r} = (\kappa^\circ - \kappa^r)\frac{\mathrm{e}^{-\kappa^r t}}{\kappa^\circ - \kappa^r} = \mathrm{e}^{-\kappa^r t} \tag{S138}$$

##### S1.9.3 Example: a general system with a single entry state

A system with a single entry state where all downstream states are fed irreversibly from it and not from the external source:

$$\xrightarrow{\kappa^\circ} \underset{\downarrow}{S^\circ} \longrightarrow \underset{\downarrow}{Q^r}\,. \tag{S139}$$

i.e., this is a generalization of the previous case (S1.9.2), where the second step does not have to be a single state, but can represent a whole subsystem.

As we have shown (and also explained in Forney and Rothman [2014]), we can assume that the labeling function of the whole system can be described by a weighted sum of decaying exponential functions:

$$f(t) = \sum_{i=1}^{n} \delta_i \mathrm{e}^{-\kappa_i t} \tag{S140}$$

where $\kappa_i > 0$, $\delta_i \in \mathbb{R}$, and $\sum_i \delta_i = 1$. Again, we can apply Equation S135 to find the labeling of the "reduced" subsystem:

$$f_r(t) = f(t) + \frac{\dot{f}(t)}{\kappa^\circ} = \sum_{i=1}^{n} \delta_i \mathrm{e}^{-\kappa_i t} + \frac{1}{\kappa^\circ}\sum_{i=1}^{n} \delta_i(-\kappa_i)\mathrm{e}^{-\kappa_i t} = \sum_{i=1}^{n} \delta_i \left(1 - \frac{\kappa_i}{\kappa^\circ}\right)\mathrm{e}^{-\kappa_i t}\,. \tag{S141}$$

Since $\kappa^\circ$ is one of the decay rates that appear in the original sum (i.e. $\kappa^\circ = \kappa_i$ for some $i$), that exponent will drop out of the sum (because $1 - \kappa_i/\kappa^\circ = 0$), and we will be left with a weighted sum of $n-1$ exponents, as expected.

##### S1.9.4 Example: connecting two general systems together

To further generalize the previous example, we now consider a system with two sections, where the first one is the only input feeding the second one (irreversibly):

$$\longrightarrow \underset{\downarrow}{Q^{\circ}} \longrightarrow \underset{\downarrow}{Q^{r}}\,. \tag{S142}$$

$$\begin{aligned} f^{\circ}(t) &= \sum_{j=1}^{m} \alpha_j \mathrm{e}^{-\kappa_j^{\circ} t} \\ f_r(t) &= \sum_{i=1}^{n} \beta_i \mathrm{e}^{-\kappa_i^{r} t} \end{aligned} \tag{S143}$$

In this case, we cannot use the deconvolution method from Equation S135 since the input is not a single exponent. However, we can instead compute the convolution directly and then compare the different coefficients of the exponents in the sum:

$$\begin{aligned} f(t) = f^{\circ}(t) - (\dot{f}^{\circ} * f_r)(t) &= \sum_{j} \alpha_j \mathrm{e}^{-\kappa_j^{\circ} t} - \sum_{i,j} \beta_i \alpha_j \kappa_j^{\circ} (\mathrm{e}^{-\kappa_j^{\circ} t} * \mathrm{e}^{-\kappa_i^{r} t}) \\ &= \sum_{j} \alpha_j \mathrm{e}^{-\kappa_j^{\circ} t} - \sum_{i,j} \beta_i \alpha_j \kappa_j^{\circ} \left( \frac{\mathrm{e}^{-\kappa_j^{\circ} t}}{\kappa_i^{r} - \kappa_j^{\circ}} + \frac{\mathrm{e}^{-\kappa_i^{r} t}}{\kappa_j^{\circ} - \kappa_i^{r}} \right) \\ &= \sum_{j} \underbrace{\alpha_j \left( 1 - \sum_{i} \frac{\beta_i \kappa_j^{\circ}}{\kappa_i^{r} - \kappa_j^{\circ}} \right)}_{\gamma_j} \mathrm{e}^{-\kappa_j^{\circ} t} + \sum_{i} \underbrace{\left( \sum_{j} \frac{\beta_i \alpha_j \kappa_j^{\circ}}{\kappa_j^{\circ} - \kappa_i^{r}} \right)}_{\delta_i} \mathrm{e}^{-\kappa_i^{r} t} \\ &= \sum_{j} \underbrace{\alpha_j \left( 1 + \sum_{i} \frac{\beta_i}{1 - \kappa_i^{r}/\kappa_j^{\circ}} \right)}_{\gamma_j} \mathrm{e}^{-\kappa_j^{\circ} t} + \sum_{i} \underbrace{\beta_i \left( \sum_{j} \frac{\alpha_j}{1 - \kappa_i^{r}/\kappa_j^{\circ}} \right)}_{\delta_i} \mathrm{e}^{-\kappa_i^{r} t} \end{aligned} \tag{S144}$$

where we use the fact that $(\mathrm{e}^{-\alpha t} * \mathrm{e}^{-\beta t}) = (\beta - \alpha)\mathrm{e}^{-\alpha t} + (\alpha - \beta)\mathrm{e}^{-\beta t}$. Now, assume that we can find the values of $\{\gamma_j\}$, $\{\delta_i\}$, $\{\kappa_j^{\circ}\}$, and $\{\kappa_i^{r}\}$ by fitting data collected from the whole system. Then, we can deduce the values of $\{\beta_i\}$ using:

$$\beta_i = \delta_i \left( \sum_{j} \frac{\alpha_j}{1 - \kappa_i^{r}/\kappa_j^{\circ}} \right)^{-1} \tag{S145}$$

and we can see that indeed Equation S141 is a special case where $m = 1$ and $\alpha_j = 1$.

In summary, if we have functions for describing the whole system labeling as well as the labeling of the input subsystem:

$$\begin{aligned} f(t) &= \sum_{j=1}^{m} \gamma_j \mathrm{e}^{-\kappa_j^{\circ} t} + \sum_{i=1}^{n} \delta_i \mathrm{e}^{-\kappa_i^{r} t} \\ f^{\circ}(t) &= \sum_{j=1}^{m} \alpha_j \mathrm{e}^{-\kappa_j^{\circ} t} \end{aligned} \tag{S146}$$

then the reduced subsystem labeling will be:

$$f_r(t) = \sum_{i=1}^{n} \delta_i \left( \sum_{j=1}^{m} \frac{\alpha_j}{1 - \kappa_i^{r}/\kappa_j^{\circ}} \right)^{-1} \mathrm{e}^{-\kappa_i^{r} t} \tag{S147}$$

Note, that the $\{\gamma_j\}$ values are not used in the final result since these exponential terms drop out.

#### S1.10 Estimation of dynamic quantities

##### S1.10.1 Fitting a compartmental models to experimental labeling data

CMs can be used both to test hypotheses about metabolic networks and to realistically fit the experimental data in order to extract dynamic parameters of analyzed systems. In both cases, the experimental data ($\mathbf{f}(t_i)$ values measured at multiple time points – $\{t_i\}$ – and for one or more internal pool combinations) can be fitted with a non-linear least-squares solver, based on Equation 21:

$$\mathbf{M}^*, \mathbf{w}^* = \min_{\mathbf{M},\mathbf{w}} \sum_i \left(\mathbf{w}^\top \mathbf{f}(t_i) - \mathbf{w}^\top \mathrm{e}^{\mathbf{M}t_i}\mathbf{1}_n\right)^2 \tag{S148}$$

where $\mathbf{M}$ satisfies the conditions for compartmental matrices (namely, that none of the off-diagonal values are positive, and the sum over each row is non-positive, see Appendix S1.14), and $\mathbf{w} \in \mathbb{R}^n_+$ are the relative pool weights in the sample (and sum up to 1). If the data comprise more than one pool combination – $\{\mathbf{w}_j\}$ – minimization will be over the sum of all of them. Importantly, if the measured system is growing at a rate $\mu$, then $\mathbf{M}$ has to additionally satisfy the mass-balance constraints in Equation 24. We can show (Lemma S1.19 in the appendix) that given $\mathbf{M}$ and $\mu$, there exists a pool size vector $\mathbf{s}$ that satisfies the mass balance constraints if and only if the largest eigenvalue of $\mathbf{M}$ is smaller than $-\mu$. If the pool weights $\mathbf{w}$ reflect the real sizes of the metabolic pools $\mathbf{s}$, e.g., if the CM represents a whole system, then both $\mathbf{M}$ and $\mathbf{w}$ must be constrained by mass balance, according to Equation 24.

Beyond fitting the values of the system parameters quantitatively, one can also use the above procedure for model selection. For instance, the Akaike's Information Criterion Burnham and Anderson [2004] (AIC in short) can be used to guide the choice of appropriate CMs to represent experimental data, such as finding the most likely number of intermediate pools.

Once the CM is parametrized, all the dynamic parameters of the modeled metabolic system or any part thereof (Section 1.6) can be computed using the fitted CM parameters (Tables 1 and S1). In addition, individual influxes and effluxes can be reconstructed from the values in the fitted $\mathbf{M}^*$ and $\mathbf{w}^*$.

##### S1.10.2 Non-parametric estimation of mean age using trapezoid integration

Whereas CMs require making assumptions about the topology of the analyzed system, it is also useful to have nonparametric methods for evaluating the dynamic properties, even if their explaining power is reduced in comparison. For example, based on Equation 4, the mean age ($\bar{\mathcal{A}}$) is equal to the integral of the labeling curve and can be approximated simply by estimating the area under the curve using numerical methods, e.g. dividing it into trapezoids and calculating their total area. Similarly, based on Equation 9, we can evaluate metabolite decay rates by calculating the initial slope of the labeling curve (Appendix S1.10.3).

As we have seen in the previous sections, parametrizing a CM requires making assumptions about the topology of the system (e.g. the number of compartments and their connections) and often can lead to non-identifiable parameters. It can be, therefore, useful to have methods that require fewer assumptions, even if their explaining power is reduced in comparison. For example, based on Equation 4, we can see that the mean age can be directly calculated from $f(t)$ by integration, i.e. we can directly estimate $\bar{\mathcal{A}}$ using numerical integration methods and skip the need to fit the CM parameters (namely, $\mathbf{s}$ and $\mathbf{M}$).

Since we typically will only have a small finite set of time-course measurements of the labeling curve, there will be a need to interpolate them in order to get the full $f(t)$ trajectory. Furthermore, we will need to extrapolate it beyond the last time point (to integrate it to infinity). Fortunately, there is a simple solution that provides reasonable estimates, as long as the data is not too sparse.

We assume that the function is piecewise linear, that is, between every two measured data points $f(t)$ follows a straight line connecting them. For the range beyond the last time point, we simply assume that the slope $\dot{f}(t)$ remains the same as in the previous section (i.e. between the last two measured points). Then, the integral of $f(t)$ which is the area under the curve, will be approximated by the sum of the areas of all the trapezoids defined by this piece-wise linear function. The last section will actually be a triangle that will stretch until the function hits the x-axis. An illustrative example can be found in Figure S9A and in the code examples provided (see Materials and Methods).

Importantly, we have already seen that the age distribution is valid regardless of whether the system has delayed input and is not affected by growth dilution. Therefore, the non-parametric estimate of the mean age is a robust measure which can be used as-is, and without the need to ensure that constraints such as mass-balance (Equation 24 are satisfied.

##### S1.10.3 Non-parametric estimation of expected decay rate using the initial slope

Another useful dynamic quantity, which is yet simpler to calculate non-parametrically, is the expected decay rate. As we have seen in Equation 9, there is a simple relationship between $\bar{\kappa}$ and the initial slope of $f(t)$. Similarly, the expected decay rate in growing systems, can be calculated in the same way except for subtracting the growth rate (Equation S21). In figure S9B, one can find an example of how to estimate the decay rate using this method.

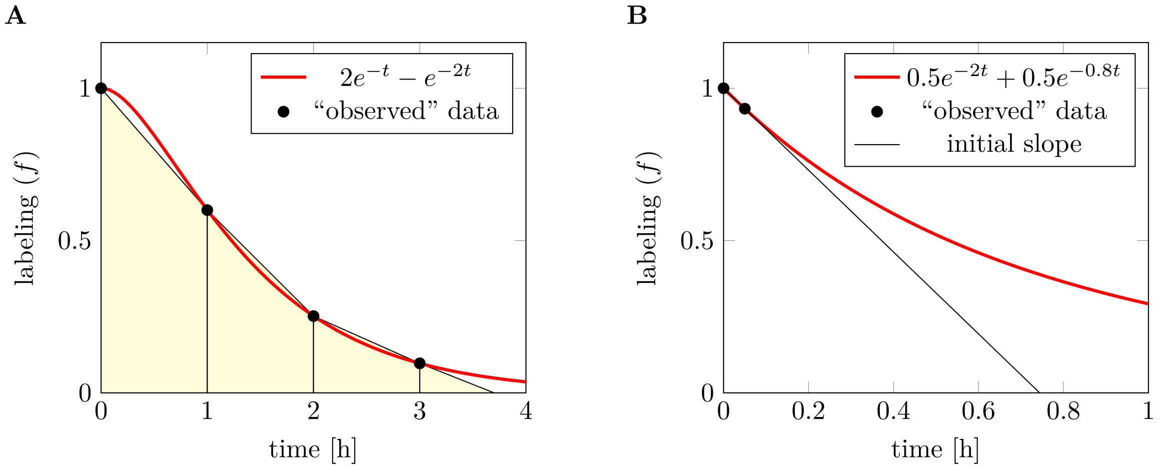


Figure S9: **Non-parametric estimation of mean age and expected decay rate.** The “observed” data points (black circles) were simulated using a CM, which is shown as a red curve. (A) Using the CM for the analytical integration yields $\bar{\mathcal{A}} = 1.5$ h. For the estimation of mean age without using the analytical function, we assume that the labeling curve is piecewise linear, and interpolate the points (thin black lines). Then, we can calculate the total area of the yellow trapezoids to get an estimate of the mean age. Here, the result would be 1.43 h, which is only $\sim 5\%$ error. As we have seen, mean age calculations don’t require direct labeling, and in this example the system is indeed has delayed input ($f(t)$ has an inflection point at $t = 1$). Therefore, the numerical method can be either provide an over- or under-estimate, depending on the shape of the curve and position of the measurements. (B) Using the CM to analytically calculate the expected decay rate yields $\bar{\kappa} = -\dot{f}(0) = 1.4\ h^{-1}$. The non-parametric method requires a measurement close enough to the labeling initiation time (i.e., where the labeled fraction is still very close to 1). Using the two marked points to calculate the slope gives an estimate of 1.35 $h^{-1}$, corresponding to an error of $\sim 4\%$. Since decay rate calculations require direct input, $f(t)$ must be convex, and (assuming there is no measurement noise) the numerical estimate will always be an underestimate of the actual decay rate. Moreover, this method is very sensitive to early-stage effects on labeling, such as precursor pools that are unaccounted for.

#### S1.11 Other mathematical derivation

##### S1.11.1 General solution for the overall labeling

To find a description for $f_{\mathbf{s}}$ as a function of time, we first calculate the time derivative:

$$\begin{aligned}\dot{f}_{\mathbf{s}} &= \frac{\mathbf{s}^\top \dot{\mathbf{f}}}{\mathbf{s}^\top \mathbf{1}_n} = \frac{\mathbf{s}^\top \mathbf{M}\mathbf{f} + \mathbf{s}^\top \mathbf{g}}{\mathbf{s}^\top \mathbf{1}_n} \\ &= \frac{\mathbf{s}^\top \mathrm{diag}\left(\mathbf{s}\right)^{-1}\left[\mathbf{V}^\top - \mathrm{diag}\left(\mathbf{V}^\top \mathbf{1}_n + \mathbf{Y}^\top \mathbf{1}_q\right)\right]\mathbf{f} + \mathbf{s}^\top \mathrm{diag}\left(\mathbf{s}\right)^{-1}\mathbf{Y}^\top \mathbf{h}}{\mathbf{s}^\top \mathbf{1}_n} \\ &= \frac{\mathbf{1}_n^\top\left[\mathbf{V}^\top - \mathrm{diag}\left(\mathbf{V}^\top \mathbf{1}_n + \mathbf{Y}^\top \mathbf{1}_q\right)\right]\mathbf{f} + \mathbf{1}_n^\top \mathbf{Y}^\top \mathbf{h}}{\mathbf{s}^\top \mathbf{1}_n} \\ &= -\frac{\left[\mathbf{1}_q^\top \mathbf{Y} + \mathbf{1}_n^\top \mathbf{V} - \mathbf{1}_n^\top \mathbf{V}^\top\right]\mathbf{f} + \mathbf{1}_n^\top \mathbf{Y}^\top \mathbf{h}}{\mathbf{s}^\top \mathbf{1}_n}.\end{aligned} \tag{S149}$$

where we used the formula for the derivative of $\dot{\mathbf{f}}$ from Equation S63, and the fact that $\forall \mathbf{x}\ \ \mathbf{1}_n^\top \mathrm{diag}\left(\mathbf{x}\right) = \mathbf{x}^\top$. Looking more closely at this result, we note that the denominator is the total pool size ($\mathbf{s}^\top \mathbf{1}_n = \sum_i s_i$). The last term in the numerator ($\mathbf{1}_n^\top \mathbf{Y}^\top \mathbf{h} = \sum_i \sum_k h_k y_{k\to i}$) is the total rate at which labeled molecules enter into the system. The other terms in the numerator are the total rate at which labeled molecules leave the system, which we can prove by using the flux-balance equation for each $S_i$:

$$-\left[\mathbf{1}_q^\top \mathbf{Y} + \mathbf{1}_n^\top \mathbf{V} - \mathbf{1}_n^\top \mathbf{V}^\top\right]\mathbf{f} = \sum_i f_i \left(\sum_k y_{k\to i} + \sum_j v_{i\to j} - \sum_j v_{j\to i}\right) = \sum_i f_i v_{i\to\emptyset} \tag{S150}$$

This is a useful way to think about the system, as a single state with one input ($\sum_i \sum_k h_k y_{k\to i}$), one output ($\sum_i f_i v_{i\to\emptyset}$), and one combined pool ($\sum_i s_i$). If, as before, we assume that $\mathbf{h}^\circ = \mathbf{0}_q$ (i.e. anything entering the system is labeled) we can drop the last term in the numerator of Equation S149:

$$\dot{f}_{\mathbf{s}} = -\frac{\left[\mathbf{1}_q^\top \mathbf{Y} + \mathbf{1}_n^\top \mathbf{V} - \mathbf{1}_n^\top \mathbf{V}^\top\right]\mathbf{f}}{\mathbf{s}^\top \mathbf{1}_n} \tag{S151}$$

**Internalizing external fluxes** Consider a system with no external fluxes ($\mathbf{Y} = 0$). Instead, we can imagine a subset of states that have only outgoing irreversible fluxes. We set their labeling pattern once at time $t = 0$ and note that their labeling will remain constant over time since nothing is coming into them to change it. Another way to see this, is realizing that the column in $\mathbf{M}$ corresponding to such a state $i$ will be all 0s (since $\forall j \; v_{j\to i} = 0$). In the exponent $e^{\mathbf{M}t}$, it will become a unit vector (1 on the diagonal and 0s everywhere else). Therefore, the value of $f_i(t)$ will simply be $f_i^\circ$ for any $t > 0$.

To simulate a wash-out experiment, we can define $S_0$ as the source of label, which switches off at $t = 0$, while all other states start completely labeled – i.e. $\mathbf{f}^\circ = (0, 1, \ldots, 1)$. This is mathematically equivalent to having $S_0$ removed from the system and representing all the fluxes that go from it to other states by external fluxes (instead of being in the first row of $\mathbf{V}$, they will be in $\mathbf{Y}$).

##### S1.11.2 An expression for the vector of output exchange fluxes

To prove that $-\mathbf{s}^\top\mathbf{M}$ is the vector of output exchange fluxes, we use the definition of $\mathbf{M}$ to derive an expression based directly on the fluxes:

$$\begin{aligned}
-\mathbf{s}^\top\mathbf{M} &= -\mathbf{s}^\top \mathrm{diag}\left(\mathbf{s}\right)^{-1}\left[\mathbf{V}^\top - \mathrm{diag}\left(\mathbf{V}^\top\mathbf{1}_n + \mathbf{Y}^\top\mathbf{1}_q\right)\right] \\
&= -\mathbf{1}_n\left[\mathbf{V}^\top + \mathrm{diag}\left(\mathbf{V}^\top\mathbf{1}_n + \mathbf{Y}^\top\mathbf{1}_q\right)\right] \\
&= -\mathbf{1}_n^\top\mathbf{V}^\top + \mathbf{1}_n^\top\mathrm{diag}\left(\mathbf{V}^\top\mathbf{1}_n\right) + \mathbf{1}_n^\top\mathrm{diag}\left(\mathbf{Y}^\top\mathbf{1}_q\right) \\
&= -\mathbf{1}_n^\top\mathbf{V}^\top + \left(\mathbf{V}^\top\mathbf{1}_n\right)^\top + \left(\mathbf{Y}^\top\mathbf{1}_q\right)^\top \\
&= -\mathbf{1}_n^\top\mathbf{V}^\top + \mathbf{1}_n^\top\mathbf{V} + \mathbf{1}_q^\top\mathbf{Y}
\end{aligned} \tag{S152}$$

where we used the fact that $\forall\mathbf{x} \;\; \mathbf{1}_n^\top\mathrm{diag}\left(\mathbf{x}\right) = \mathbf{x}^\top$.

Now, we continue by explicitly writing the expression for the $i$th row in $-\mathbf{s}^\top\mathbf{M}$:

$$-\sum_j v_{i\to j} + \sum_j v_{j\to i} + \sum_j y_{j\to i} \tag{S153}$$

and since we assume there is a flux-balance for each state (i.e. all fluxes coming into $S_i$ equal all outgoing fluxes), we can replace $\sum_j v_{j\to i} + \sum_j y_{j\to i}$ with $\sum_j v_{i\to j} + v_{i\to\emptyset}$, which leaves us with $v_{i\to\emptyset}$.

##### S1.11.3 Mathematical proofs regarding the labeling curve

**Lemma S1.4.** *Let $g(x)$ be a real function, which satisfies:*

$$\lim_{x\to 0} x\; g(x) = 0 \qquad \textit{and} \qquad \lim_{x\to\infty} x\; g(x) = 0$$

*Then the following holds:*

$$\int_0^\infty \frac{dg}{dx}\; x\; dx = -\int_0^\infty g\; dx$$

*Proof.* Using integration by parts:

$$\int_0^\infty \frac{dg}{dx}\; x\; dx = x\; g(x)\Big|_0^\infty - \int_0^\infty g(x)dx$$

and since the two limits of $x\; g(x)$ are equal to 0, the first term can be dropped. ■

**Corollary S1.4.1.** Let $X$ be a random variable which accepts only positive values ($x \geq 0$), and satisfies

$$\lim_{x\to\infty} x\left[1 - P_X(x)\right] = 0\,.$$

Then

$$\mathbb{E}[X] = \int_0^\infty \left[1 - P_X(x)\right] dx\,.$$

*Proof.* Define $g(x) \equiv 1 - P_X(x)$. Since $P_X(0) = 0$, we can see that $g(0) = 1$ and thus $\lim_{x\to 0} x\; g(x) = 0$. The other limit is satisfied directly from the condition of the corollary. Therefore:

$$\begin{aligned}
\mathbb{E}[X] \;\; &\equiv \int_0^\infty p_X(x)\; x\; dx = -\int_0^\infty \frac{dg}{dx}\; x\; dx = -x\; g(x)\Big|_0^\infty + \int_0^\infty g dx \\
&= -\cancel{\lim_{x\to\infty} x\; g(x)} + \cancel{\lim_{x\to 0} x\; g(x)} + \int_0^\infty g\; dx = \int_0^\infty \left[1 - P_X(x)\right] dx
\end{aligned}$$

■

##### S1.11.4 Mathematical proofs involving compartmental matrices

Consider a real square matrix $\mathbf{A} \in \mathbb{R}^{n\times n}$, $\mathbf{A} = (a_{ij})$.

**Definition S1.5.** Let $\lambda_i \in \mathbb{C}$ be the eigenvalues of $\mathbf{A}$.
The *spectral radius* of $\mathbf{A}$, denoted $\rho(\mathbf{A})$, is the largest modulus of its eigenvalues:

$$\rho(\mathbf{A}) = \max_i \{|\lambda_i|\} .$$

The *spectral abscissa* of $\mathbf{A}$, denoted $\eta(\mathbf{A})$, is the largest real part of all of its eigenvalues:

$$\eta(\mathbf{A}) = \max_i \{Re(\lambda_i)\} .$$

**Definition S1.6.** $\mathbf{A}$ is called *non-negative* if all of its elements are non-negative: $a_{ij} \geq 0,\ \forall i, j$.

**Lemma S1.7.** *Let $\mathbf{A} \in \mathbb{R}^{n\times n}$ non-negative. Then:*

$$\rho(\mathbf{A}) \leq \max_i \sum_j a_{ij}$$

*Proof.* Before proving the lemma, we consider Gershgorin's circle theorem Geršgorin [1931]. Let $\mathbf{A}$ be a complex square matrix and define $R_i \equiv \sum_{j\neq i} |a_{ij}|$, i.e. the sum of the absolute values of the non-diagonal entries in the $i$-th row. Let $D(a_{ii}, R_i) \subseteq \mathbb{C}$ be a closed disc centered at $a_{ii}$ with radius $R_i$. Such a disc is called a *Gershgorin disc.*

**Gershgorin Circle Theorem**: Every eigenvalue of $\mathbf{A}$ lies within at least one of the Gershgorin discs $D(a_{ii}, R_i)$.
Since $\rho(\mathbf{A})$ is the modulus of one of the eigenvalues (say $|\lambda_j|$) and that eigenvalue lies within one of the discs (say $R_i$), we can write:

$$\sum_{j\neq i} |a_{ij}| = R_i \geq |\lambda_j - a_{ii}| \geq |\lambda_j| - |a_{ii}| = \rho(\mathbf{A}) - |a_{ii}|$$

where we use the triangle inequality. Since all values in $\mathbf{A}$ are non-negative and real, we can drop the $|\cdot|$ for them:

$$\sum_{j\neq i} a_{ij} \geq \rho(\mathbf{A}) - a_{ii} \quad \Rightarrow \quad \rho(\mathbf{A}) \leq \sum_j a_{ij} .$$

As this is true for row $i$, it will definitely be true for the maximum of all rows. ■

**Lemma S1.8.** *Let $\mathbf{A} \in \mathbb{R}^{n\times n}$ non-negative. Then, the spectral radius $\rho(\mathbf{A})$ is an eigenvalue of $\mathbf{A}$, and corresponds to a non-zero eigenvector, $\mathbf{x} \geq \mathbf{0}_n$.*

*Proof.* The proof is a corollary of the *Perron-Frobenius theorem* and appears in Theorem 2.20 from Varga [2000]. ■

**Definition S1.9.** $\mathbf{A}$ is called *Metzler* if all of its elements are non-negative, except for those on the main diagonal: $a_{ij} \geq 0,\ \forall i \neq j$.

**Definition S1.10.** A Metzler matrix is called *strict* if it contains at least one strictly negative entry (which must be on the main diagonal).

**Lemma S1.11.** *If $\mathbf{B} \in \mathbb{R}^{n\times n}$ is strict Metzler, then*

$$\rho(\mathbf{B} + h\mathbf{I}_n) = \eta(\mathbf{B}) + h$$

*for any $h > 0$ such that $\mathbf{B} + h\mathbf{I}_n$ is non-negative.*

*Proof.* This appears as lemma 1 in Cvetković [2019]. ■

**Lemma S1.12.** *If $\mathbf{B} \in \mathbb{R}^{n\times n}$ is strict Metzler, then it has a non-zero eigenvector, $\mathbf{x} \geq \mathbf{0}_n$, whose eigenvalue is $\eta(\mathbf{B})$.*

*Proof.* Let $h > 0$ be large enough so that $\mathbf{B} + h\mathbf{I}_n$ is non-negative. From lemma S1.8 we know that it has a non-zero eigenvector, $\mathbf{x} \geq \mathbf{0}_n$, whose eigenvalue is $\rho(\mathbf{B} + h\mathbf{I}_n)$ and from lemma S1.11 we can replace $\rho(\mathbf{B} + h\mathbf{I}_n)$ with $\eta(\mathbf{B}) + h$. Therefore:

$$\begin{aligned}(\mathbf{B} + h\mathbf{I}_n)\mathbf{x} &= (\eta(\mathbf{B}) + h)\mathbf{x} \\ \mathbf{B}\mathbf{x} + h\mathbf{x} &= \eta(\mathbf{B})\mathbf{x} + h\mathbf{x} \\ \mathbf{B}\mathbf{x} &= \eta(\mathbf{B})\mathbf{x}\end{aligned}$$

■

**Lemma S1.13.** *If $\mathbf{B} \in \mathbb{R}^{n\times n}$ is strict Metzler and given $\mu \in \mathbb{R}$. Then:*

1. *If $\mu \geq \eta(\mathbf{B})$, there exists a non-zero eigenvector, $\mathbf{x} \geq \mathbf{0}_n$, where $\mathbf{B}\mathbf{x} \leq \mu\mathbf{x}$*
2. *If $\mu < \eta(\mathbf{B})$, there is no strictly positive vector $\mathbf{u}$ which satisfies $\mathbf{B}\mathbf{u} \leq \mu\mathbf{u}$*

*Proof.* Proving statement 1 follows directly from lemma S1.12 by choosing $\mathbf{x}$ as the eigenvector: $\mathbf{Bx} = \eta(\mathbf{B})\mathbf{x} \leq \mu\mathbf{x}$. For statement 2 we will use a proof by contradiction. Assume that there is such a vector $\mathbf{u}$. We again define $\mathbf{x}$ as the eigenvector from lemma S1.12, but for the transposed matrix $\mathbf{B}^\top$, so that $\mathbf{B}^\top\mathbf{x} = \eta(\mathbf{B}^\top)\mathbf{x}$. Now we can write:

$$\mu\mathbf{u}^\top\mathbf{x} \geq (\mathbf{Bu})^\top\mathbf{x} = \mathbf{u}^\top\mathbf{B}^\top\mathbf{x} = \mathbf{u}^\top\eta(\mathbf{B}^\top)\mathbf{x} = \mathbf{u}^\top\eta(\mathbf{B})\mathbf{x} = \eta(\mathbf{B})\mathbf{u}^\top\mathbf{x}$$

Since $\mathbf{x} \geq \mathbf{0}_n$, $\mathbf{u} > \mathbf{0}_n$, and $\mathbf{x}$ is non-zero, we can be sure that their inner product is strictly positive: $\mathbf{u}^\top\mathbf{x} > 0$. Therefore, we can eliminate the inner product from both sides of the inequality and get:

$$\mu \geq \eta(\mathbf{B})$$

which contradicts the assumption of statement 2. ∎

**Definition S1.14.** A matrix $\mathbf{B} \in \mathbb{R}^{n\times n}$ is called *compartmental* if it is strict Metzler and for each row $i$, $-b_{ii} \geq \sum_{j\neq i} b_{ij}$. $\mathbf{B} \in \mathbb{R}^{n\times n}$ is called *strict compartmental* if the inequality is strict for all of the rows.

**Lemma S1.15.** *If $\mathbf{B} \in \mathbb{R}^{n\times n}$ is (strict) compartmental then $\eta(\mathbf{B})$ is (strictly) negative.*

*Proof.* Define the non-negative matrix $\mathbf{A} = \mathbf{B} + h\mathbf{I}_n$. We can see that

$$\eta(\mathbf{B}) = \rho(\mathbf{A}) + h \leq \max_i \sum_j a_{ij} - h = \max_i \sum_j b_{ij} + h - h = \max_i \left( b_{ii} + \sum_{j\neq i} b_{ij} \right) \leq 0$$

where for the first step we use lemma S1.11 and for the second step – lemma S1.8. For the strict compartmental case, the last inequality will be strict. ∎

**Lemma S1.16.** *If $\mathbf{B} \in \mathbb{R}^{n\times n}$ is (strict) compartmental then $\mathbf{B1}_n$ is (strictly) negative.*

*Proof.* The value of the $i$-th element in $\mathbf{B1}_n$ is:

$$\sum_j b_{ij} = b_{ii} + \sum_{j\neq i} b_{ij} \leq b_{ii} - b_{ii} = 0$$

For the strict compartmental case, the inequality will be strict. ∎

**Lemma S1.17.** *If $\mathbf{B} \in \mathbb{R}^{n\times n}$ is compartmental then there exists a non-zero eigenvector, $\mathbf{x} \geq \mathbf{0}_n$, where $\mathbf{Bx} \leq \mathbf{0}_n$.*

*Proof.* From lemma S1.15 we know that $\eta(\mathbf{B}) \leq 0$. If we choose $\mu = 0 \geq \eta(\mathbf{B})$ then we can apply statement 1 of lemma S1.13 which proves that claim. ∎

**Lemma S1.18.** *For any compartmental $\mathbf{M}$-matrix, there exists a non-zero state-size vector $\mathbf{s} \geq \mathbf{0}_n$ and an efflux vector $\mathbf{v}_\emptyset \geq \mathbf{0}_n$ which together satisfy the mass-balance constraints, i.e.*

$$\forall i \quad \underbrace{s_i\kappa^*_{ei}}_{v_{ei}} + \sum_{j\neq i} \underbrace{s_i\kappa_{ji}}_{v_{ji}} = \sum_{j\neq i} \underbrace{s_j\kappa_{ij}}_{v_{ij}} + v_{i\emptyset} \tag{S154}$$

*where the left-hand side is the sum of incoming fluxes into compartment $S_i$ and the right-hand side is the corresponding sum of outgoing fluxes.*

*Proof.* Since $\mathbf{M}^\top$ qualifies for the conditions of lemma S1.17, we know that there exists a non-zero eigenvector, $\mathbf{x} \geq \mathbf{0}_n$, which satisfies

$$\mathbf{M}^\top\mathbf{x} \leq \mathbf{0}_n \tag{S155}$$

We define $\mathbf{s} \equiv \mathbf{x}^\top$ and $\mathbf{v}_\emptyset \equiv -\mathbf{s}^\top\mathbf{M}$. We can see that $\mathbf{s}$ and $\mathbf{v}_\emptyset$ are non-negative. So, we are only left to show that the mass-balance constraints are met. First, we rearrange them:

$$\forall i \quad -s_i \left( \underbrace{\kappa^*_{ei}}_{v_{ei}} + \sum_{j\neq i} \underbrace{\kappa_{ji}}_{v_{ji}} \right) + \sum_{j\neq i} \underbrace{s_j\kappa_{ij}}_{v_{ij}} = -v_{i\emptyset} \tag{S156}$$

Based on the definition of $\mathbf{M}$ (see Appendix S1.6.2), the left-hand side of this equation is the $i$-th element of $\mathbf{s}^\top\mathbf{M}$, and the right-hand side is the $i$-th element of $\mathbf{v}_\emptyset$. Therefore, by its definition, $\mathbf{s}^\top\mathbf{M} = -\mathbf{v}_\emptyset$ proves the lemma. ∎

**Lemma S1.19.** *For any compartmental* $\mathbf{M}$*-matrix and growth rate* $\mu \leq -\eta(\mathbf{M})$*, there exists a non-zero state-size vector* $\mathbf{s} \geq \mathbf{0}_n$ *and an efflux vector* $\mathbf{v}_\emptyset \geq \mathbf{0}_n$ *which together satisfy the mass-balance constraints under a system with growth rate* $\mu$*, i.e.*

$$\forall i \quad \underbrace{s_i \kappa^*_{ei}}_{v_{ei}} + \sum_{j \neq i} \underbrace{s_i \kappa_{ji}}_{v_{ji}} = \sum_{j \neq i} \underbrace{s_j \kappa_{ij}}_{v_{ij}} + v_{i\emptyset} + \mu s_i \tag{S157}$$

*where the left-hand side is the sum of incoming fluxes into compartment* $S_i$ *and the right-hand side is the corresponding sum of outgoing fluxes and dilution by growth.*

*Proof.* The proof follows almost exactly the one from lemma S1.18, except that we use lemma S1.13 to find $\mathbf{x}$ where $\mathbf{M}^\top \mathbf{x} \leq -\mu \mathbf{x}$ and we define $\mathbf{s} \equiv \mathbf{x}^\top$, $\mathbf{v}_\emptyset \equiv -\mathbf{s}^\top(\mathbf{M} + \mu \mathbf{I}_n) \geq \mathbf{0}_n$. Therefore, we can see that $\mathbf{s}^\top \mathbf{M} = -\mu \mathbf{s}^\top - \mathbf{v}_\emptyset$ directly translate to the mass-balance constraints. ■

**Lemma S1.20.** *If* $\mathbf{B}$ *is a Metzler matrix, then* $\mathrm{e}^{\mathbf{B}}$ *is non-negative.*

*Proof.* We will use the formula

$$\mathrm{e}^{\mathbf{B}} = \lim_{k \to \infty} \left( \mathbf{I}_n + \frac{\mathbf{B}}{k} \right)^k .$$

The off-diagonal values of $\mathbf{I}_n + \frac{\mathbf{B}}{k}$ are always non-negative (because $\forall i \neq j \;\; b_{ij}/k \geq 0$). Define $K = -\min_i b_{ii}$. For any $k > K$, the diagonal values $(1 + b_{ii}/k)$ will be positive – and thus all values of $\mathbf{I}_n + \frac{\mathbf{B}}{k}$ will be non-negative. So, it is clear that also $\left(\mathbf{I}_n + \frac{\mathbf{B}}{k}\right)^k$ will contain only non-negative values. Therefore, this holds true also at the limit $k \to \infty$, and thus $\mathrm{e}^{\mathbf{B}}$ is a non-negative matrix. ■

**Corollary S1.20.1.** If $\mathbf{B}$ is a compartmental matrix, then $\mathbf{B}\mathrm{e}^{\mathbf{B}t}\mathbf{1}_n \leq \mathbf{0}_n$, $\forall t > 0$.

*Proof.* Every matrix commutes with its exponent, i.e. $\mathbf{A}\mathrm{e}^{\mathbf{A}} = \mathrm{e}^{\mathbf{A}}\mathbf{A}$, and therefore we get that:

$$\mathbf{B}\mathrm{e}^{\mathbf{B}t}\mathbf{1}_n = \mathrm{e}^{\mathbf{B}t}\mathbf{B}\mathbf{1}_n .$$

Since $\mathbf{B}t$ is itself also a Metzler matrix, $\mathrm{e}^{\mathbf{B}t}$ is non-negative (lemma S1.20). $\mathrm{e}^{\mathbf{B}t}\mathbf{B}\mathbf{1}_n$ is a product of a non-negative matrix and a non-positive vector (from lemma S1.16), thus it is itself a non-positive vector. ■

**Lemma S1.21.** *For any matrix* $\mathbf{B}$*, with* $\eta(\mathbf{B}) < 0$*, and any* $k \in \mathbb{R}$*:*

$$\lim_{t \to \infty} t^k \mathrm{e}^{\mathbf{B}t} = \mathbf{0}_n$$

*Proof.* $\eta(\mathbf{B}) < 0$ means that all of its eigenvalues have negative real parts, i.e. $\mathbf{B}$ is Hurwitz stable. Therefore, we know that there exists positive numbers $\beta$ and $\alpha$ such that $||\mathrm{e}^{\mathbf{B}t}|| \leq \beta \mathrm{e}^{-\alpha t}$ for all $t$. We can now write:

$$\lim_{t \to \infty} ||t^k \mathrm{e}^{\mathbf{B}t}|| = \lim_{t \to \infty} t^k ||\mathrm{e}^{\mathbf{B}t}|| \leq \beta \lim_{t \to \infty} t^k \mathrm{e}^{-\alpha t} = 0$$

which proves the lemma. ■

Importantly, for compartmental matrices, we can confidently assume that $\eta(\mathbf{M}) < 0$. Although theoretically possible, systems with $\eta(\mathbf{M}) = 0$ are not realistic in real-world settings because that means they have a cycle flux without any degradation or dilution. Moreover, systems growing at rate $\mu$ will be described by matrices that have $\eta(\mathbf{M}) \leq -\mu$ which must be negative.

### S2 Extended materials and methods

#### S2.1 Yeast strains and cell culture conditions

Dynamic labeling assays were performed using a haploid BY4742 strain (*MATα his3Δ1 leu2Δ0 lys2Δ0 ura3Δ0*) auxotrophic for lysine. Cell cultures were grown at 30°C or 37°C in standard synthetic complete medium (6.7 g/L yeast nitrogen base without amino acids, ammonium sulfate as a nitrogen source, 2% glucose, necessary amino acid supplements) and supplemented at 25 mg/L (fc) with normal $[^{12}C_6,^{14}N_2]$ L-lysine or heavy $[^{13}C_6,^{15}N_2]$ L-lysine isotopomer (Silantes), referred to as "light medium" and "heavy medium", respectively.

Before the labeling pulse, yeast cultures were propagated for a minimum of 16 hours in light medium at respective temperatures until $OD_{600}$ 0.4-0.5 and pulsed using heavy culture medium pre-warmed at the same temperature. Before inoculation in heavy medium, the light medium was removed by filtration on a 0.8 µm nitrocellulose membrane (Whatmann cat# 10565361) followed by a brief wash with 25 ml of pre-warmed heavy medium on the membrane.

Biomass growth was monitored throughout the duration of the labeling pulse by measuring cell culture turbidity at 600 nm ($OD_{600}$). To ensure exponential growth, for the duration of the labeling pulse, cell cultures were maintained at $OD_{600}$ in the 0.5-0.8 range by periodic dilution with heavy pre-warmed medium. $OD_{600}$ measurements were taken before sample collection and after dilution. The growth dilution rates were then calculated by linear regression of the log-transformed biomass growth.

Culture samples were harvested at different time points after the labeling pulse by filtration. The samples corresponding to time point zero were collected immediately before the start of labeling on a nitrocellulose membrane, briefly washed on the membrane with distilled water and frozen in liquid nitrogen. The rest of the labeling time-course samples were collected similarly at the respective time points.

#### S2.2 In-gel tryptic digestion

The frozen yeast pellets were resuspended in 1 ml of Lysis Buffer (50 mM HEPES pH 7.9, 150 mM NaCl, 2 mM EDTA, 5% v/v glycerol, 1 mM DTT, protease inhibitor cocktail (Merck, P8215) and transferred to 2 ml screw-cap micro tubes (Sarstedt Inc) prefilled with approximately 1 mL of 0.5 mm glass beads (Biospec products). To avoid air inclusions, the tubes were spun down and filled with the Lysis Buffer. Cells were lysed by four one-minute cycles at 3500 oscillations per minute in a FastPrep-24 Classic grinder (MP Biomedicals), with 1-minute cooling intermissions in an ice-water mixture between cycles. Cell lysates were clarified by 30 s centrifugation at 10,000 RPM. 25 µl of the supernatant was mixed with 25 µl 2x SDS loading buffer and electrofocussed by SDS-PAGE in a stacking gel. The gel was fixed (50% (v/v) methanol, 10% (v/v) acetic acid, 40% (v/v) water) for 15-25 minutes. Proteins were visualized by SimplyBlueTM SafeStain staining (Invitrogen, LC6060).

The excised gel areas were processed with a standard in-gel tryptic digestion protocol: All aqueous solutions were prepared in 100 mM ammonium bicarbonate. Proteins were reduced with 6.5 mM DTT for 1 h at 60°C, alkylated with 54 mM iodoacetamide for 30 min in the dark at room temperature, digested with 1.25 µg of sequencing grade porcine trypsin (Promega, V5113) at 37°C for 18 h. The tryptic peptides were desalted in Oasis HLB 96-well µ Elution Plate according to manufacturer protocol and recovered in 15-25 µl of Buffer A (0.5% (v/v) formic acid) supplemented with 1:50 (v/v) of Biognosis iRT peptides. The samples were transferred to MS vials for further mass spectrometric analysis.

#### S2.3 Mass spectrometry acquisition

About 1 $\mu$g of tryptic peptides dissolved in 2% acetonitrile (ACN), 0.5% formic acid (FA), were injected into an Ultimate 3000 RSLC system (Thermo Scientific, Sunnyvale, California, USA) connected online to an Orbitrap Eclipse Tribrid mass spectrometer (Thermo Scientific, Bremen, Germany) equipped with EASY-spray nano-electrospray ionization source (Thermo Scientific). The sample was loaded and desalted on a pre-column (Acclaim PepMap 100, 2cm x 75$\mu$m ID nanoViper column, packed with 3$\mu$m C18 beads) at a flow rate of 5 $\mu$l/min for 5 min with 0.1% trifluoroacetic acid.

Peptides were separated during a biphasic ACN gradient from two nanoflow UHPLC pumps (flow rate of 200 nl/min) on a 50 cm analytical column (EASY-Spray™ ES903, 15 cm x 75 µm C18, operated at 45°C). Solvent A and B were 0.1% FA (vol/vol) in water and 100% ACN, respectively. The gradient composition was 5%$B$ during trapping (5 min) followed by $5-7\%B$ over 1 min, $7-20\%$B over the next 111 min, $20-32\%$B over the next 30 min, and $32-85\%B$ over 3 min. Isocratic flow was maintained for 10 minutes, followed by $85-5\%B$ over the next 5 minutes. During this process, the injection valve was switched back to load position (6 1) at 153 min, and inject (1 2) at 157 min, to flush the column and valves. The analytical program was run for total of 2.5 hours at 5%$B$ (flow rate 200 nl/min). The analytical method was followed by a 15-minute injection valve wash step (flow rate 200 nl/min) at 5%$B$. The spray and ion-source parameters were as follows. Ion spray voltage = 1800V, no sheath and auxiliary gas flow, and capillary temperature = 275°C. The instrument was controlled through Thermo Scientific SII for Xcalibur.

The mass spectrometer was operated in a DIA-mode (data-independent-acquisition) to automatically switch between one full scan MS and MS/MS acquisition of 41 mass segments with varying window sizes and 1 m/z overlap. Instrument control was managed through Orbitrap Eclipse Tune and Xcalibur. MS spectra were acquired in the scan range 350-1150 m/z with resolution R = 120 000 at m/z 200, automatic gain control (AGC) target of $8 \times 10^5$ and a maximum injection time (IT) set to 246 ms. All ions in the m/z window were sequentially isolated to a target value (AGC) of $1 \times 10^5$ and a maximum IT of 66 ms in the C-trap before HCD fragmentation (Higher-Energy Collision Dissociation). Fragmentation was performed with a normalized collision energy (NCE) of 30%, and fragments were detected in the Orbitrap at a resolution of 30 000 at m/z 200. Lock-mass internal calibration was not enabled.

#### S2.4 Analysis of mass spectrometry data

Unlabeled spectral libraries were generated with FragPipe 23.0-build7 essentially as described in Yu et al. [2023] based on the total of 18 DDA runs of the full yeast lysate fractionated with SDS-PAGE. In short, the raw files were converted to mzML with MSConvert (Proteowizard v 3.0.24313-968a764). The mzML files were searched with

MSFragger (version 4.1) using the SGD database containing 6,713 proteins plus one protein entry for the concatenated sequence of Biognosis iRT peptides. For MSFragger, the reversed decoy sequences were generated and appended to the target database. FragPipe was used to process the outputs of MSFragger to perform MSBooster (v1.2.64) deep-learning-based rescoring, Percolator (version 3.07.1) PSM re-ranking, ProteinProphet protein grouping, and Philosopher (version 5.1.1-RC16) FDR filtering. The "Default" workflow was applied with the adjusted maximum allowed 2 missed cleavages. Acetylation of the protein N-termini and oxidation of methionine were set as variable modifications, and carboxymethylation of cysteine was set as a fixed modification. Libraries were exported as .parquet files and further used in DIA-NN 2.1.0 Demichev et al. [2020] to analyze the pulse SILAC time courses (both 30°C and 37°C) at the same time. SILAC workflow was implemented by assigning the mass modification of the labeled lysine channel *in silico* to existing empirically derived unlabeled spectral libraries. The rest of the DIA-NN settings were default except "cross-run normalization", which was turned off.

#### S2.5 Quantification of protein labeling and abundance

Protein labeling quantification was performed based on the DIA-NN report (see above). The data were filtered to remove contaminants and non-proteotypic peptides. The dataset was split by channel (Light and Heavy), and only precursors containing a single lysine residue were retained. Precursor-level labeling was calculated as the fraction of Light intensity relative to the total signal per precursor:

$$\text{Labeling} = \frac{L}{H+L}$$

Precursors lacking detectable intensity in either channel were excluded. Protein-level labeling was obtained by aggregating precursor values per protein group (PG) using the median.

Relative protein abundance was derived from the same DIA-NN reports as used for labeling quantification. After splitting the data by channel, missing channel intensities were imputed as zero. For each precursor, the sum of Light and Heavy intensities ($H + L$) was calculated and subsequently aggregated by summation across all precursors within a protein group (PG). Protein-group intensities were then normalized by dividing by the total precursor intensity of the run. Finally, values were $\log_2$-transformed and used as run-normalized relative abundances.

#### S2.6 Determination of free lysine labeling dynamics

Mid-log phase cultures were grown in light medium at 30 or 37°C, harvested by filtration (WCN membrane filters, 0.8 µm, Whatman), washed on the filter, and transferred to pre-warmed heavy lysine medium. Immediately thereafter, the initial culture turbidity was measured at 600 nm. Following medium exchange, culture samples were collected at 5, 10, 20, 30, 45, 60, and 90 minutes to determine turbidity and harvest cells for the extraction of soluble lysine. The samples were spun for 1 min at 4K rpm; the cell pellets were washed twice with 1 ml of MS-grade water, and the washed cell pellets were immediately frozen at -20°C. For the extraction of soluble amino acids, the pellets were resuspended in MS-grade water, boiled for 15 min at 100°C saving the supernatant (soluble phase).

The soluble phase samples were analyzed using UPLC-HRMS with Dionex UltiMate 3000 liquid chromatography system (UPLC) coupled with a Q-Exactive mass spectrometer (HRMS) and interconnected with a heated electrospray ionization source (H-ESI) (Thermo Fisher Scientific, Sunnyvale, CA, USA). Separation of the analytes was performed on an Acquity Premier BEH Amide VanGuard Fit Column (1.7 µm, 2.1 mm × 100 mm; Waters, Ireland). For analysis, the column was kept at 50°C; the injection volume was kept at 1 µl and the flow rate was 0.2 ml min$^{-1}$ for a total run time of 15 min. Mobile phase A consisted of 3% acetonitrile and mobile phase B of 90% acetonitrile, both buffered with 10 mM ammonium acetate. Separation of individual compounds was achieved using a multistep gradient of A and B, where B composition started with 85% and reduced to 80% in 2 min and further decreased to 50% over 8 min before being brought to 40% over 1 min for washout. For equilibration, the B concentration was turned back to 85% over 4 min.

Ions were monitored in positive targeted single ion monitoring (t-SIM) mode with a resolution of 70,000 at m/z = 200 and an isolation window of 17 m/z, using an inclusion parameter list determined using a water-based standard mixture containing 0.5 mM of L-lysine. Other MS parameters were spray voltage 3.5 kV, sheath gas flow rates 48 units, auxiliary gas flow rate 11 units, sweep gas flow rates 2 units, capillary temperature 256°C;, auxiliary gas heater temperature 413°C;, stacked-ring ion guide (S-lens) radio frequency (RF) level 30, automatic gain control (AGC) 2×105 ions, and maximum injection time 200 ms. Exact mass acquisition and quantification were carried out using the Thermo XCalibur Quan Browser software 4.0.27.42, with 10 ppm mass tolerance. Before and after each analysis series, six-point dilutions of equimolar $[^{13}C_6,^{15}N_2]/[^{13}C_6,^{15}N_2]$-L-lysine mix prepared in MS-grade water were injected and analysed, covering the concentration range from 0.0005 to 0.5 mM and used to determine the relative molar signal weights $w_L$ and $w_H$ of the light and heavy isotopomers.

The fractional content (labeling) of light lysine was determined based on the integrated peak intensities $I_L$ and $I_H$ corresponding to heavy and light lysine as:

$$f(t) = \frac{w_L \cdot I_L}{w_L \cdot I_L + w_H \cdot I_H} \tag{S158}$$

#### S2.7 Bioinformatic analysis

**Differential protein abundance analysis.** For a comparison of protein abundances between 30°C and 37°C conditions, the relative abundance of each protein in each sample was determined by summing the intensity of the precursors in the heavy and light channels and normalizing it for the total protein abundance in the sample. Genes with $\geq 2$ samples per temperature were tested for significant differences in protein abundance using a two-sided Welch's $t$-test on log2-transformed relative abundance values. The magnitude of the effect was defined as the mean difference in $\log_2$ relative abundance ($30°\text{C} - 37°\text{C}$).

**Comparison of protein ages decay rates between temperatures.** To account for growth rate differences between temperatures, mean ages were expressed in units of maximum mean age, $\bar{\mathcal{A}}_{max} = \mu^{-1}$, multiplying the calculated mean age values by the respective culture growth rate $\mu$, while the expected decay rate values $\bar{\kappa}$ were expressed in units of growth rate by normalizing for the growth rate.

**Gene Ontology (GO) enrichment analyses.**

For the analysis of temperature-specific differences, only proteins with paired measurements at 30° C and 37° C were included. Gene names were annotated with GO terms, namespaces, and term sizes using the g:Profiler API (gprofiler-official Python package). To compare temperature-specific age differences across GO categories, only terms with $\geq 5$ paired genes were tested, and very broad categories (term size >200) were excluded. Paired comparisons were performed using a paired t-test with Benjamini–Hochberg FDR correction; terms with $q < 0.05$ were considered significant. To reduce redundancy and highlight distinct biological processes, we applied Jaccard-based pruning (overlap $> 0.3$), retaining the representative term with the largest coverage of query genes.

For GO enrichment analysis of proteins better described by a 2-pool CM, input gene lists were defined by filtering for proteins with a minimum pool weight $\geq 0.05$. GO enrichment was performed with g:Profiler restricted to S. cerevisiae annotations and the three namespaces (BP, MF, CC). Terms with a very large size ($> 150$) or zero overlaps were excluded. The results were stored as tables and for visualization, only terms with raw enrichment $p \leq 0.001$ were retained. The top 10 terms per condition were plotted as bubble charts, where bubble size reflects the intersection size and the x position represents the fraction of query genes captured (intersection/term size, %).

#### S2.8 Python package for simulating and fitting CMs to experimental data

As part of this study, we developed and published an open-source Python package (`https://gitlab.com/elad.noor/symbolic-compartmental-model`) based on SymPy Meurer et al. [2017] for simulating and fitting CMs. Some of the features include constructing a CM from the **M**-matrix (i.e., the contributed turnovers) and observed pool weights, reducing a model by removing a set of states, fitting a given set of free parameters based on experimental labeling data, and simulating the labeling curve. In addition, the package provides a long list of functions for calculating the dynamic parameters such as the distributions of ages, residence times, and decay rates – either numerically or symbolically.

#### S2.9 Quantification of dynamic parameters of yeast proteins at different growth temperatures

##### S2.9.1 CM for fitting metabolic ages of lysine

For the estimation of the metabolic ages of lysine in protein pools (Figure 10A), we used a two-state metabolic chain CM in which the unobserved lysine pool (Lys) feeds an observed protein pool (P):

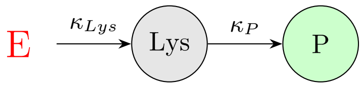


$$\mathbf{M} = \begin{bmatrix} -\kappa_{Lys} & 0 \\ \kappa_P & -\kappa_P \end{bmatrix} \tag{S159}$$

The free parameters of the CM – i.e. the contributed turnovers $\kappa_{Lys}$ and $\kappa_P$ – were fitted using the labeling dynamics of the respective proteins, where the lower bounds of both parameters were set as the growth rate to ensure that the **M**-matrix satisfies the mass balance constraint (see Section 1.7):

$$\mu \leq \kappa_{Lys}, \kappa_P \leq 100 \cdot \mu\,.$$

The vector of the observed pool weighs $(w_{Lys}, w_P)$ was set to $(0, 1)$, based on the assumption that the second pool (protein) is the only observed pool.

To determine which proteins will be used as reference proteins, the metabolic age of lysine for each one was quantified from the fitted CM and the 50 proteins with the highest metabolic age of lysine were selected. To reduce the contribution of measurement noise, we used only 1000 most abundant proteins for the selection and removed from the set of 50 the proteins with ΔAIC of the fit below -50 (i.e. bad fits).

The CMs used to determine the labeling delay (Figure 10B) were constructed as follows.

**Direct input** To model single-exponential decay, we used a 1-state CM with an unconstrained free parameter $k_P$:

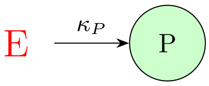


$$\mathbf{M} = \begin{bmatrix} -\kappa_P \end{bmatrix}$$

Notably, this is more of a heuristic rather than a physical model since its **M**-matrix may not always satisfy the mass balance constraint.

**1-state lysine source** The "1-state lysine source" CM was constructed and bounded as the one used to estimate the metabolic age of lysine (see model S159) except that the contributed turnover for the protein pool $k_P$ was fixed to the cell culture growth rate $\mu$:

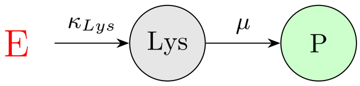


$$\mathbf{M} = \begin{bmatrix} -\kappa_{Lys} & 0 \\ \mu & -\mu \end{bmatrix}$$

The free parameter of this CM $\kappa_{Lys}$ was fitted using the median labeling values of the reference proteins, and the labeling dynamics of the lysine source $f^{\circ}(t)$ was then calculated from the fitted CM by resetting the vector of observed pool weights $(w_{Lys}, w_P)$ to $(1, 0)$.

**2-state lysine source** For construction of the "2-state lysine source" CM we used the same principles as for the "1-state lysine source" model except that the lysine source was split into two parallel pools that contributed to the growth dilution $\mu$ of the reference protein pool according to their relative weights $w_{L_0} = q_{Lys}$ and $w_{L_1} = 1 - q_{Lys}$:

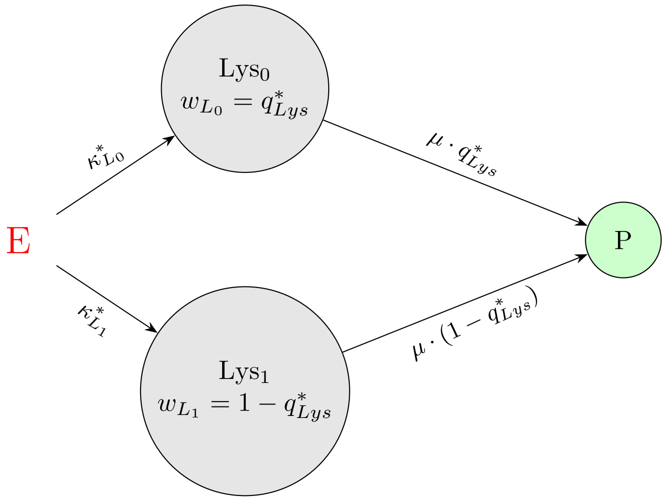


$$\mathbf{M} = \begin{bmatrix} -\kappa_{L_0} & 0 & 0 \\ 0 & -\kappa_{L_1} & 0 \\ q_{Lys} \cdot \mu & (1 - q_{Lys}) \cdot \mu & -\mu \end{bmatrix} \tag{S160}$$

To satisfy the mass-balance constraint, the free parameters of the "2-state lysine source" CM were bounded as:

$$\begin{aligned} \mu \leq \kappa_{L_0}, \kappa_{L_1} &\leq 100 \cdot \mu \\ 0 \leq \qquad q_{Lys} &\leq 1 \end{aligned}$$

For fitting of this CM, the vector of the observed pool weights was set to $(0, 0, 1)$ as only the protein pool is observed. The dynamics of lysine source labeling and its mean age (Figure 10B) were determined by resetting the observed pool weights in the fitted CM to $(q_{Lys}, 1 - q_{Lys}, 0)$ according to the relative weights of the two lysine pools.

##### S2.9.2 CMs for fitting protein dynamic parameters

In determining the dynamic parameters of the proteins, we have used CMs of 1, 2 or 3 pools to represent the proteins themselves that were fed by the known two-pool lysine source (see model S160).

**1 protein pool** Specifically, for the one-pool option the CM was constructed as the latter model except that the parameters of the lysine source denoted $(\kappa^*_{L_0}, \kappa^*_{L_1}, q^*_{Lys})$ were fixed to those quantified from the lysine input fitting (see model S160) while the contributed turnover of the protein pool $k_P$ was kept as a free parameter bounded by the mass-balance constraint:

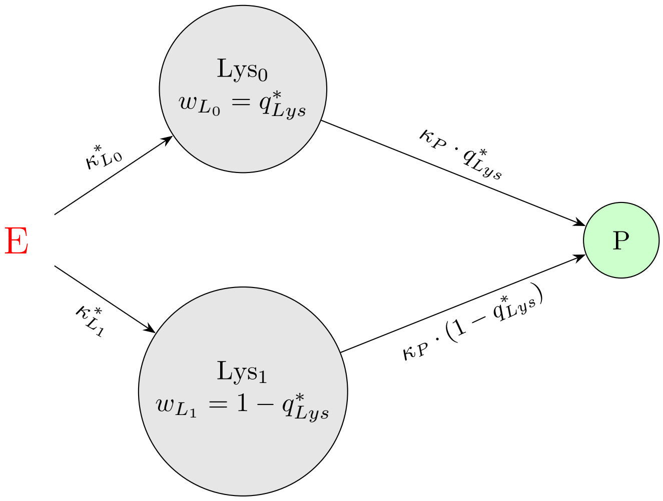


$$\mathbf{M} = \begin{bmatrix} -\kappa^*_{L_0} & 0 & 0 \\ 0 & -\kappa^*_{L_1} & 0 \\ q^*_{Lys} \cdot \kappa_P & (1 - q^*_{Lys}) \cdot \kappa_P & -\kappa_P \end{bmatrix} \tag{S161}$$

$$\mu \leq \kappa_{\kappa_P} \leq 100 \cdot \mu$$

The fitted model was then reduced to retain only the protein pool $S_2(P)$, and the reduced fit was then used to quantify various dynamic parameters of the respective proteins.

**2 protein pools** The two-pool analogue of the single-pool model was constructed analogously except that the lysine source in this case fed two parallel protein pools with their own contributed turnovers $(\kappa_{P1}, \kappa_{P2})$ and weights $(q_P, 1 - q_P)$ as free parameters satisfying the mass balance constraints:

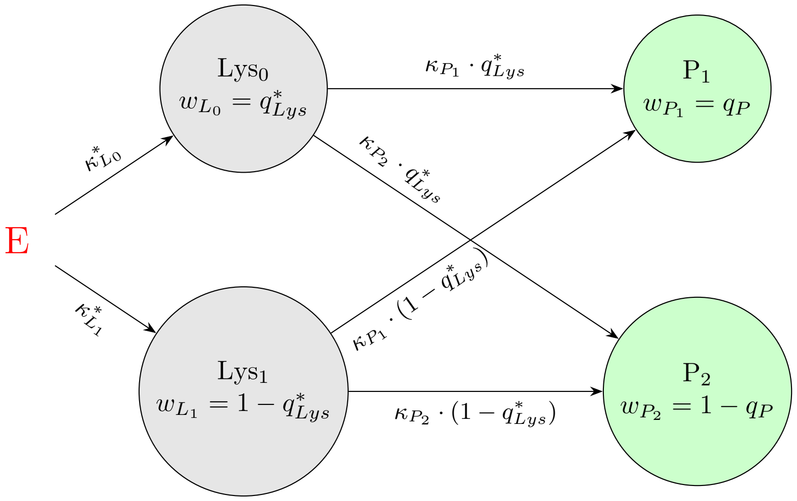


$$\mathbf{M} = \begin{bmatrix} -\kappa^*_{L_0} & 0 & 0 & 0 \\ 0 & -\kappa^*_{L_1} & 0 & 0 \\ \kappa_{P_1} \cdot q^*_{Lys} & \kappa_{P_1} \cdot (1 - q^*_{Lys}) & -\kappa_{P_1} & 0 \\ \kappa_{P_2} \cdot q^*_{Lys} & \kappa_{P_2} \cdot (1 - q^*_{Lys}) & 0 & -\kappa_{P_2} \end{bmatrix} \tag{S162}$$

$$\begin{aligned} \mu \leq \kappa_{P_1}, \kappa_{P_2} &\leq 100 \cdot \mu \\ 0 \leq \quad q_P &\leq 1 \end{aligned}$$

In this case, the vector of the observed pool weights used for the fitting was set to $(0, 0, q_P, 1 - q_P)$ corresponding to the observed two-pool subsystem representing the protein.

As in the previous model, the quantification of the dynamic parameters of the respective proteins was performed by reducing the fitted model to retain only the protein subsystem. To determine the dynamic parameters of the individual protein pools, the full model was reduced to retain only this pool, and its observed pool weight was reset to 1.

**3 protein pools** The three-pool was constructed similarly to the 2-pool one, with contributed turnovers $(\kappa_{P_1}, \kappa_{P_2}, \kappa_{P_3})$ and weights $(q_{P_1}, (1 - q_{P_1})q_{P_2}, (1 - q_{P_1})(1 - q_{P_2}))$ as free parameters satisfying the mass balance constraints:

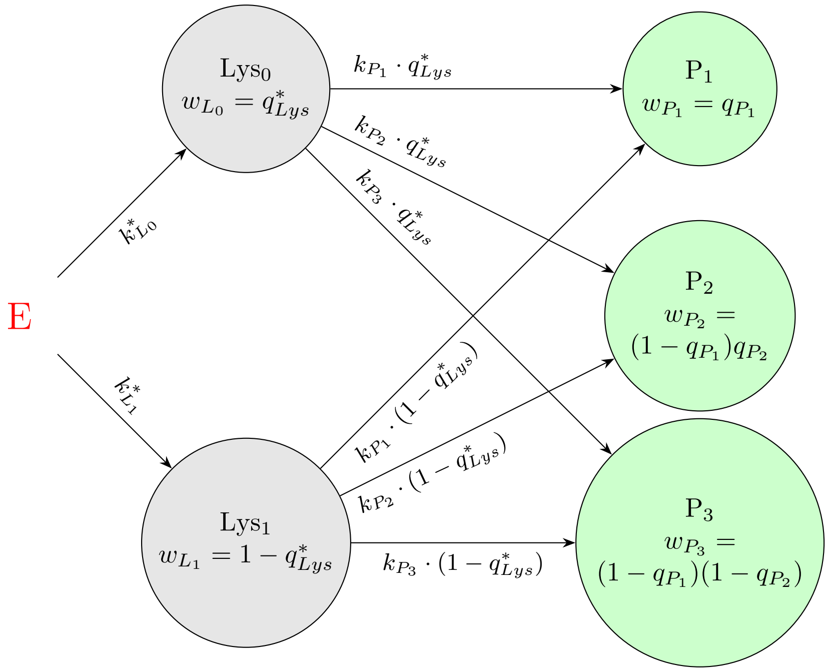

$$\mathbf{M} = \begin{bmatrix} -\kappa^*_{L_0} & 0 & 0 & 0 & 0 \\ 0 & -\kappa^*_{L_1} & 0 & 0 & 0 \\ \kappa_{P_1} \cdot q^*_{Lys} & \kappa_{P_1} \cdot (1 - q^*_{Lys}) & -\kappa_{P_1} & 0 & 0 \\ \kappa_{P_2} \cdot q^*_{Lys} & \kappa_{P_2} \cdot (1 - q^*_{Lys}) & 0 & -\kappa_{P_2} & 0 \\ \kappa_{P_3} \cdot q^*_{Lys} & \kappa_{P_3} \cdot (1 - q^*_{Lys}) & 0 & 0 & -\kappa_{P_3} \end{bmatrix} \tag{S163}$$

$$\begin{aligned} \mu \leq \kappa_{P_1}, \kappa_{P_2}, \kappa_{P_3} &\leq 100 \cdot \mu \\ 0 \leq \quad q_{P_1}, q_{P_2} &\leq 1 \end{aligned}$$

In this case, the vector of the observed pool weights used for the fitting was set to $(0, 0, q_{P_1}, (1-q_{P_1})q_{P_2}, (1-q_{P_1})(1-q_{P_2}))$ corresponding to the observed three-pool subsystem representing the protein.

As in the previous model, the quantification of the dynamic parameters of the respective proteins was performed by reducing the fitted model to retain only the protein subsystem. To determine the dynamic parameters of the individual protein pools, the full model was reduced to retain only this pool, and its observed pool weight was reset to 1.

##### S2.9.3 Data Availability

The raw data, scripts, and age scores themselves can be found at `https://gitlab.com/elad.noor/karmage`.

#### S2.10 Age score of NPC bait-prey pairs and minimum maturation time estimates

In Onischenko et al. [2020], we collected dynamic labeling values from 3 time points for 320 bait-prey pairs of affinity-isolated complexes of yeast NPC proteins (10 baits, 32 preys). As a way to integrate all the labeling measurements into a single high-confidence value that reflects the overall labeling speed and can be used to reliably determine the assembly order, we defined the *age score* – the AUC of the fitted curve in the first 90 minutes after the metabolic pulse (Supplemental Figure S14A and C). Since the half-life of the labeling for all measured pairs was longer than 90 minutes, using the age score (rather than the mean age) was a reasonable compromise, as it is less sensitive to the shape of the labeling curve at long, unmeasured timescales. Notably, the age score is analogous to the age-cohort mean value of the metabolic age (see Appendix S1.3.6) which accounts only for the lysine which is younger than the last measurement time point.

In more detail, to calculate the age score for each bait-prey pair, we fitted the experimental values once using a 3-state model and once using a 4-state model in which a uniformly decaying chain of variable length (see Appendix S1.7.7) representing maturation intermediates of the prey in the growing culture was fed by a free lysine pool. In both cases, the first state represented the free lysine pool and its contributed turnover was a fixed value (set to $\mu/\psi_{\text{Lys}}$, where $\psi_{\text{Lys}} = 0.35$) which was the size determined in the previous study Onischenko et al. [2020]. The other states represent the precursor, observed, and (optionally) inaccessible pools.

**3-state model**

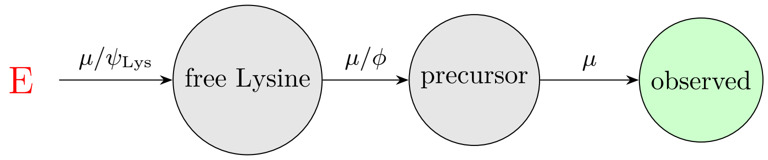


$$\mathbf{M} = \begin{bmatrix} -\mu/0.35 & 0 & 0 \\ \mu/\phi & -\mu/\phi & 0 \\ 0 & \mu & -\mu \end{bmatrix}$$

**4-state model**

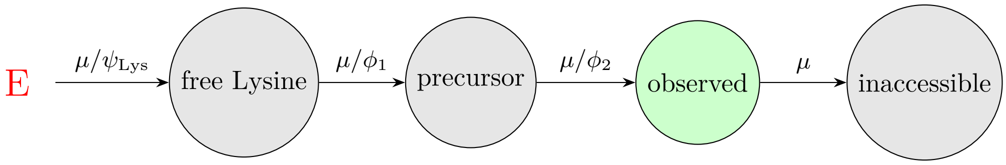


$$\mathbf{M} = \begin{bmatrix} -\mu/0.35 & 0 & 0 & 0 \\ \mu/\phi_1 & -\mu/\phi_1 & 0 & 0 \\ 0 & \mu/\phi_2 & -\mu/\phi_2 & 0 \\ 0 & 0 & \mu & -\mu \end{bmatrix}$$

After fitting, the model with the better AIC was chosen and the age score was calculated using the analytical expression for the AUC between 0 and 90 minutes.

For quantification of the minimum maturation times required for the co-assembly of prey NPC proteins with the Mlp1 bait (Figure S8D), the mean metabolic age of lysine in each prey co-isolated with Mlp1 was quantified as the full AUC of prey labeling as described above, subtracting the mean age of lysine input and the maximum possible reduced mean age of prey in the Mlp1 bound fraction ($\psi_{\mathrm{Lys}}/\mu$, and $\mu^{-1}$, respectively here $\psi_{\mathrm{Lys}} = 0.35$).

The raw data, scripts, and age scores themselves can be found at https://gitlab.com/elad.noor/karmage.

#### S2.11 Ranking NPC assembly models based on similarity to age scores

In order to assess different assembly models (whole hierarchies of co-assembly events) for agreement with experimental data, we designed a scoring system based on the similarity between the experimentally observed ordering inferred from the age scores and the ordering of preys in the model. For each prey, the ordering from the model was derived from the sequence of nodes in the hierarchy connecting the root (full complex) and the leaf representing the prey. The similarity was calculated using Spearman rank correlations, once for each prey, and then averaged into a single score (Supplemental Figure S14B).

Overall, we had 9 preys/baits in the dataset, which meant that there are around 2 million possible assembly orders (represented by binary trees). Iterating through all of them and calculating the Spearman correlations took around 20 minutes on a workstation utilizing 30 CPU (Intel Core i9-10980XE) and about 20 GB of RAM in total.

As a measure of significance, we generated a random assembly model and compared the Spearman scores of the age scores versus a matrix of randomly permuted scores. We repeated this 1 million times and compared the distributions of the two scores (“real” versus “random”) using a histogram plot (Supplemental Figure S14D). The random scores distribute around a mean of 0, while the real ones have a mean of about 0.25. Also the best tree fitted on the randomized data scored roughly 30% lower than the best tree fitted on real data. This indicated to us that the age scores calculated for the bait-prey pairs significantly match a non-random assembly order.

The scripts, tree scores and bootstrapping results can all be found at https://gitlab.com/elad.noor/karmage.

#### S2.12 Appendix: effects of delayed labeling and age-dependent decay on the quantification of dynamic parameters

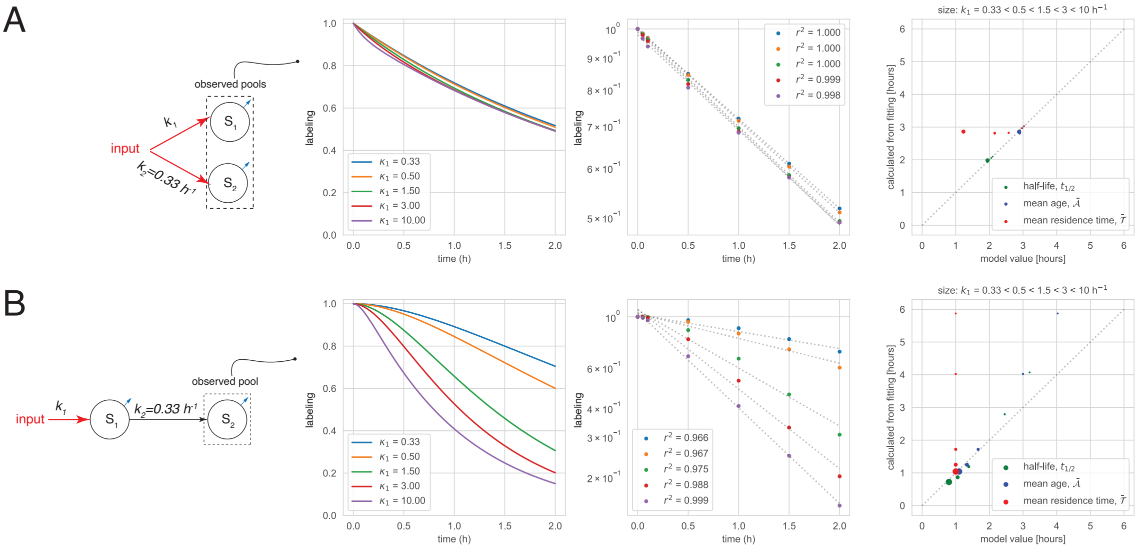


Figure S10: **Effects of delayed input and complex decay patterns on the quantification of dynamic parameters by semi-log fitting.** (A) The effect of age-dependent decay simulated with a two-pool model and a variable turnover of one of the pools. (B) The effect of delayed input simulated by a varying contributed turnover of the unobserved pool $S_1$. Note that despite acceptable goodness of fit the dynamic parameter evaluations based on the linear regression and log-transformed labeling values (semi-log fitting) can significantly deviate from the actual dynamic parameters.

#### S2.13 Appendix: quantifying dynamics of yeast proteome in mild heat stress

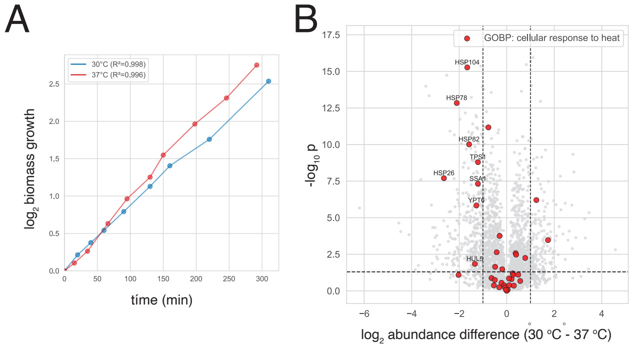


Figure S11: **Growth dynamics and protein abundances of yeast cultures in dynamic SILAC assays at 30°C and 37°C.** (A) Semi-log plot of biomass growth based on culture turbidity measurements. Note linear dynamics at both temperatures ($R^2 > 0.995$), indicating balanced exponential growth. (B) Differences in relative abundance of yeast proteins at 30°C vs 37°C were determined based on summed (heavy + light) peptide intensities across all labeling time points. Significance thresholds (dashed lines) were set at $p < 0.05$ and $|\log_2 \text{abundance ratio}| \geq 1$. Heat-response genes (GO: cellular response to heat) were highlighted in red, with significant hits labeled. Note that heat shock proteins (red, labeled) have significantly higher abundances at 37°C. (C) Mean relative abundance of reference proteins. Note that the abundance of reference proteins remains relatively stable throughout the labeling time course.

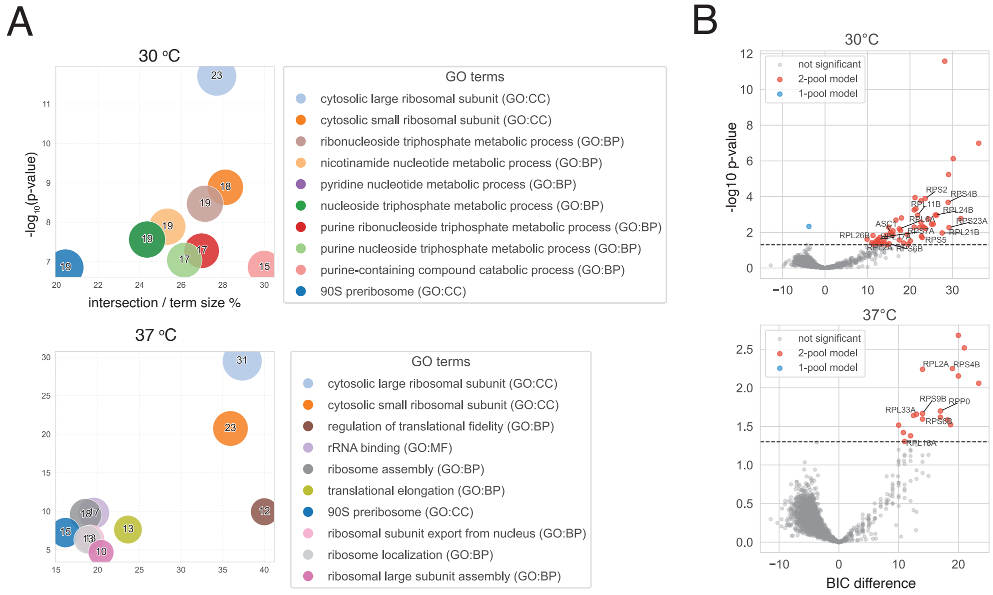


Figure S12: **Complex kinetic pool structure of ribosomal proteins.** (A) GO enrichment analysis of proteins whose labeling dynamics in better described by a 2-pool CM (Figure 11A). Note that at both temperatures ribosomal subunits are enriched at the 2-pool category. (B) Certainty of 2-pool model description evaluated using bootstrapping. Mean and SD values of BIC for the fits with 1-pool and 2-pool models were evaluated by resampling the labeling data with repetition. The p-values that BIC is the same for both models were calculated considering them random normally distributed independent variables with the corresponding mean and SD. Note that low p-values indicate confident preference for the respective model.

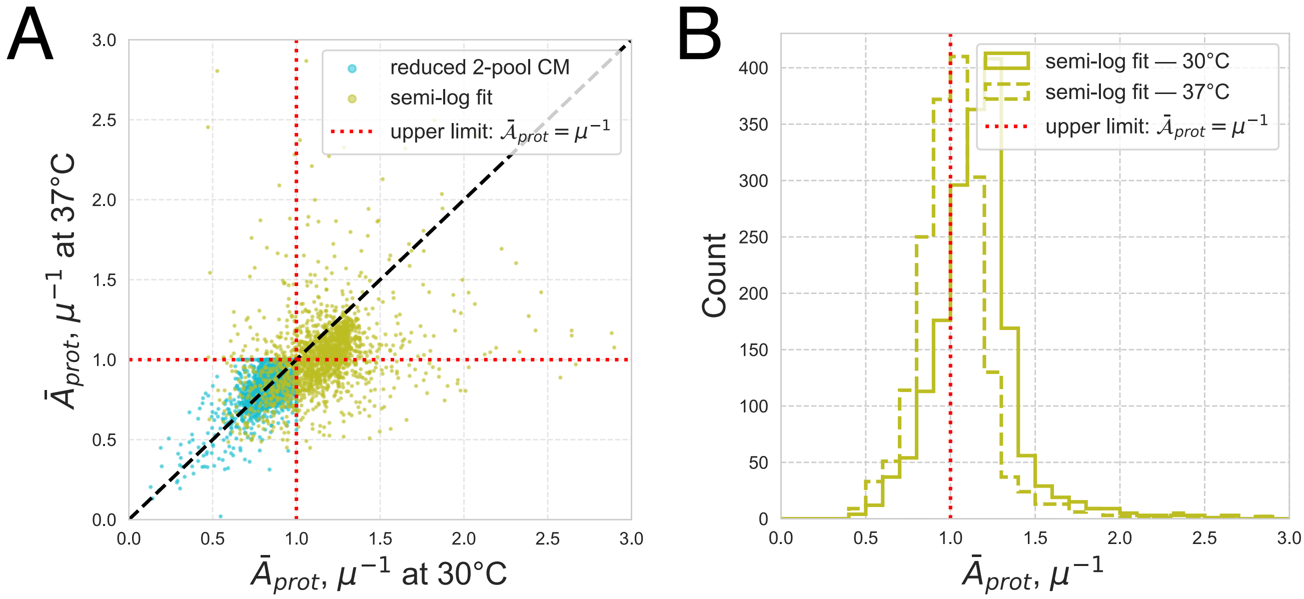


Figure S13: **Mean metabolic ages of yeast proteins quantified using alternative models.** (A) Scatter plot showing comparison of mean metabolic ages of yeast proteins quantified at 30°$C$ and 37°$C$ using reduced 2-pool CM (see Figure 12) versus a classical method relying on the direct fitting of log-transformed labeling dynamics with linear regression Schwanhäusser et al. [2011] to calculate metabolic ages based on the labeling curve slope.(B) Histogram representation of mean metabolic age distribution obtained using semi-log fitting. In both cases metabolic ages are expressed in units of inverse growth rate $\mu^{-1}$ equivalent to the maximum possible mean age of a protein in a growing system (red dotted line). Note that semi-log fitting produces a systematic bias rendering mean metabolic ages of the majority of proteins significantly exceed this limit.

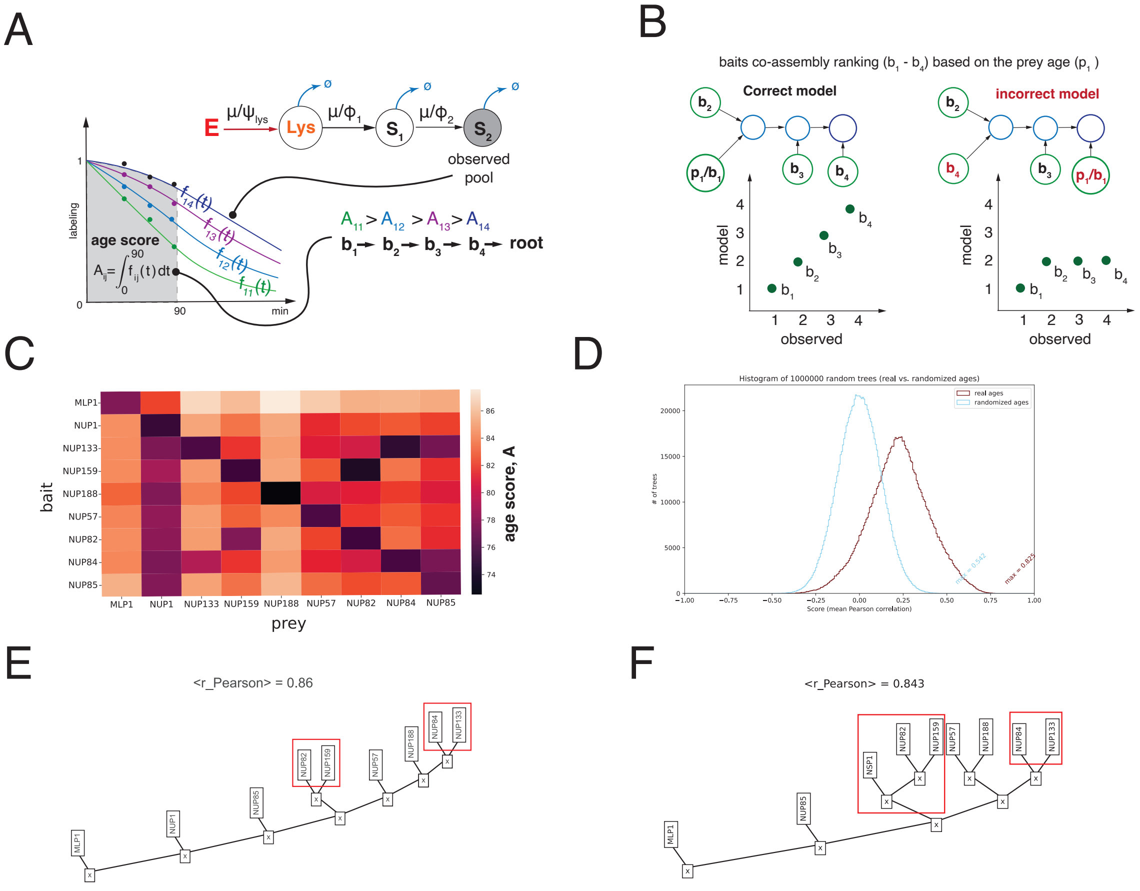


Figure S14: **Determining the order of NPC assembly using dynamic labeling.** (A) Computing bait-prey assembly order based on the dynamic labeling. The labeling readouts obtained with each bait for a given prey were interpolated with a 3- or 4-step CM and the fit was used to compute the age score defined as the AUC in the experiment time interval [0, 90 min]. The age score was in turn used to compare the hight of prey labeling curves obtained with different baits and to determine bait assembly order with each prey. (B) Example of correlation relationships between observed and model-predicted assembly order of baits with a given prey. Note that incorrect models would produce poorer correlations. (C) Age scores calculated for 9 baits co-isolating each other. (D) Comparison of the model scores based on real labeling data (real age scores) an based on the randomly permuted age scores. Note that randomized data do not provide nearly as good agreement with any assembly model as the real data. (E and F) Best-scoring assembly models produced using nonparametric quantification of the age scores for the same bait-prey pairs as above (E) or when one of the baits (Nup1) was replaced with Nsp1 that does not have a single assembly path (Nsp1 is a component of two subcomplexes, Nup82-Nup159-Nsp1 and Nsp1-Nup57-Nup49) (F). Red boxes denote directly interacting proteins. Note that using a nonparametric model or changing selection of baits do not perturb the overall predicted assembly order.

Aaron Meurer, Christopher P. Smith, Mateusz Paprocki, Ondřej Čertík, Sergey B. Kirpichev, Matthew Rocklin, AMiT Kumar, Sergiu Ivanov, Jason K. Moore, Sartaj Singh, Thilina Rathnayake, Sean Vig, Brian E. Granger, Richard P. Muller, Francesco Bonazzi, Harsh Gupta, Shivam Vats, Fredrik Johansson, Fabian Pedregosa, Matthew J. Curry, Andy R. Terrel, Štěpán Roučka, Ashutosh Saboo, Isuru Fernando, Sumith Kulal, Robert Cimrman, and Anthony Scopatz. SymPy: symbolic computing in Python. *PeerJ Computer Science*, 3:e103, January 2017. ISSN 2376-5992. doi: 10.7717/peerj-cs.103. URL https://peerj.com/articles/cs-103. Publisher: PeerJ Inc.

Björn Schwanhäusser, Dorothea Busse, Na Li, Gunnar Dittmar, Johannes Schuchhardt, Jana Wolf, Wei Chen, and Matthias Selbach. Global quantification of mammalian gene expression control. *Nature*, 473(7347):337–342, May 2011. ISSN 0028-0836, 1476-4687. doi: 10.1038/nature10098. URL https://www.nature.com/articles/nature10098.
